# Dynamic proximity interaction profiling suggests that YPEL2 is involved in cellular stress surveillance

**DOI:** 10.1101/2023.07.31.551286

**Authors:** Gizem Turan, Çağla Ece Olgun, Hazal Ayten, Pelin Toker, Annageldi Ashyralyyev, Büşra Savaş, Ezgi Karaca, Mesut Muyan

**Author notes:** Equal contribution, should be considered as the first author.

## Abstract

YPEL2 is a member of the evolutionarily conserved YPEL family involved in cellular proliferation, mobility, differentiation as well as senescence and death. However, the mechanism by which YPEL2, or YPEL proteins, mediates its effects is yet unknown. Proteins perform their functions in a network of proteins whose identities, amounts, and compositions change spatiotemporally in a lineage-specific manner in response to internal and external stimuli. We here explored interaction partners of YPEL2 by using dynamic TurboID-coupled mass spectrometry analyses to infer a function for the protein. Our results using inducible transgene expressions in COS7 cells indicate that proximity interaction partners of YPEL2 are largely involved in RNA and mRNA metabolic processes, ribonucleoprotein complex biogenesis, regulation of gene silencing by miRNA, and cellular responses to stress. We showed that YPEL2 interacts with RNA binding protein ELAVL1 and selective autophagy receptor SQSTM1. We also found that YPEL2 participates in events associated with the formation/disassembly of stress granules in response to sodium arsenite an oxidative stress inducer. Establishing a point of departure in the delineation of structural/functional features of YPEL2, our results suggest that YPEL2 may be involved in stress surveillance mechanisms.

## INTRODUCTION

17β-Estradiol (E2), the most potent estrogen hormone in the circulation, is involved in a wide variety of vital physiological functions ranging from the development and maintenance of reproductive organs to the regulation of cardiovascular, musculoskeletal, immune, and central nervous system homeostasis^1–3^. Effects of E2 on cells are primarily mediated by estrogen receptors (ER) α and β, which are the products of distinct genes and act as transcription factors^1–3^. Upon binding to E2, the activated ER modulates the expression of E2 target genes, whose protein products are involved in the regulation of cellular proliferation, differentiation, and death^1–3^. Despite the contribution of both inherited and environmental factors, perturbations in E2 signaling are considered to be the major factor for the initiation and development of estrogen-target tissue malignancies, including breast cancer^1–3^. Due to the central role of the E2-ERα signaling in the physiology and pathophysiology of estrogen target tissues, identifying estrogen-responsive genes and functional characterization of their protein products involved in the manifestation of cellular phenotype could provide new prognostic tools and/or therapeutic targets. In an effort directed at deciphering the underlying mechanism of E2-ERα actions at genomic levels, we previously identified *Yippee Like 2* (*YPEL2*) as an E2-ER responsive gene^4^.

*YPEL2* is a member of the highly conserved YPEL gene family named after the *Drosophila* Yippee protein. The *YPEL* family has 100 genes from 68 species ranging from yeast, *C. elegans*, flies, and plants to mammals^5, 6^. Although the number of YPEL genes varies among eukaryotes, the human *YPEL* family, like other mammals, includes five distinct YPEL genes: *YPEL1, 2, 3, 4,* and *5* located in different chromosomes^5, 6^. The YPEL family genes encode small proteins with molecular masses (MMs) ranging from 13,5 to 17,5 kDa that show remarkably high amino-acid sequence identities (43.8-96.6%). Amino-acid sequence alignments based on the highly conserved cysteine residues of all identified YPEL proteins in different species revealed two cysteine pairs that are 52 amino acids apart (C-X_2_-C-X_52_-C-X_2_-C) predicted to form a zinc-finger like metal binding pocket, or the Yippee domain^5, 7^. This high degree of evolutionary conservation among YPEL proteins implies that YPELs are critically involved in various cellular processes. Indeed, experimental studies suggest that the YPEL proteins, including YPEL2, participate in cellular proliferation, mitochondrial function, morphology, mobility, differentiation as well as senescence and death^6, 8–27^. Moreover, deregulated expression of YPEL genes could be associated with, or contribute to, the initiation/development of various disorders, malignancies, and resistance to therapies^11, 13, 19, 24, 28–32^. Similarly, high levels of YPEL2 expression are suggested to be one factor contributing to Dominant Retinitis Pigmentosa^33^ and autophagy processes in breast cancer^34^.

However, the mechanisms by which YPEL proteins mediate their effects are yet unclear. Proteins perform their functions in a network of proteins whose identities, amounts, and compositions change spatiotemporally in a lineage-specific manner in response to various internal and external stimuli^35, 36^. To better understand the functional features of YPEL2 in mediating cellular processes in both physiology and pathophysiology, we initially explored the identification of protein interaction networks of YPEL2 dynamically. Using inducible transgene expression in COS7 cells followed by time-resolved TurboID-coupled mass spectrometry analyses, we found that YPEL2 proximity interacts with proteins largely involved in RNA and mRNA metabolic processes, ribonucleoprotein complex biogenesis, regulation of gene silencing by miRNA, and cellular responses to stress. We also found that YPEL2, as endogenous YPEL proteins in COS7 cells, participates in events associated with the formation/disassembly of stress granules in response to sodium arsenite an oxidative stress inducer. Our results suggest that YPEL2 is a critical component in cellular responses to oxidative stress.

### Experimental Procedures

#### In silico analyses

The alignment of amino acid sequences of human YPEL proteins was generated with the Jalview^37^ program (https://www.jalview.org/) using the ClustalOmega^38^ plug-in (https://www.ebi.ac.uk/Tools/msa/clustalo/). For the tertiary structure prediction and superimposition of the tertiary structures of YPEL1-5 proteins, we used the AlphaFold^40, 41^ server (https://alphafold.ebi.ac.uk/) with the ChimeraX molecular visualization^42, 43^ program (https://www.cgl.ucsf.edu/chimerax/). For the homology modeling of YPEL proteins, we used the Phyre2^44^ protein fold recognition server (http://www.sbg.bio.ic.ac.uk/~phyre2/html/page.cgi?id=index). We also used a Venn diagram tool (https://bioinformatics.psb.ugent.be/webtools/Venn/) to visualize the intersections of proximity interaction partners. Tissue-specific gene expression analyses were carried out with the GTEx portal (https://gtexportal.org/home/). For chromatin features of transcription start sites, we used the Cistrome (http://cistrome.org/) database^45^. Functional features, interactome profiles, and membership analyses of putative interacting proteins of YPEL2 were carried out with the Metascape^46^ portal (https://metascape.org/).

### Biochemicals

Restriction and DNA modifying enzymes were obtained from New England Bio-Labs (Beverly, MA, USA) or ThermoFisher (ThermoFisher, Waltham, MA, USA). Chemicals were obtained from Sigma-Aldrich (Germany), ThermoFisher, or BioTechne-Tocris Corp. (Minnesota, USA). DNA and RNA isolation kits were purchased from ZymoResearch (California, USA). SYBR Green Supermix kit was procured from Bio-Rad Life Sciences Inc. (California, USA). Pageruler Prestained Protein Ladder (ThermoFisher, 26616) or Pageruler Plus Prestained Protein Ladder (ThermoFisher; 26620) was used as the molecular mass marker. Sodium Arsenite (SA, NaAsO_2_, S7400) was acquired from Sigma-Aldrich Inc. (Missouri, USA).

Monoclonal mouse Flag-M2 antibody (F1804) was purchased from Sigma-Aldrich. Rabbit polyclonal HDAC1 (ab19845), rabbit polyclonal β-actin (ab8227), rabbit polyclonal biotin (ab53494) antibodies, and Alexa Fluor 647-conjugated goat anti-rabbit IgG (ab150083) were acquired from Abcam Inc. (Connecticut, USA). Rabbit polyclonal pan-YPEL (sc-99727), mouse monoclonal G3BP1 (sc-365338), mouse monoclonal ELAVL1 (sc-5261), mouse monoclonal Vimentin (sc-6260), mouse monoclonal γ-tubulin (sc-17787), mouse monoclonal SQSTM1 (sc-48402) antibodies together with a siRNA poll specific for ELAVL1 transcripts (sc-35619) and HRP-conjugated goat anti-mouse IgG (sc-2005) were purchased from Santa Cruz Biotechnology, Inc. (SCBT, Texas, USA). A rabbit polyclonal antibody specific to G3BP1 (13057-2-AP) was obtained from Proteintech Group (Illinois, USA). We obtained Alexa Fluor 488-conjugated goat anti-mouse IgG (BioLegend 405319) from BioLegend Inc. (California, USA) and HRP-conjugated goat anti-rabbit IgG (Advansta R-05072-500) from Advansta Inc. (California, USA). AllStar negative control siRNA (SI03650318) was purchased from Qiagen Inc. (Germany).

### Cell lines and growth conditions

The estrogen-responsive and ERα-synthesizing MCF7 cells derived from a breast adenocarcinoma and COS7 cells derived from African green monkey kidney fibroblasts were maintained in high glucose (4.5 g/L) Dulbecco’s modified eagle medium (DMEM) without phenol red (Biochrom F0475) supplemented with 10% fetal bovine serum (FBS, Biochrom S0115), 1% penicillin-streptomycin, and 0.5% L-glutamine as we described previously^47, 48^. To test the effects of ligands on YPEL gene expressions in MCF7 cells, we used E2 (Sigma-Aldrich, E2758) and Imperial Chemical Industries (ICI) 182,780, (BioTechne-Tocris, 1047), a complete antagonist for ERα. For 10^-9^ M E2 and/or 10^-7^ M ICI treatments, MCF7 cells were grown in DMEM supplemented with charcoal-dextran-stripped FBS (CD-FBS) to reduce endogenous steroid hormone content. For the induction of transgene expression by doxycycline (Dox), COS7 cells were maintained and transfected in the DMEM medium containing 10% tetracycline-free FBS (Tet-free FBS, Biowest, France, S181T) to minimize the effect of endogenous tetracycline on transgene expression. Media of cultured cells incubated in a humidified incubator with 5% CO_2_ at 37°C were refreshed every three days. Cells were passaged for a maximum of eight passages.

### RNA isolation and RT-qPCR

Total RNA isolation was carried out with QuickRNA Miniprep Kit (ZymoResearch) according to the manufacturer’s instructions including on-column DNase I digestion as we described^49^. Isolated total RNA from MCF7 cells treated without (ethanol control) or with 10^-9^ M E2 and/or 10^-7^ M ICI for 24 hours was used for cDNA synthesis (The RevertAid First Strand cDNA Synthesis Kit, Thermo-Fisher) with oligo (dT)_18_ primers according to manufacturer’s instructions. We also used total RNA from COS7 cells maintained under the steady-state condition for cDNA synthesis. The expression of YPEL transcripts was then assessed with RT-qPCR using transcript-specific primer sets (Supplemental Information, Table S1) and the SsoAdvanced Universal SYBR Green Supermix kit (Bio-Rad). Expression levels of transcripts were assessed with the efficiency corrected form of the 2^−ΔCT^ method and normalized using the *RPLP0* expression levels^50^. Relative expression levels of YPEL genes were assessed using the 2^−ΔΔ^CT method^50^ and normalized using the *RPLP0* expression levels. Results were arbitrarily adjusted to the expression level of YPEL1. In all RT-qPCR experiments, MIQE Guidelines were followed^51^.

### Cloning of YPEL1 through YPEL5 cDNAs

For the cloning of open reading frames of *YPEL1-YPEL5*, we used RefSeq mRNA sequences obtained from the NCBI database, which were aligned with the MUSCLE (MUltiple Sequence Comparison by Log-Expectation) software^52^. Two sets of primers using NCBI Primer Blast (https://www.ncbi.nlm.nih.gov/tools/primer-blast/) were generated for each gene based on the sequences in the NCBI database. The first set of primers consisted of a forward primer from 5’UTR and a reverse primer from 3’UTR. The second set of primers internal to the first set of primers, nested primers, contained restriction enzyme sites that were designed to provide flexibility in cloning. For YPEL3, we also used transcript variant 1, which has the identical core sequence of transcript variant 2, containing an extended 5’ sequence that encodes an additional 38 amino acids at the amino terminus. In the engineering of all constructs, the first ATG was designed to be embedded into the consensus Kozak sequence (CCATGG) for efficient translation; reverse cloning primers were also designed to contain a polyA signal (TAATAAA) that comprises a stop codon as we described previously^49, 53, 54^. The generated YPEL cDNAs were then cloned into the pcDNA 3.1(-) vector. We also cloned YPEL cDNAs into the pcDNA 3.1(-) vector bearing in-frame sequences encoding for an amino-terminally located 3xFlag (3F) tag at the 5’-end of the multiple cloning site. All constructs were sequenced for the fidelity of encoding sequences. All primer sequences are given in Supplemental Information, Table S1.

For inducible expression of the YPEL2 cDNA, we used a tetracycline-responsive expression system, pINDUCER20^55^ obtained from Addgene (Plasmid #44012; Massachusetts, USA). pINDUCER20 is a single vector system that encodes the reverse tetracycline-controlled transactivator (rtTA3) and the neomycin resistance gene as a constitutive, bicistronic transcript^55^. rtTA3 binds to and activates transgene expression from the TRE promoter, which is a seven-repeat of a 19-nucleotide long tetracycline operator (tetO), only in the presence of tetracycline or the stable tetracycline analog doxycycline, Dox. We modified pINDUCER20 by inserting a DNA fragment bearing a Multiple Cloning Site (MCS) on gateway destination sequences for easy cloning of the gene of interest (pINDUCER20-MCS). We then cloned 3F-YPEL2 cDNA into the pINDUCER20-MCS and sequenced it to ensure the sequence fidelity.

### Establishment of the TurboID System

Enzyme-catalyzed proximity labeling (PL) is an effective tool for unbiased proteomic analysis of macromolecular complexes, organelles, and protein interaction networks. Following its introduction, the proximity-dependent biotinylation (BioID) approach has been widely used, as we have^54^, for the identification of interacting partners of many proteins^56^. The BioID approach is based on the genetic fusion of a promiscuous biotin ligase, a mutant *E. coli* biotin ligase enzyme, BirA*(R118G), which is defective in both self-association and DNA binding, to a protein-of-interest to biotinylate proximity proteins^56^. The addition of biotin initiates the covalent tagging of endogenous proteins within a few nanometers of the promiscuous enzyme. Biotinylated proteins are then selectively isolated with biotin-affinity capture and identified with mass spectrometry (MS). However, one major disadvantage of BioID is its slow kinetics in biotinylation requiring hours to produce sufficient biotinylated material for proteomic analysis^57^. Studies for the dynamic proximity protein interactions necessitate the actuation of proximity biotin labeling with greater efficiency occurring on a timescale of minutes than the BioID approach. For the identification of interacting partners of YPEL2, we envisioned that the generation of time-dependent profiling of stable, transient, and/or weak protein interactions could provide important clues about the functional features of YPEL2. For this purpose, we used 3xHA-TurboID-NLS-pcDNA3^57^ (Plasmid #107171; Addgene, Massachusetts, USA). TurboID system uses a biotin ligase engineered through yeast display-based directed evolution of BirA* that catalyzes proximity biotin labeling of proteins with higher efficiencies and faster kinetics compared to BirA*. This allows the investigation of protein interaction networks dynamically^57^. The TurboID-HA cDNA was generated by PCR with a primer containing sequences encoding a carboxyl-terminally located HA tag with a stop codon present in a polyA motif (TAATAAA) and a primer encoding sequences with or without the initiation methionine codon (iMet) by the use of the 3xHA-TurboID-NLS-pcDNA3 vector as the template. In this engineering, our basic aim was to generate transgene proteins that display an overlapping intracellular distribution thereby minimizing false-negative results (Supplemental Information, Fig. S1). The TurboID-HA amplicon with the iMet codon was then cloned with appropriate restriction enzyme sites into pcDNA3.1(-) to generate pcDNA-TurboID-HA. We then cloned the 3F-YPEL2 cDNA into pcDNA-TurboID-HA to generate the pcDNA-3F-YPEL2-TurboID-HA expression vector. The constructs were sequenced for sequence fidelity.

#### Immunocytochemistry (ICC) and Western Blot (WB)

To assess the intracellular location of transgene products, we transiently transfected COS7 cells with the expression vector pcDNA3.1(-) bearing none (EV) or a cDNA for 3F-YPEL2, Turbo-HA or 3F-YPEL2-TurboID-HA. Cells after a 24h transient transfection were processed for ICC and WB analyses as we described previously^58^. In brief, for ICC, cells on coverslips were washed three times with 1x PBS and fixed with 3.7% formaldehyde for 30 min. Cells were then permeabilized with 0.4% Triton-X-100 for 10 min. Blocking was carried out with 10% Bovine Serum Albumin (BSA) for 1 h. The Flag (1:250, Sigma Aldrich, F-1804) in 3% BSA, the HA (1:500, Abcam ab9119), or the Biotin (1:500, in 3% BSA, Abcam ab53494) primary antibody in 3% BSA was added sequentially onto the cells for 2 h. Cells were then washed three times with 1x PBS and incubated with a secondary antibody for 30 min for each. An Alexa Fluor 488-conjugated goat anti-mouse (1:1000 in 3% BSA; Abcam ab150113) for the Flag antibody or an Alexa Fluor 594-conjugated goat anti-rabbit secondary antibody (1:1000 in 3% BSA; Abcam ab150080) for the HA or the Biotin antibody was used. After three 1x PBS washes, coverslips were mounted onto the glass slides with the Fluoroshield Mounting Medium containing 4′,6-diamidino-2-phenylindole (DAPI). Imaging was carried out with a Nikon Eclipse 50i Fluorescence Microscope. We observed that 3F-YPEL2-Turbo-HA as 3F-YPEL2 shows diffuse intracellular staining encompassing the nucleus and cytoplasm (Supplemental Information, Fig. S1A). For WB analysis, equal amounts (50 µg) of cellular extracts were subjected to Sodium Dodecyl Sulfate (SDS)-10% Polyacrylamide Gel Electrophoresis (PAGE), and proteins were scanned with the Flag antibody for 3F-YPEL2 and 3F-YPEL2-Turbo-HA or the HA antibody for Turbo-HA. Results indicated that proteins were synthesized at expected MMs (Supplemental Information, Fig. S1B). To assess the incorporation of biotin into the endogenous proteins, cells transfected with pcDNA-3F-YPEL2-TurboID-HA expression vector for 24 h were incubated without or with 50 µM biotin in the presence of 1 mM ATP for 3 or 16 h. Equal amounts of cellular extracts were then subjected to WB analysis using a biotin antibody (1:500 dilution, Abcam ab53494). Results indicated that 3F-YPEL2-TurboID-HA effectively induces the incorporation of biotin into the proteins (Supplemental Information, Fig. S1C).

#### Dynamic analysis of the proximity interaction network of YPEL2 with TurboID

Based on our findings that TurboID can be effectively used for dynamic protein interaction profiling, we cloned the 3F-YPEL2, 3F-YPEL2-Turbo-HA, or Turbo-HA cDNA into pINDUCER20-MCS to dictate the timing and the level of protein synthesis in transiently transfected cells to minimize potential adverse effects of transgene overexpression.

#### Assessing Dox concentrations for an effective transgene synthesis with WB

COS7 cells, 6×10^4^, seeded in six-well tissue culture plates and grown in DMEM containing 10% Tet-Free FBS for 48 h were transiently transfected with the pINDUCER20-MCS vector bearing none (empty vector, EV), 3F-YPEL2, 3F-YPEL2-Turbo-HA, or Turbo-HA cDNA. Twenty-four hours after transfection, cells were incubated in a fresh medium containing various concentrations (0-1000 ng/ml) of Dox for 24 h. Cells were then collected with trypsinization and pelleted by centrifugation. Pellets were washed twice with 1x PBS and lysed with M-PER (ThermoFisher; 78,833) containing freshly added protease and phosphatase inhibitors. Total cell protein concentrations were assessed with Bradford Protein Assay (Bio-Rad, USA, 500-0201). Equal amounts (50 µg) of proteins were subjected to SDS-10%PAGE followed by WB as described above using an antibody specific to the Flag (1:1000; Flag-M2, Sigma-Aldrich, F-1804) or the HA (1:500; Abcam ab9119) epitope followed by an HRP conjugated goat anti-mouse (for the Flag-M2) or anti-rabbit (for the HA antibody) secondary antibody (1:4000). Membranes were then subjected to the chemiluminescence substrate (WesternBright ECL substrate, Advansta) for visualization, and were imaged with the Bio-Rad Image Processor. Our results indicated that Dox at the 10 ng/ml amount is an optimal Dox concentration for the maximal level of 3F-YPEL2-Turbo-HA protein synthesis (Supplemental Information, Fig. S2A).

#### Assessing the optimal concentration and the effects of doxycycline on the incorporation of Biotin onto intracellular proteins

COS7 cells grown in DMEM containing 10% TET-Free FBS for 48 h were transfected with the pINDUCER20-MCS vector bearing the Turbo-HA or the 3F-YPEL2-Turbo-HA cDNA for 24 h. Cells were then treated with 10 ng/ml Dox for transgene synthesis for 24 h. Cells were then incubated in a medium containing 50, 100, or 250 µM biotin in the presence of 1 mM ATP for 30 min, 1 h, or 3 h. Cells were lysed and equal amounts (50 µg) of proteins were fractionated with SDS-10%PAGE for WB analysis using the biotin antibody (1:500 dilution, Abcam ab53494). Our results indicated that TurboID alone or as the 3F-YPEL2-Turbo-HA fusion protein effectively conjugates biotin onto intracellular proteins as early as 30 min even at 25 µM biotin concentration (Supplemental Information, Fig. S2B). Based on these results, we selected to use 50 µM biotin concentration for subsequent experiments.

### Assessing the effects of YPEL2 on cellular growth

To assess the effects of YPEL2 on cellular growth, 6×10^4^ COS7 cells grown in six-well culture plates for 48 h were transiently transfected with pINDUCER20-MCS vector bearing none (EV), or the 3F-YPEL2 cDNA for 24 h. Cells were then incubated in a fresh medium without or with 10 ng/ml Dox for 96 h with refreshing at every 24 h. At the end of an experiment, cells were collected by trypsinization, and a portion of cells was subjected to counting using a hemocytometer as described previously^4, 58^ or subjected to crystal violet staining^59^ at 96 h. We also subjected cellular extracts collected every 24 h to WB analysis to ensure that cells synthesize 3F-YPEL2 in response to Dox at similar levels throughout the experiment.

In assessing the effects of YPEL2 on cell cycle distribution, we used flow cytometry as we described previously^58^. In short, transfected cells as described for cellular growth for 24 h were collected with trypsinization, washed with PBS, and pelleted. Cells were gently re-suspended in 100 µl of 2%CD-FBS containing PBS, fixed, and permeabilized with ice-cold 70% ethanol overnight. Cells were subsequently incubated with 200 µl of PBS containing propidium iodide (20 µg/ml; Sigma-Aldrich), 200 µg/ml RNase A (Thermo-Fisher), and 0.1% (v/v) Triton X-100 (AppliChem, Germany) for 30 min. Cell cycle analyses were then carried out with flow cytometry (BD Accuri C6 Cytometer; BD Biosciences, San Jose, CA, USA).

For the Annexin V staining, we used Tonbo APC Annexin V Apoptosis Detection Kit (Cytek Biosciences, The Netherlands). Transfected cells as described for cellular growth for 24 h were harvested and washed twice with cold 1X PBS and then re-suspended in 1 ml Stain Buffer. Cells were then centrifuged at 300-400 x g for 5 minutes at room temperature. The cell pellet was resuspended in 100 µl Annexin V Binding Buffer (1X). PE-APC Annexin V conjugate (5 µl) and 7-Amino-Actinomycin (7-ADD) solution (5 µl) were then added to 100 µl of cell suspension which was gently mixed and incubated for 15 min at room temperature in the dark followed by the addition of 400 µl of 1X binding buffer. Cells were then subjected to flow cytometry (BD Accuri C6 cytometer, BD Biosciences). We also used apoptosis inducer doxorubicin (DRC) at 10 or 100 µM concentration for 24 h as a control for the induction of apoptosis in un-transfected cells.

### Dynamic analyses of the putative YPEL2 proximity protein partners with TurboID

To identify proximity interaction partners of YPEL2, we carried out dynamic proximity-dependent biotinylation using the TurboID approach. COS7 cells (2 × 10^6^/10 cm^2^ culture dish of a total of four culture dishes) grown for 48 h were transiently transfected with pINDUCER20-MCS vector bearing none (EV), the Turbo-HA, or the 3F-YPEL2-Turbo-HA cDNA for 24 h. Cells were then subjected to 10 ng/ml Dox treatment for 24 h. Cells were subsequently incubated in a fresh medium containing 50 μM Biotin 1 mM ATP for 1, 3, 6, or 16 h with 10 ng/ml Dox. At the end of a time point, cells were collected with trypsinization and pelleted. Pellets were then washed twice with ice-cold 1x PBS and lysed at RT in 400 µl lysis buffer [50 mM Tris, pH 7.4; 500 mM NaCl; 0.4% SDS; 5 mM EDTA; 2% TritonX; 1 mM DTT with freshly added protease (Roche; 5892970001)] and phosphatase (Roche; 4906845001)] inhibitors. Cell lysates were sonicated for a total of 7.5 min (with 10-sec pulse and 15-sec rest in between pulses) and centrifuged at 7500 rpm for 10 min at 4°C. Protein concentrations were assessed with the QuickStart Bradford Protein assay and 6 mg of protein samples were incubated with 500 µl Streptavidin magnetic beads (NEB, S1420S) overnight. Beads were collected with a magnetic rack and washed twice with Wash Buffer I (2% SDS in water) for 10 min. Beads were then washed once with Wash Buffer II (2% deoxycholate; 1% TritonX; 50 mM NaCl; 50 mM HEPES pH 7.5; 1 mM EDTA) for 10 min, once with Wash Buffer III (0.5% NP-40; 0.5% deoxycholate; 1% TritonX; 500 mM NaCl; 1 mM EDTA; 10 mM Tris pH 8.0) for 10 min, and once with Wash Buffer IV (50 mM Tris, pH 7.4; 50 mM NaCl) for 30 min at RT. A fraction, 5%, of bound proteins were eluted from the streptavidin beads with 50 µl of Laemmli-DTT sample buffer containing 500 nM D-Biotin for WB analyses using the biotin antibody (Ab533494), and the remaining samples were subjected to Mass Spectrometry (MS) analyses. Experiments were carried out two independent times.

#### Protein identification by mass spectrometry

Mass spectrometry (MS) analyses were carried out at the Koç University Proteomic Facility (Istanbul, Türkiye) as we described previously^54^. The protein-bound streptavidin beads were washed with 50 mM NH_4_HCO_3_, followed by reduction with 100 mM DTT in 50 mM NH_4_HCO_3_ at 56 °C for 45 min, and alkylation with 100 mM iodoacetamide at RT in the dark for 30 min. MS Grade Trypsin Protease (Pierce) was added onto the beads for overnight digestion at 37°C (enzyme:protein ratio of 1:100). The resulting peptides were purified using C18 StageTips (ThermoFisher). Peptides were analyzed by C18 nanoflow reversed-phase HPLC (2D nanoLC; Eksigent) linked to a Q-Exactive Orbitrap mass spectrometer (ThermoFisher). The data sets were searched against the human SWISS-PROT database version 2014_08. Proteome Discoverer (version 1.4; ThermoFisher) was used to identify proteins. The final protein lists were analyzed using the STRING v11.5^60^ and DAVID v2022q4^61^ databases.

### Assessing the interaction of Putative interaction Protein Partners with YPEL2

Subtractive analyses of MS results suggested that YPEL2 interacts with a large number of proteins with or without overlapping patterns at different time points. Based on their functional properties and being common at different biotinylation time points, we initially selected ADSS (Adenylosuccinate Synthase), EEF1D (Eukaryotic Translation Elongation Factor 1 Delta), G3BP1 (G3BP Stress Granule Assembly Factor 1), ELAVL1 (ELAV-like RNA Binding Protein 1), and SQSTM1 (Sequestosome1) to verify that they are interacting partners of YPEL2.

#### Cloning

For the generation of cDNAs for ADSS, EEF1D, ELAVL1, G3BP1, and SQSTM1, total RNA from COS7 cells was processed for cDNA, which was used as template in PCR with primer sets designed for each gene based on transcript sequences in the NCBI database (https://www.ncbi.nlm.nih.gov/). PCR amplicons were then cloned into pcDNA 3.1(-) vector that bears in-frame sequences encoding for an amino-terminally located 3F or HA tag at the 5’-end of the multiple cloning site with appropriate restriction enzymes. Constructs were then sequenced for sequence fidelity.

#### Synthesis and intracellular location of putative YPEL2 protein partners

pcDNA 3.1(-) expression vector bearing none (EV) or a cDNA for a putative YPEL2 interaction partner were transiently transfected into COS7 cells and processed for WB or ICC as described above.

#### Co-Immunoprecipitation (Co-IP)

Co-IP was carried out as described previously^54^. Briefly, COS7 cells, 6×10^4^ cells/well, were seeded into six-well tissue culture plates. Cells were transfected with pcDNA 3.1(-) expression vector (0.5 µg/well) bearing the 3F-YPEL2 cDNA and/or the HA-tagged vector driving the expression of the ADSS, EEF1D, ELAVL1, G3BP1, or SQSTM1 cDNA for 48 h. In co-transfections, the total amount of transfected DNA was 1 µg/well. To equalize the total amount of DNA (1 µg/well) in transfections that were carried out with a single construct, 0.5 µg pcDNA 3.1(-) driving a cDNA expression was used together with 0.5 µg pcDNA 3.1(-) bearing no cDNA. Cells were then collected with trypsinization and lysed with M-PER (ThermoFisher; 78,833) containing freshly added protease and phosphatase inhibitors. The protein concentration of lysates was assessed with the Bradford Protein Assay. To block non-specific protein binding to magnetic beads, 500 μg lysates were incubated with non-specific IgG (5 μg) together with 25 μl Protein A/G conjugated magnetic beads at 4 °C for 1 h. The supernatant was transferred to a clean 1.5 ml microcentrifuge tube and the beads were discarded. The pre-cleared lysates were subsequently incubated with 5 µg of the HA or Flag antibody at 4 °C overnight and followed by the addition of 25 μl Protein A/G conjugated magnetic beads at 4 °C for 1.5 h. The beads were then washed two times with 500 μl IP buffer (150 mM NaCl, 10 mM HEPES pH 7.5, 10 mM MgCl2, 0.5% Igepal, protease inhibitors, and phosphatase inhibitors). Bead pellets were resuspended in 30 μl of 2×Laemmli-SDS buffer [187.5 mM Tris–HCl (pH 6.8), 6% (w/v) SDS, 30% glycerol, 150 mM DTT, 0.03% (w/v) bromophenol blue, 2% β-mercaptoethanol] and incubated at 95 °C for 5 min. Samples were then applied to a magnetic field for 30 sec and supernatants were subjected to SDS-10%PAGE for WB analysis using the Flag or the HA antibody followed by the HRP-conjugated VeriBlot (Abcam, ab131366). To ensure that the interactions between proteins are not due to specific tags, we also carried out co-immunoprecipitation with the reverse tag constructs.

#### Proximity ligation assay (PLA)

To further verify the interaction between YPEL2 and ELAVL1 or SQSTM1 we carried out a PLA assay using the Duolink In Situ Red Starter kit (Sigma-Aldrich) as described previously^54^. In brief, COS7 cells (2.5 × 10^4^) grown on glass coverslips in a well of a 12-well tissue culture plate for 48 h were transiently transfected with the expression vectors bearing none, the 3F-YPEL2 and/or HA-ELAVL1, and/or HA-SQSTM1 cDNA for 48 h. Cells were then fixed with 3.2% PFA in PBS for 10 min, permeabilized with 0.1% Triton-X for 5 min, and then blocked with Duolink blocking solution at 37 °C for 30 min. Cells were subsequently probed with the Flag (1:250) and/or the HA (1:500) antibody overnight at 4 °C. Cells were then treated with fluorescent probes for 1 h at 37 °C. Cells were washed with wash buffer A for 10 min at RT and incubated with secondary antibodies conjugated to plus and minus PLA probes for 1 h at 37 °C. After repeating the washing step with wash buffer A for 10 min at RT, cells were incubated with the ligase for 30 min at 37 °C. Cells were then incubated with the polymerase in the amplification buffer for 100 min at 37 °C. Cells were washed in 1x wash buffer B for 20 min and then with 0.01x wash buffer B for 1 min at RT. Duolink *In Situ* Mounting media containing DAPI was used for nuclear staining. Images were captured with a Nikon Eclipse 50i Fluorescence Microscope. ImageJ software was used for image analysis.

### Assessing the formation of stress granules (SGs)

SGs are dynamic phase-separated, membrane-less cytoplasmic ribonucleoprotein assemblies that promote cell survival by condensing translationally stalled mRNAs, ribosomal components, translation initiation factors, and RNA-binding proteins (RBPs)^62–64^. To assess the formation of SGs in COS7 cells in response to various concentrations (0-400 µM) of sodium arsenite (NaAsO_2_, SA, Sigma Aldrich, S7400), as an oxidative stress inducer, we initially examined the intracellular localization of G3BP1 as one of the canonical SG nucleating factors^65^ (Supplemental Information, Fig. S3). For ICC, COS7 cells grown on coverslips in 12-well tissue culture plates were treated without or with 100, 200, and 400 µM SA for 1h. Cells were fixed, washed, blocked, and incubated with a mouse monoclonal G3BP1 antibody (SCBT, sc-365338) followed by an AlexaFluor 488-conjugated goat anti-mouse secondary antibody (BioLegend, 405319). Images were captured with a fluorescence microscope. Results revealed that 200 µM SA is the optimal concentration for inducing SG-like foci in cells without inducing cell death. Based on these preliminary studies, we selected 200 µM of SA as the optimal amount to induce SG formation following 1 h treatment.

To further ensure the identity of SGs, we carried out RNA-fluorescence *in situ* hybridization (RNA-FISH) with an ATTO488-conjugated oligo-dT (50 nucleotides in length) probe (Integrated DNA Technologies, IDT, Belgium) for mRNA detection followed by the identification of G3BP1 protein with ICC. Cells were then fixed in 3,7% formaldehyde in PBS at room temperature for 30 min and permeabilized with 0.4% Triton-X in PBS for 10 min. Cells were washed twice with 1x PBS and then washed twice with 2× saline sodium citrate (SSC) for 5 min. Cells were subsequently incubated with the ATTO488 conjugated oligo-(dT)_50_ probe (16 ng/µl) in hybridization buffer (%50 Formamide, 10% w/v Dextran sulfate, 1 mg/ml Salmon sperm diluted in 2×SSC) at 37° C overnight. Cells were washed with 2× SSC (2×5 min) and 1x PBS (2×5 min). Images were captured with a fluorescence microscope.

#### Assessing co-localization of YPEL and ELAVL1 or G3BP1 in SGs in response to SA

To examine the localization of endogenous YPEL proteins in SGs in response to SA for 1 h, COS7 cells grown on coverslips were subjected to ICC using the pan-YPEL (sc-99727), ELAVL1 (sc-5261), and/or G3BP1 (sc-365338) antibody. We used an Alexa Fluor-647 conjugated secondary antibody for the pan-YPEL antibody and/or a secondary antibody conjugated with Alexa Fluor-488 (Biolegend, 405319) for the ELAVL1 (sc-5261) or G3BP1 antibody. Samples were mounted onto glass slides with a mounting medium containing DAPI for imaging with a fluorescence microscope.

Similarly, we assessed the kinetics of SG formation in response to SA and the disassembly of SGs following the withdrawal of SA. For the formation of SGs, COS7 cells grown on coverslips were subjected to 200 µM SA for 0, 15, 30, 60, and 120 min, and were processed for ICC using the YPEL (sc-99727) or the G3BP1 (sc-365338) antibody. For the disassembly of SG following a 1 h of SA treatment (post-stress, PS), cells were washed and maintained in fresh medium without SA for 0, 1, 2, 3, and 4 h. Cells were then fixed and processed for ICC.

To assess the effects of reduced ELAVL1 levels on the co-localization of the endogenous YPEL proteins in SGs, COS7 cells were transiently transfected with 10 nM of a control siRNA (AllStar) or a siRNA pool that targets ELAVL1 transcripts for 24 h. For WB analysis, cell extracts from COS7 cells grown in six-well tissue culture plates were subjected to SDS-10%PAGE followed by WB using the ELAVL1-specific antibody (sc-5261). Proteins were visualized with an HRP-conjugated secondary antibody (BioLegend 405319). For ICC, cells grown on coverslips and transfected with 10 nM of a control siRNA (Qiagen, AllStar, SI03650318) or a siRNA pool that targets ELAVL1 transcripts (sc-35619) for 24 h were subjected to 200 µM SA for 1 h. Cells were then processed for ICC using the ELAVL1 (sc-5261) or a G3BP1 rabbit polyclonal (13057-2-APG3BP1) antibody. Samples mounted onto glass slides with a mounting medium containing DAPI were subjected to visualization using a fluorescence microscope.

#### Quantification of SGs

To assess the effects of repressed levels of ELAVL1 on the number and size of SGs in the absence or presence of SA, we carried out SG quantification on ICC images using CellProfiler^66^ as an open-source image analysis program (http://www.cellprofiler.org). We initially used the Gaussian Filter method to clean up the images from artifact objects by setting the artifact diameter as 5. For nuclei detection (‘PrimaryObjects’), we used the global threshold followed by the Otsu threshold. The nucleus was used to identify cytoplasm (‘SecondaryObjects’) with the Distance-B method as a parameter of 100 pixels outward from the nucleus. Cytoplasm was then extended a few pixels to keep granules present on the cell surface boundaries through ‘ExpandOrShrinkObjects’ modules. For small circular granules, we used enhancement with the ‘EnhanceOrSuppressFeatures’ module. Enhanced images were finally used to measure the size and number of granules *per* cell in two independent experiments with a total of 100 cells. For statistical analyses, we used a two-tailed Student’s t-test with p<0.05 as the statistical limit of significance.

### Experimental Design and Statistical Rationale

In examining the expression of the YPEL family genes in MCF7 and COS7 cells as well as the E2 signaling on *YPEL2, YPEL3, and YPEL5*, all samples were processed as three biological replicates and three technical repeats. Results were analyzed by a two-tailed unpaired Student’s t-test and depicted as mean ± SD with a p<0.05 as the statistical limit of significance. In assessing the effects of 3F-YPEL on cellular growth and cell cycle distribution, experiments were carried out as three biological replicates. Results were analyzed by a two-tailed unpaired Student’s t-test and depicted as mean ± SD with a p<0.05 as the statistical limit of significance. All other experiments including dynamic proximity labeling, WB, ICC, and PLA were conducted at least two independent times.

## RESULTS

In an effort directed at deciphering the underlying mechanism of E2-ERα actions at genomic levels, we previously identified *Yippee Like 2* (YPEL2) as an E2-ERα responsive gene^4^. Yippee of *Drosophila melanogaster* was first described in a yeast interaction trap screen as a protein physically interacting with the hemolin protein of *Hyalophora cecropia*^7^. Subsequent studies with the sequencing of a cDNA library generated from total RNA from mouse embryonic branchial arch tissue, which is an integral part of the development of the craniofacial complex, led to the identification of a cDNA of 2396 bp in length, with a single open reading frame (ORF) of 357 bp encoding 119 residue Yippee-like 1 (Ypel1) protein that shows very high amino-acid sequence homology to the Yippee protein^8^. Screening of the human expressed sequence tag (EST) with a mouse Ypel1 ORF probe identified the human YPEL1 gene that maps to human chromosome 22q11.2, a region associated with several congenital anomalies involving craniofacial malformation including DiGeorge syndrome and Velocardiofacial syndrome^67^. Sequence comparison at nucleotide levels indicated a high degree of conservation (91.2%) between the human YPEL1 and mouse Ypel1 ORFs and also among many vertebrate and non-vertebrate species^8^. Similarly, a differential gene expression analysis of 32Dcl3 myeloblastic cells in the presence or absence of interleukin 3 identified a gene that encodes the Small unstable apoptotic protein (SUAP), which is now named YPEL3, as a homolog of Yippee and Ypel1^22^. Later studies using the YPEL1 cDNA as a query sequence for the entire human genome revealed four paralogs with remarkably high nucleotide sequence homologies (Supplemental Information, Fig. S4) on four different chromosomes: YPEL2 (17q23.2), YPEL3 (16p11.2), YPEL4 (11q12.1), and YPEL5 (2q23.1) giving rise to the YPEL gene family^5^.

Although the number of YPEL genes varies with species, they are expressed essentially in all eukaryotes^5, 6^. In agreement with previous studies conducted with Northern blot or PCR analyses of human tissue cDNA libraries^5, 22^, our tissue-specific gene expression analyses with the GTEx portal (https://gtexportal.org/home/) (Supplemental Information, Fig. S5) suggest that while *YPEL1* is expressed primarily in the brain and testis; the expression of *YPEL4* is restricted mainly to the brain. On the other hand, *YPEL2*, *3*, and *5* appear to be expressed in all tissues examined. Since *YPEL2, YPEL3*, and *YPEL5* show similar tissue expressions, we initially wanted to explore *in silico* chromatin features of transcription start sites (TSSs) as regions containing putative promoter elements as well as transcription factors/co-regulators associated with the expression of the YPEL2, YPEL3, and YPEL5 genes in the human mammary tissue, including E2 responsive and ERα synthesizing MCF7 cells derived from a breast adenocarcinoma. To accomplish this, we used the Cistrome (http://cistrome.org/) database, which is a resource of human and mouse cis-regulatory information derived from ChIP-seq, DNase-seq, and ATAC-seq chromatin profiling assays to map the genome-wide locations of transcription factor (TF) binding sites^45^. Our analyses using the regulatory potential score (defined as the half-decay distance to the highest value at TSS) of ±10 kb distance to TSS (http://dbtoolkit.cistrome.org/) suggest that TSSs of *YPEL2, YPEL3*, and *YPEL5,* which are located in or vicinity of short CpG islands (http://genome.ucsc.edu), display accessible chromatin structure assessed with DNAse I sensitivity and ATAC-Seq analyses and are decorated with histone proteins modified with H3K27ac and H3K4me3, markers for active transcription and are associated with of POLR2A (RNA Polymerase II Subunit A). This indicates that YPEL2, 3, and 5 are actively transcribed in MCF7 cells (Supplemental Information, Fig. S6). Cistrome analyses further suggest that ERα can be associated with the putative regulatory regions. Moreover, our analyses of the human mammary gland datasets with the Signaling Pathways Project (SPP) portal (https://www.signalingpathways.org/), which is an integrated omics knowledgebase for mammalian cellular signaling pathways^68^ suggest that epidermal growth factor (EGF), the insulin family (Insulin and IGF1), or estrogen signaling is involved in *YPEL2, YPEL3*, and *YPEL5* expressions (Supplemental Information, Fig. S7). Collectively, these results imply that a single signaling pathway in tissue has the potential to co-modulate the expression of the *YPEL* family member genes. Consistent with this, it was shown that the suppression of *MDM4* or *MDM2* transcript levels involved in the mediation of p53 activities with a siRNA approach in MCF7 cells alters the expression of *YPEL2*, *YPEL3*, and *YPEL5*^69^.

The YPEL gene family encodes proteins with very high amino acid sequence identities such that the homology among the YPEL1-YPEL4 proteins is 83.2–96.6%; whereas, YPEL5 shows the lowest homology (43.8%)^5^ as our analyses also indicate (Fig. 1A). It should be noted that there exist two isoforms of YPEL3 resulting from alternative splicing in the first exon of YPEL3 (https://www.ncbi.nlm.nih.gov/gene?Db=gene&Cmd=DetailsSearch&Term=83719). The YPEL3 isoform2 has an additional 38 amino acids at the amino terminus generating a 157 aa long protein with an estimated MM of 17,5; whereas the YPEL3 isoform1 is composed of 119 amino acids with an estimated MM of 13,6 kDa. As YPEL3 isoform1, YPEL1, and YPEL2 are 119 aa in length proteins with estimated MMs of 13,5 kDa. The YPEL4 protein is 127 aa in length with an estimated MM of 14,3 kDa; whereas YPEL5 is composed of 121 amino acids with an estimated MM of 13,8 kDa. The extraordinarily high degree of amino-acid sequence identity among YPEL proteins identifies a consensus sequence of C-X_2_-C-X_19_-G-X_3_-L-X_5_-N-X_13_-G-X_8_-C-X_2_-C-X_4_-GWXY-X_10_-K-X_6_-E for all YPEL proteins^5, 7^. Based on the primary structures of YPEL proteins, our analyses to predict tertiary structures using the AlphaFold^40, 41^ server (https://alphafold.ebi.ac.uk/) with the ChimeraX molecular visualization^42, 43^ program (https://www.cgl.ucsf.edu/chimerax/) suggested that YPEL proteins, except the short sequences at the amino-termini, fold into a superimposable globular structure, or Yippee domain, (Fig. 1B). Moreover, structural homologs of YPEL proteins using the Phyre2^44^ server (http://www.sbg.bio.ic.ac.uk/~phyre2/html/page.cgi?id=index) to infer a potential function(s) for YPEL proteins indicated matches for YPEL proteins with probabilities of more than 80% confidence to the Yippee-like domains of oxidoreductase, RNA binding/hydrolase, ligase/signaling, and MSS4/RABIF-like protein families (Fig. 1C). The oxidoreductase heading includes methionine-S-sulfoxide reductase (MSRA) and the methionine-R-sulfoxide reductase family (MsrB, MsrB1, and MsrB2) of both prokaryote and eukaryotes. The oxidation of methionine by reactive oxygen species (ROS) generates a mixture of methionine-S-sulfoxide and -R-sulfoxide^70^. Sulfoxide reductases catalyze the enzymatic reduction and repair of oxidized-methionine residues generated by redox processes thereby protecting cellular proteins from oxidative stress-induced damage^70^. The MSS4-like header contains the evolutionarily conserved MSS4/RABIF (mammalian suppressor of Sec4/RAB Interacting Factor) superfamily of proteins. Displaying structural similarity to methionine sulphoxide reductases^71^, MSS4/RABIF acts as a holdase chaperone in assisting the ATP-independent non-covalent folding of nucleotide-free Rab GTPases, thereby protecting from degradation^72^. Upregulation of MSS4/RABIF after stress induction is suggested to play a role in protecting cells against programmed cell death by interacting with eukaryotic translation initiation factor 3 subunit f (EIF3F)^73^. Ligase/signaling protein headings comprise the human cereblon protein^74^ which functions as a substrate receptor of the E3 ubiquitin ligase complex and targets proteins for proteolysis through a ubiquitin-proteasome pathway^75^ and plays a role in cellular stress signaling^76^. Ligase heading also includes kinetochore protein mis18 of yeast as well as MIS18A and MIS18B of the MIS18 complex in higher eukaryotes. These proteins through the Yippee-like domain^77^ are involved in the priming of centromeres for recruiting CENP-A critical for cellular proliferation^78^. RNA binding and hydrolase headers include Dicer-related helicase 3, DRH-3^79^, an ortholog of the family of RIG-I/DHX58 (Dicer- and retinoic acid-inducible gene I/RNA binding protein DExH-Box Helicase 58) and MDA5 (melanoma differentiation-associated protein 5).

**Fig 1.**
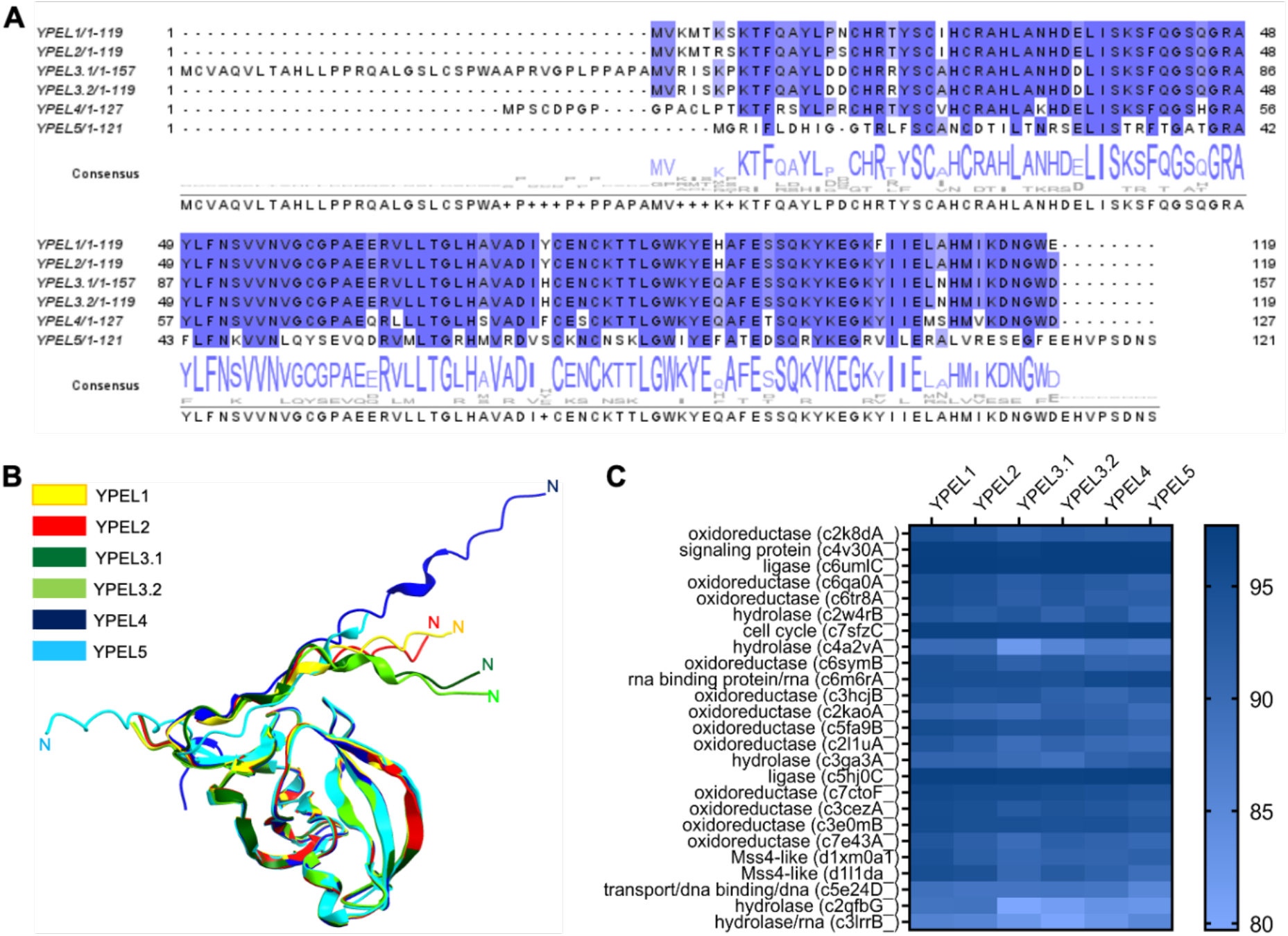
*In silico* analyses of YPEL proteins. (**A**) The alignment of amino acid sequences of YPEL proteins was generated with the Jalview program using the ClustalOmega plug-in. (**B**) Prediction and superimposition of tertiary structures of YPEL proteins were carried out with the AlphaFold server using the ChimeraX molecular visualization program. (**C**) The Phyre2 web tool was used for the homology modeling of YPEL proteins.

Potential commonalities in the regulation of the *YPEL* expressions, the evolutionary conservation in nucleotide sequences of ORFs, encoded amino acid sequences, and predicted functions of YPELs imply shared activities for YPEL proteins in cellular processes. However, this also poses experimental challenges in deciphering the function of a YPEL protein when co-expressed/synthesized with other YPELs. For example, our analyses, using the Cancer Dependency Map portal (DepMap, DM, https://depmap.org/portal/), of the average expression of the YPEL gene family members in breast cancer cell lines (DM-BCCL) or only in MCF7 cells (DM-MCF7) together with our results from MCF7 cells with qPCR (MCF7) suggest that YPEL2, 3, and 5 genes are expressed in these cell lines (Fig. 2A). Moreover, treatment of MCF7 cells, grown for 72 h in CD-FBS to reduce the endogenous estrogen hormone concentrations, with E2 (10^-9^ M) for 24 h led to suppression of *YPEL2*, *YPEL5*, and as shown previously for *YPEL3*^14^, expressions compared to control cells treated with ethanol (EtOH) (Fig.2B). E2 treatment on the other hand augmented the expression of *TFF1*, an E2-responsive gene that we used as a control^49^. The suppression of *YPEL2, 3, and 5* expressions is ER-dependent, this is because 10^-7^ M Imperial Chemical Industries 182,780 (ICI), a complete ER antagonist^80^, effectively blocked the E2-mediated repression. Thus, E2-ER signaling mediates the expression of *YPEL2, YPEL3*, and *YPEL5* in MCF7 cells. We also observed in COS7 cells grown under steady-state conditions that all YPEL genes are expressed at varying levels (Fig. 2C).

**Fig. 2.**
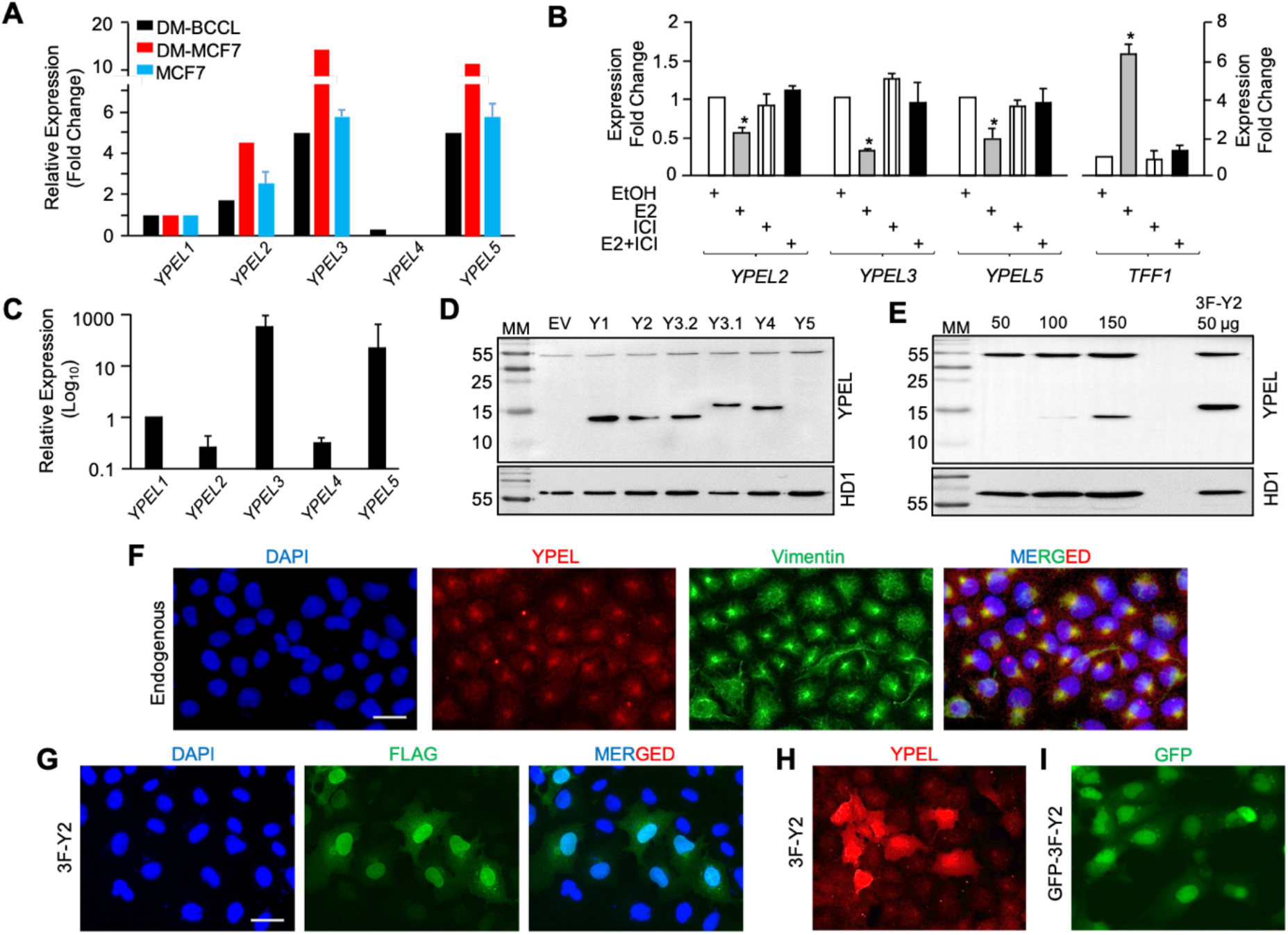
Expression of the YPEL family genes in cell lines. (**A**) Based on data from the DepMap portal, the average expression of the YPEL gene family members relative to the expression of *YPEL1* in breast cancer cell lines (DM-BCCL) or only in MCF7 cells (DM-MCF7) is shown. For comparison, our RT-qPCR analyses for the relative expression of YPEL genes in MCF7 cells (MCF7) are also indicated. (**B**) To assess whether or not E2-ERα signaling is involved in the regulation of *YPEL2*, *3*, and *5* expressions, MCF7 cells grown for 72 h in CD-FBS containing growth medium were treated without (EtOH as control) or with 10^-9^ M E2, and/or 10^-9^ M ICI, a complete ER antagonist, for 24h. cDNAs generated from total RNA were subjected to qPCR. We also used the E2-ERα responsive gene TFF1 as a control. Star indicates a significant change compared to EtOH control. (**C**) Relative to *YPEL1*, the expression of YPEL genes in COS7 cells was assessed by qPCR using cDNA from total RNA obtained from COS7 cells grown in steady-state conditions. (**D**) To evaluate the synthesis of YPEL1-5 proteins, which Y3.1 and Y3.2 indicate short and long variant YPEL3 proteins, COS7 cells were transiently transfected with the expression vector pcDNA3.1(-) bearing none (EV) or cDNA for a YPEL protein. Total protein extracts of COS7 cells grown in steady-state condition were subjected to SDS-15%PAGE followed by WB using a pan-YPEL antibody (SCBT, sc99727) and an HRP-conjugated goat-anti-rabbit secondary antibody (Advansta R-05072–500). Membranes were re-probed with an antibody specific to HDAC1 (Abcam, ab19845). Molecular masses (MM) in kDa are indicated. (**E**) To assess the synthesis of endogenous YPEL proteins in COS7 cells, protein extracts ranging from 50 to 150 µg were subjected to WB using the pan-YPEL antibody. We also used protein extracts from COS7 cells transiently transfected with pcDNA3.1(-) bearing the 3F-YPEL2 cDNA for comparison. Membranes were re-probed with the HDAC1 antibody (Abcam, ab19845). Molecular masses (MM) in kDa are indicated. (**F**) To assess the intracellular localization of endogenous YPEL (YPEL) proteins, COS7 cells grown on coverslips were subjected to ICC using the pan-YPEL antibody followed by an Alexa Fluor 594-conjugated goat anti-rabbit secondary antibody (Abcam, ab150080) or a vimentin (sc-6260) antibody followed by an Alexa Fluor 488-conjugated goat anti-mouse secondary antibody (Abcam, ab150113). DAPI was used to indicate the nucleus. The scale bar is 20 µm. (**G & H**) The intracellular localization of 3F-YPEL2 was evaluated in transiently transfected cells with the use of the Flag (**G**) or the pan-YPEL (**H**) antibody or GFP fusion (**I**). DAPI staining indicates the nucleus. The scale bar is 20 µm.

The experimental challenge was also evident when we attempted to detect the YPEL2 protein with commercially available and purportedly YPEL2-specific antibodies in WB or ICC studies in which we could not detect a YPEL protein using these YPEL2 antibodies (data not shown); despite the fact that we effectively observed the synthesis of 3F-YPEL2 as well as 3F-tagged YPEL proteins with ICC and WB using the Flag antibody in MCF7 or COS7 cells upon ectopic expressions. Of the antibodies we tested, a pan-YPEL antibody, which is an affinity-purified polyclonal antibody raised against a peptide that corresponds to an internal region of YPEL1, with amino acid sequences largely common within YPEL2, 3, and 4 of human origin (SCBT, S-14 Ypel antibody, sc99727), detected YPEL1, 2, 3, and 4 of human origin (SCBT, S-14 Ypel antibody, sc99727), detected YPEL1, 2, 3, and 4 but not YPEL5 in WB of COS7 cells (Fig. 2D). Similarly, this antibody also detected, albeit at low levels, a protein species with electrophoretic mobility of about 13-14 kDa, likely corresponding to YPEL1, YPEL2, YPEL3 subtype1, and/or YPEL4 proteins of COS7 cells in WB (Fig. 2E). This observation is supportive of a previous finding using an in-house pan-YPEL polyclonal antibody raised against a synthetic peptide corresponding to the carboxyl-terminal amino acid sequences of YPEL1-4 proteins that COS7 cells synthesize a YPEL protein(s)^5^. The use of this in-house pan-YPEL antibody also suggested that YPEL(s) localizes to the nucleus and nucleus periphery including the centrosomic region and nucleoli during interphase, and around the mitotic apparatus during the mitotic phase of COS7 cells^5^. Similarly, mouse Ypel1 as a GFP-fusion protein is reported to localize to the nucleus in transfected NIH3T3 mouse fibroblast cells^8^. Likewise, YPEL3 localization is suggested to be weakly nuclear with punctate perinuclear staining^81^. On the other hand, YPEL3, using a YPEL3-specific antibody, is reported to show diffuse staining including nucleus and cytoplasm^25^. The pan-YPEL antibody we used to detect the Flag-YPEL proteins in WB analyses reveals staining in un-transfected COS7 cells encompassing both the cytoplasm and the nucleus as well as denser staining puncta at the nuclear periphery reminiscent of the centrosomic region^5^ (Fig. 2F). Similar locations were also observed in transfected COS7 cells synthesizing 3F-YPEL2 (Fig. 2G & H) detected with the Flag (Fig 2G) or the pan-YPEL antibody (Fig. 2H). Likewise, the GFP-3FYPEL2 fusion protein showed a diffuse intracellular location (Fig. 2I).

The presence of a YPEL protein in COS7 cells suggests that YPEL protein(s) has functional roles in these cells. We, therefore, selected to use COS7 cells to assess YPEL2 functions. Although we were aware that co-expressions/co-syntheses of YPELs with functional commonalities could compensate for the effects of targeted downregulation of YPEL2, we nevertheless initially selected to use siRNA approaches to assess the role of YPEL2 in cellular processes. However, high sequence nucleotide homologies among YPEL transcripts (Supplemental Information, Fig. S4) rendered the design of siRNAs with various tools to target specifically YPEL2, any or all members of the YPEL family, to be difficult. In several attempts with our design or commercially available siRNAs, we were unsuccessful in effectively altering the YPEL expressions in COS7 cells (data not shown). We, therefore, used an overexpression system to examine YPEL2 functions. To circumvent possible adverse effects of YPEL2 overexpression on cellular phenotypes, as we observed acute cell death in MCF7 cells upon overexpression (data not shown), we decided to use an inducible expression approach. pINDUCER20 is a tightly controlled tetracycline-inducible lentiviral single-vector that provides a constitutive expression of the reverse tetracycline transactivator 3 (rtTA3) and contains promoter sequences composed of tetracycline-responsive elements (TRE) for transgene expression^55^. Upon binding to tetracycline, or its stable analog Dox, the constitutively synthesized rtTA3 induces transgene expression from the TRE-driven promoter^55^. We inserted the 3F-YPEL2 cDNA with appropriate restriction sites into pINDUCER20-MCS, which we modified by inserting a multiple cloning site (MCS) to ease transgene cloning. We then transiently transfected COS7 cells with pINDUCER20-MCS bearing none or the 3F-YPEL2 cDNA. We observed by WB and ICC that Dox effectively induced 3F-YPEL2 synthesis, the levels of which were correlated with Dox concentrations: The minimal amount of Dox required for the maximal amount of 3F-YPEL2 synthesis was 10 ng/ml (Fig. 3A). Independent of Dox concentrations, 3F-YPEL2 showed diffuse intracellular staining, as we observed with the endogenous YPEL1-4 proteins (Fig. 2F-H), including the nucleus and cytoplasm (Fig. 3B) which we further confirmed with WB using fractionated cellular proteins (Fig. 3C).

**Fig. 3.**
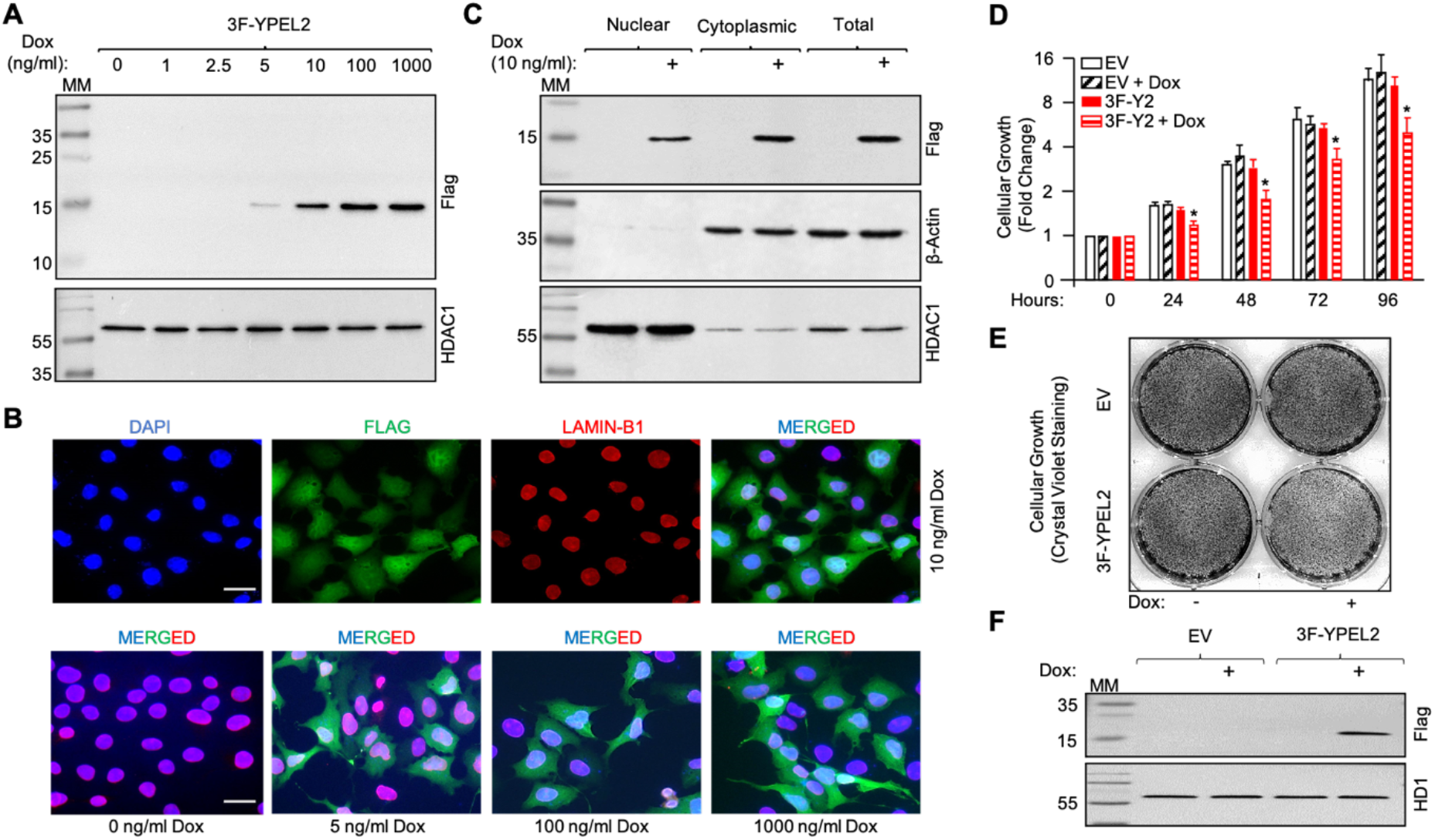
Assessing effects of 3F-YPEL2 on the growth of COS7 cells. (**A**) COS7 cells were transiently transfected with the pINDUCER20-MCS vector bearing none (EV) or the 3F-YPEL2 cDNA for 24 h. Cells were then treated without (0) or with varying concentrations (1-1000 ng/ml) of Dox for 24 h. Total cell extracts (50 µg) were then subjected to WB using the Flag antibody followed by an HRP-conjugated goat-anti-mouse secondary antibody (Advansta R-05071-500). Membranes were re-probed with the HDAC1 antibody. Molecular masses (MM) in kDa are indicated. (**B**) Transiently transfected cells with pINDUCER20-MCS bearing the 3F-YPEL2 cDNA for 24 h were then treated with 0, 5, 10, or 100 ng/ml Dox for 24 h. Cells were subsequently fixed, permeabilized, washed, and stained with the Flag antibody followed by an Alexa Fluor 488-conjugated goat anti-mouse secondary antibody (Abcam, ab150113) or a Lamin B1 antibody (Abcam, ab16048) followed by an Alexa Fluor 647-conjugated goat anti-rabbit secondary antibody. DAPI was used to indicate the nucleus. The scale bar is 20 µm. (**C**) To assess the subcellular levels of inducibly synthesized 3F-YPEL2 in transiently transfected cells treated without or with various concentrations of Dox for 24 h, fractionated nuclear and cytoplasmic protein extracts were subjected to WB using the Flag antibody. Membranes were re-probed with an antibody specific to β-actin (Abcam, ab8227) or HDAC1 (Abcam, ab19845). Molecular masses (MM) in kDa are indicated. (**D**) To assess the effects of 3F-YPEL2 on cellular growth, COS7 cells transiently transfected with pINDUCER20-MCS bearing none (EV) or the 3F-YPEL2 cDNA for 24 h were treated without or with 10 ng/ml Dox for 24 h intervals up to 96 h. At every 24 h, cells were collected and counted with a hematocytometer. The asterisk indicates significant changes (P<0.05) in cellular growth depicted as fold changes. (**E**) Crystal violet staining for cellular growth and (**F**) WB analysis with the Flag antibody of COS7 cells inducibly synthesizing 3F-YPEL2 at 96 h are shown.

Since the YPEL family proteins appear to be involved in alterations of cellular phenotype, we initially assessed the effect of YPEL2 on cellular proliferation in COS7 cells transfected with pINDUCER20-MCS bearing none or the 3F-YPEL2 cDNA for 24 h. Cells were then treated without or with 10 ng/ml Dox for up to 96 h with fresh medium every 24 h. We found by cell counting (Fig. 3D) and crystal violet staining, which is shown at 96 h (Fig. 3E), that 10 ng/ml Dox-induced 3F-YPEL2 synthesis reduces cellular proliferation (Fig. 3F). Interestingly, the decline in cellular proliferation appeared to occur only at 24 h Dox treatment without altering the rate of proliferation at subsequent time points which remained the same throughout (Fig. 3D) even though cell synthesized 3F-YPEL2 at levels that were similar at all time points (Supplemental Information, Fig. S8A). However, we could not ascertain how 3F-YPEL2 synthesis led to the repression in cell growth at 24 h at which we observed no significant changes in cell cycle distribution assessed with flow cytometry (Supplemental Information, Fig. 8B) or in cell death by Annexin V staining (Supplemental Information, Fig. 8C).

### Dynamic analyses of proximity protein interaction network of YPEL2

Although YPEL2 appears to be involved in cellular proliferation, the underlying mechanism is unclear. Since proteins function within protein interaction networks critical for cellular processes, we reasoned that the identification of interacting partners of YPEL2 in a time-dependent manner could provide critical information about YPEL2 functions. To explore this issue, we used the TurboID proximity labeling approach, a derivative of the BioID system^56, 82^ which we utilized previously^54^. TurboID uses a biotin ligase engineered through yeast display-based directed evolution of a promiscuous mutant of *E. coli* biotin ligase, BirA* that catalyzes proximity biotin labeling of proteins with higher efficiencies and faster kinetics compared to BirA*. This allows the investigation of protein interaction networks dynamically^57^. Based on our initial analyses that TurboID can be effectively used for dynamic protein interaction profiling of YPEL2 (Supplemental Information, Fig. S1 & S2), we cloned the Turbo-HA, 3F-YPEL2, or 3F-YPEL2-Turbo-HA cDNA into pINDUCER20-MCS to efficiently dictate the timing and the level of protein synthesis in transiently transfected cells to minimize potential adverse effects of transgene overexpression. We transiently transfected COS7 cells with the expression vectors for 24 h. Cells were then treated with 10 ng/ml Dox for 24 h to induce 3F-YPEL2 synthesis. We then treated cells without or with 50 μM biotin and 1 mM ATP for 1, 3, 6, or 16 h in the presence of 10 ng/ml Dox followed by WB (Fig. 4A & 4B) and ICC (Fig. 4C) using an antibody specific for the Flag, HA, or biotin. Results, shown are biotin labeling for 3 h, revealed that cells only in the presence of Dox synthesize 3F-YPEL2, Turbo-HA, or 3F-YPEL2-TurboID-HA with an expected molecular mass (MM) of approximately 17, 37, or 54 kDa protein respectively (Fig. 4A) and the proteins show diffuse intracellular staining encompassing both the nucleus and cytoplasm independently of the exogenously added biotin (Fig. 4C). Importantly, the detection of many biotinylated proteins only in the presence of biotin in transfected cells synthesizing 3F-YPEL2-TurboID-HA indicates that the fusion protein is functional as well. The detection of Turbo-HA or 3F-YPEL2-TurboID-HA with the Biotin antibody in the absence of exogenously added biotin also suggests self-labeling of the proteins with residual endogenous biotin (Fig. 4A).

**Fig. 4.**
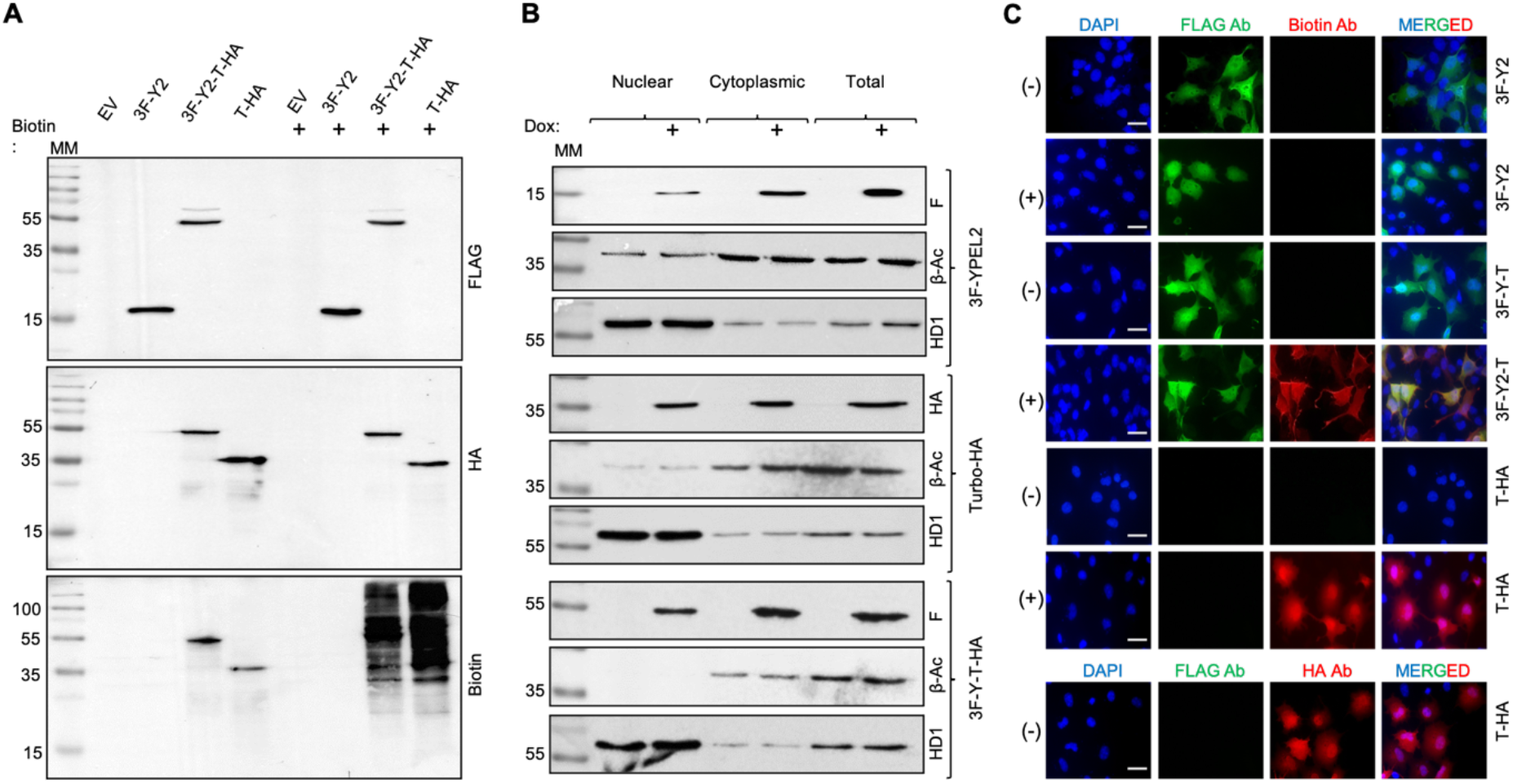
Synthesis, subcellular location, and effects of TurboID constructs on the biotinylation of intracellular proteins in COS7 cells. (**A**) COS7 cells were transfected with pINDUCER20-MCS carrying none (EV), the 3F-YPEL2 (3F-Y2), 3F-YPEL2-Turbo-HA (3F-Y2-T-HA), or Turbo-HA (T-HA) cDNA. 24 hours after transfection, cells were treated without or with 10 ng/ml Dox to induce protein synthesis for 24 h in the absence or presence of biotin (50 μM) for the biotinylation of endogenous proteins. Cells were then collected and equal amounts (50 µg) of protein extracts were subjected to SDS-10%PAGE electrophoresis followed by WB analyses using the Flag, HA, or Biotin (Abcam, ab53494) antibody. Molecular masses are indicated in kDa. (**B**) To assess subcellular distributions of 3F-YPEL2 (3F-Y2), Turbo-HA (T-HA), or 3F-YPEL2-Turbo-HA (3F-Y2-T-HA) proteins following 10 ng/ml Dox induction for 24 h in transiently transfected cells, cytoplasmic, nuclear, or the total protein extracts (50 µg) were subjected to WB analyses wherein β-actin (β-Ac, ab8227) and HDAC1 (HD1) were used as the cytoplasmic and nuclear protein loading controls. Molecular masses are depicted in kDa. (**C**) Intracellular locations of the TurboID constructs in the transiently transfected cell grown on coverslips without (-) with Biotin (+) were assessed with the Flag antibody followed by the Alexa Fluor 488-conjugated secondary antibody (Abcam, ab150113), or HA and/or biotin antibody followed by the Alexa Fluor 594-conjugated secondary antibody (Abcam, ab150080). DAPI staining indicates the nucleus. The scale bar is 20 µm.

Based on these results, and our preliminary studies (Supplemental Information, Fig. S1 & S2), we carried out large-scale biotin labeling of intracellular protein for MS analyses. COS7 cells transiently transfected with pINDUCER20-MCS bearing i) none (EV), ii) the Turbo-HA, or iii) the 3F-YPEL2-Turbo-HA cDNA for 24 h were subjected to 10 ng/ml Dox for 24 h. Cells were then incubated in a fresh medium containing 50 µM biotin and 1 mM ATP for 1, 3, 6, and 16 h. Cells were lysed and biotinylated proteins in cell lysates were captured with streptavidin-conjugated magnetic beads. Protein fragments following on-bead tryptic proteolysis of the captured proteins were subjected to MS analyses. MS identified many proteins from each group conducted as two biological replicates. Subtractive analyses of identified proteins generated from cells transfected with pINDUCER20-MCS bearing i) none (EV), ii) the Turbo-HA, or iii) the 3F-YPEL2-Turbo-HA cDNA revealed 130, 99, 157, and 115 specific proximal interactors of YPEL2 at 1, 3, 6, and 16 h respectively (Supplementary Information, Table S2). We found that a large number of proteins was present only at a distinct time point, some were common at specific time points or all time points examined (Fig. 5A & 5B).

**Fig. 5.**
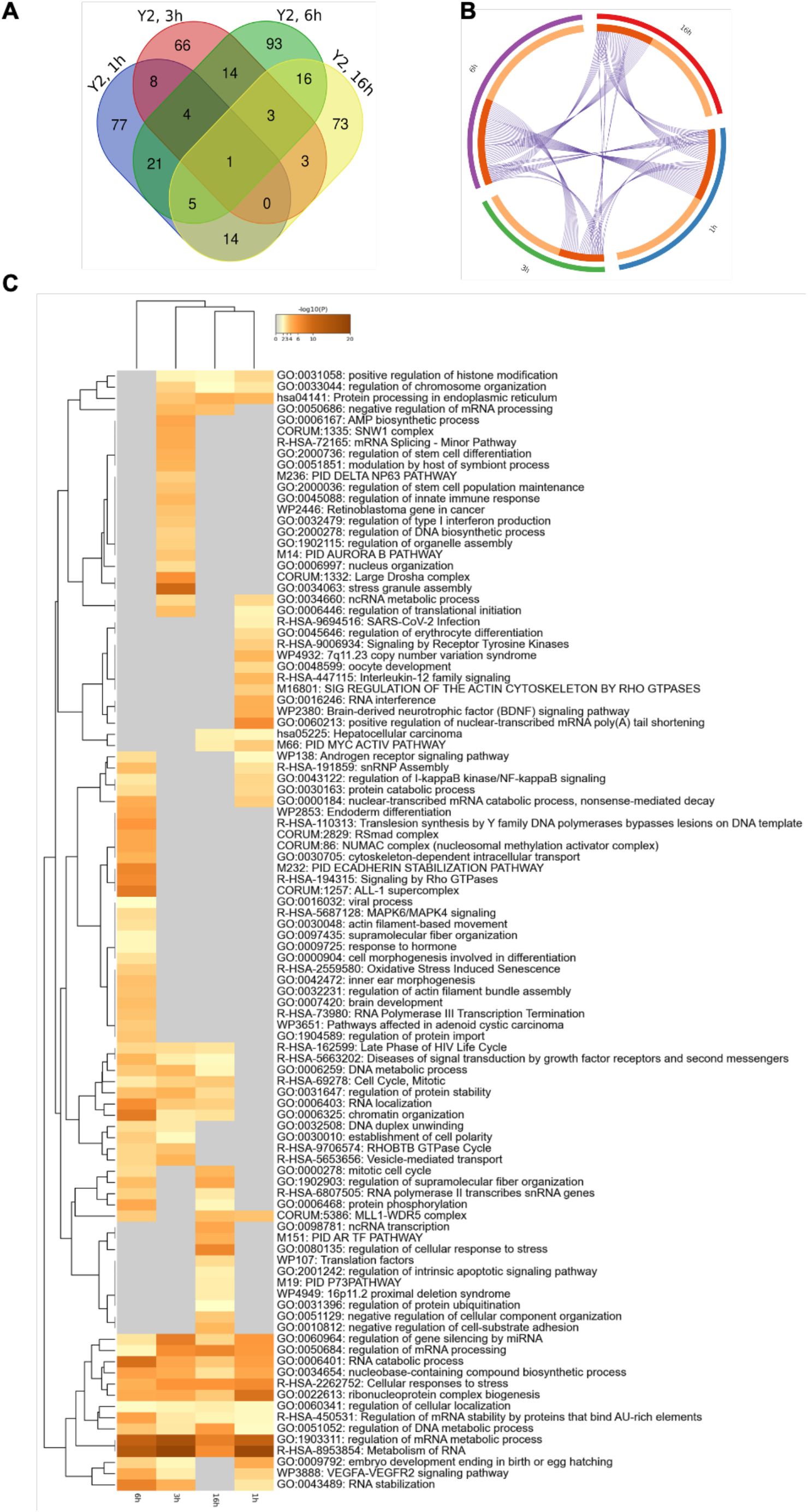
*In silico* analysis of proximal interaction partners of YPEL2. (**A**) Intersections of proximal protein partners of YPEL2 at different time points were constructed using a Venn diagram tool (https://bioinformatics.psb.ugent.be/webtools/Venn/). (**B**) The Circos plot generated with Metascape indicates time-dependent (outside arcs with distinct colors for each time point) proximal interaction partners of YPEL2 that are shared among time points (inside arcs with a dark orange color that are linked with purple lines). (**C**) A hierarchically clustered heatmap of enriched terms related to functions of proteins across all time points colored by p-values was generated by Metascape.

It should be noted that of the previously identified proteins STRN as a YPEL1 interacting protein^83^; SH2D4A^84^, and SRKP2^85^ as YPEL2 interactors; LARP4, SH2D4A, SPG21, SRPK2, TP53, and TRIP6 as YPEL3 interacting partners^84^; DDX5^86^, ELAVL1^87^, LARP4^88^, PFDN5^84^, and RANBP9^6, 86^ as YPEL5 interacting proteins are also present in our protein interaction network of YPEL2. This suggests YPEL2 and other YPEL proteins function similarly.

To assess ontologies of putative interacting proteins of YPEL2 for biological functions, we analyzed the MS results with the Metascape portal (https://metascape.org/), which is an integrated web-based system that allows functional enrichment, interactome analysis, gene annotation, and membership search^46^. Results revealed that the profile of proximity interaction partners of YPEL2 changes dynamically generating a temporal enrichment of Gene Ontology (GO; http://geneontology.org/) and Reactome pathway (https://reactome.org/) terms (Fig. 5C). We find common enrichment of proteins at all time points in GO and Reactome terms that encompass RNA and mRNA metabolic processes, ribonucleoprotein complex biogenesis, regulation of gene silencing by miRNA, and cellular responses to stress, as the secondary neighbor analyses of YPEL2 partners at all time points similarly indicate (Supplemental Information, Fig. S9). Proximity interaction partners of YPEL2 involved in RNA metabolic processes include CELF1, DDX6, DDX52, EEF1D, EIF4H, ELAVL1, G3BP1, HNRNPA0, HNRNPDL, LARP4B, PTBP1, SMN1, SQSTM1, and UPF1; whereas, ARID1A, ARID2, EMSY, HELLS, MBD3, DNMT1, KMT2D, RUVBL1/2, NCOA3 & 5, NCOR1/2, POLA1, POLA2, are sub-grouped in DNA- and chromatin-associated processes. Proteins sub-grouped in cellular responses to stress include AGO2, ANAPC, PN1, BAG3, GFPT1, HSP90B1, MAP4K4, RPS, STAT3, TNRC6A, TP53, and TUBB4B. Despite the distinct identities of proximal interaction partners of YPEL2 at specific time points, similar GO-term enrichments at all time points suggest the interdependency of regulatory networks that YPEL2 takes part in.

Furthermore, and importantly, the identities of a substantial number of the putative protein partners of YPEL2 at all time points appear to be coincident with proteins involved in the formation of processing bodies (p-bodies, PBs)^89^, and stress granules (SGs)^90^ as cytoplasmic ribonucleoprotein (RNP) assemblies (Fig. 6A & B). These include canonical markers of PBs including AGO2, DCP1A, EDC3, and TNRC6A, markers of SGs comprising ATXN2, ELAVL1, G3BP1, NCOA3, PABPC1, SQSTM1, TARDBP, and USP10 proteins as well as proteins shared between SGs and PBs including AGO2, DDX2, XRN1. RNPs including PBs and SGs are membrane-less dynamic networks of multivalent protein-protein and protein-RNA condensates that form through liquid-liquid phase separation (LLPS) for RNA storage and/or processing^64, 91–95^. Of RNPs, PBs are constitutively present in the cytoplasm. PBs contain mRNAs that encode proteins involved in a variety of regulatory processes as reservoirs for rapid adaptation to gene expressions in response to internal and external clues^89, 95^. Whereas, cytoplasmic SGs are formed as defense mechanisms to protect cells against adverse effects of specific types of stresses including osmotic and oxidative stresses which also increase the size and/or number of PBs^64, 91–95^. In stressed cells, PBs are often found adjacent to SGs and exchange content with each other and the cytoplasm^64, 91–95^. SG-associated proteome and transcriptome profiles show that SGs sequester primarily untranslated mRNPs derived from mRNAs stalled in translation initiation including pre-initiation complexes for storage during a period of stress, and once the stress is attenuated for re-initiation of translation or degradation through autophagy^64, 91–95^. It is therefore suggested that SGs are critical for translation reprogramming by dynamically partitioning translationally silenced mRNAs. This facilitates the preferential translation of priority mRNAs from the cytoplasm or stored pool in PBs for cellular stress responses^93, 95^.

**Fig. 6.**
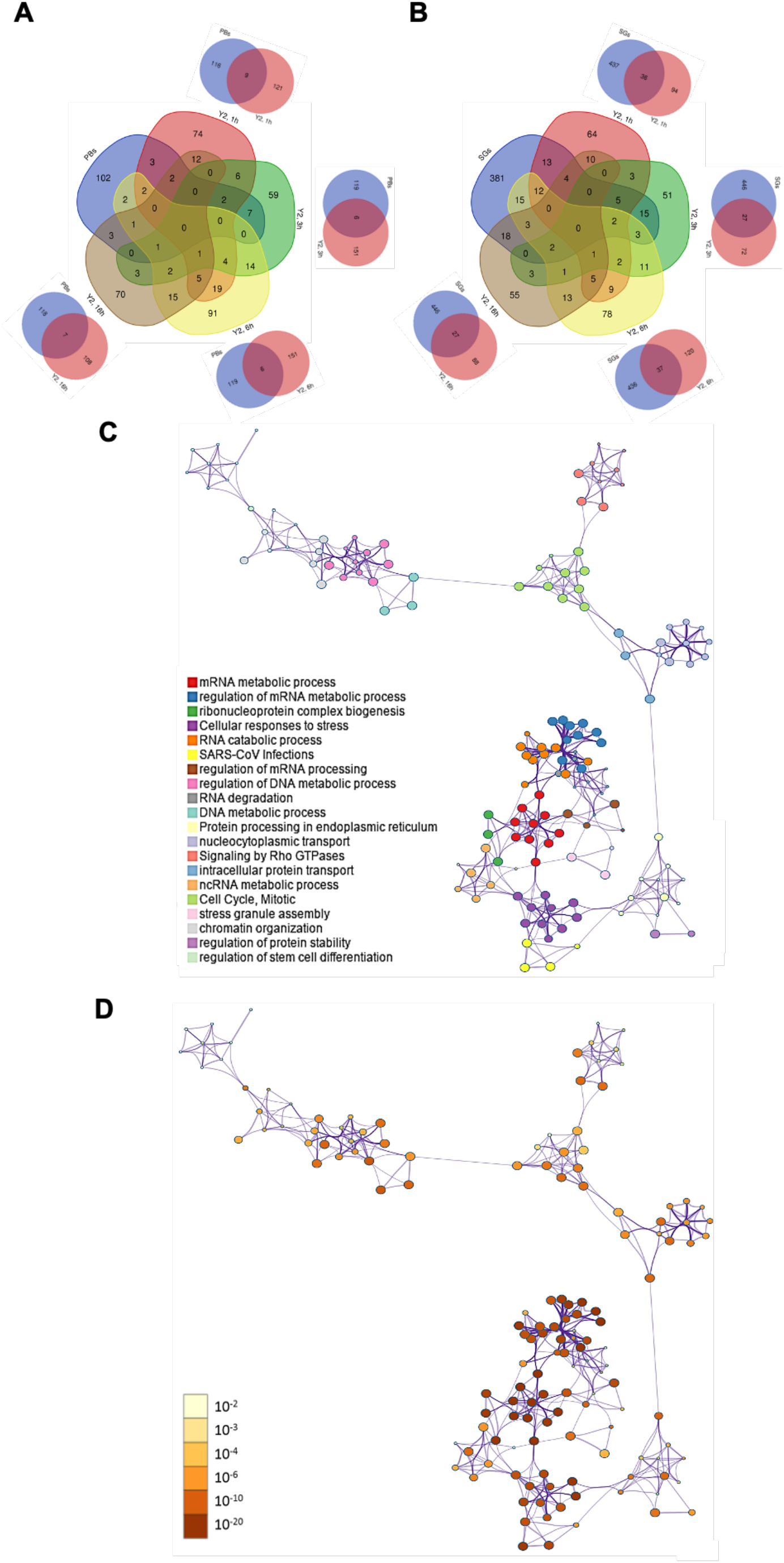
Cumulative network analysis of proximal interaction partners of YPEL2 at all points. (**A & B**) Intersections of proximal protein partners of YPEL2 at different time points with proteins of (**A**) processing bodies^89^, PBs, or (**B**) stress granules^90^, SGs, curated in the RNAgranuleDB database, version 2.0, Gold Standart (http://rnagranuledb.lunenfeld.ca/) were generated using a Venn diagram tool. (**C**) A network of enriched terms using all proximal interaction partners of YPEL2 is generated with Metascape and visualized with Cytoscape. A circle node where size is proportional to the number of proteins specific to a term is colored represented. Nodes that share the same cluster identity are represented close to each other. (**D**) The network of enriched terms is represented by colored p-values, where terms containing more proteins indicate more significant p-values.

Our findings that the putative YPEL2 partners include proteins involved in RNA metabolic processing, in the formation and/or sustaining PBs and SGs, together with the network analyses of all-time points with Metascape (Fig. 6C & D), imply the involvement of YPEL2 responses to stress in under both steady-state and stress conditions, which could result in alterations in mitotic cell cycle processes and, as our results suggest, cellular proliferation.

### Assessing the interaction of YPEL2 with putative protein partners

Based on functional importance derived from our Metascape analyses, we selected ADSS, EEF1D, ELAVL1, G3BP1, and SQSTM1 for the initial interaction screening to test whether or not they are protein partners of YPEL2.

ADSS (Adenylosuccinate synthetase-1 and −2 with estimated MMs of 50,2 and 50,09 kDa, respectively) plays a critical role in the *de novo* and the salvage pathway of purine nucleotide biosynthesis by carrying out initial steps in the biosynthesis of adenosine monophosphate (AMP) from inosine monophosphate^96^. EEF1D (Eukaryotic Translation Elongation Factor 1 Delta) with a predicted MM of 71.4 kDa is a subunit of the elongation factor-1 complex and functions as a guanine nucleotide exchange factor by stimulating the exchange of GDP to GTP in EEF1A1^97^. ELAVL1 (ELAV Like RNA Binding Protein 1), an RNA-binding protein with an estimated 36 kDa MM, binds to poly-U and AU-rich elements (AREs) in the 3’-UTR of target mRNAs and increases their stability^98, 99^. ELAVL1 is also involved in SG maturation^98, 99^. G3BP1 (G3BP stress granule assembly factor 1) with an estimated MM of 52 kDa is an RNA-binding protein and a key component of SG assembly^63, 100^. Localized also in SGs, SQSTM1 (Sequestosome-1) with a MM of 47,7 kDa is an autophagy receptor required for selective macro-autophagy by functioning as a bridge between polyubiquitinated cargo and autophagosomes^101, 102^. To validate the interaction between YPEL2 and a putative interacting partner, we transiently transfected COS7 cells with the expression vector pcDNA3.1(-) bearing none, 3F-YPEL2 and/or HA-ADSS, HA-EEF1D, HA-ELAVL1, HA-G3BP1 or HA-SQSTM1 cDNA for 48 h. Our preliminary studies using Co-IP did not detect an interaction between 3F-YPEL2 and HA-ADSS, HA-G3BP1, or HA-EEF1D (Supplemental Information, Fig. S10). We found, on the other hand, that 3F-YPEL2 interacts with HA-ELAVL1 (Fig. 7A-D) or HA-SQSTM1 (Fig. 7E-H).

**Fig. 7.**
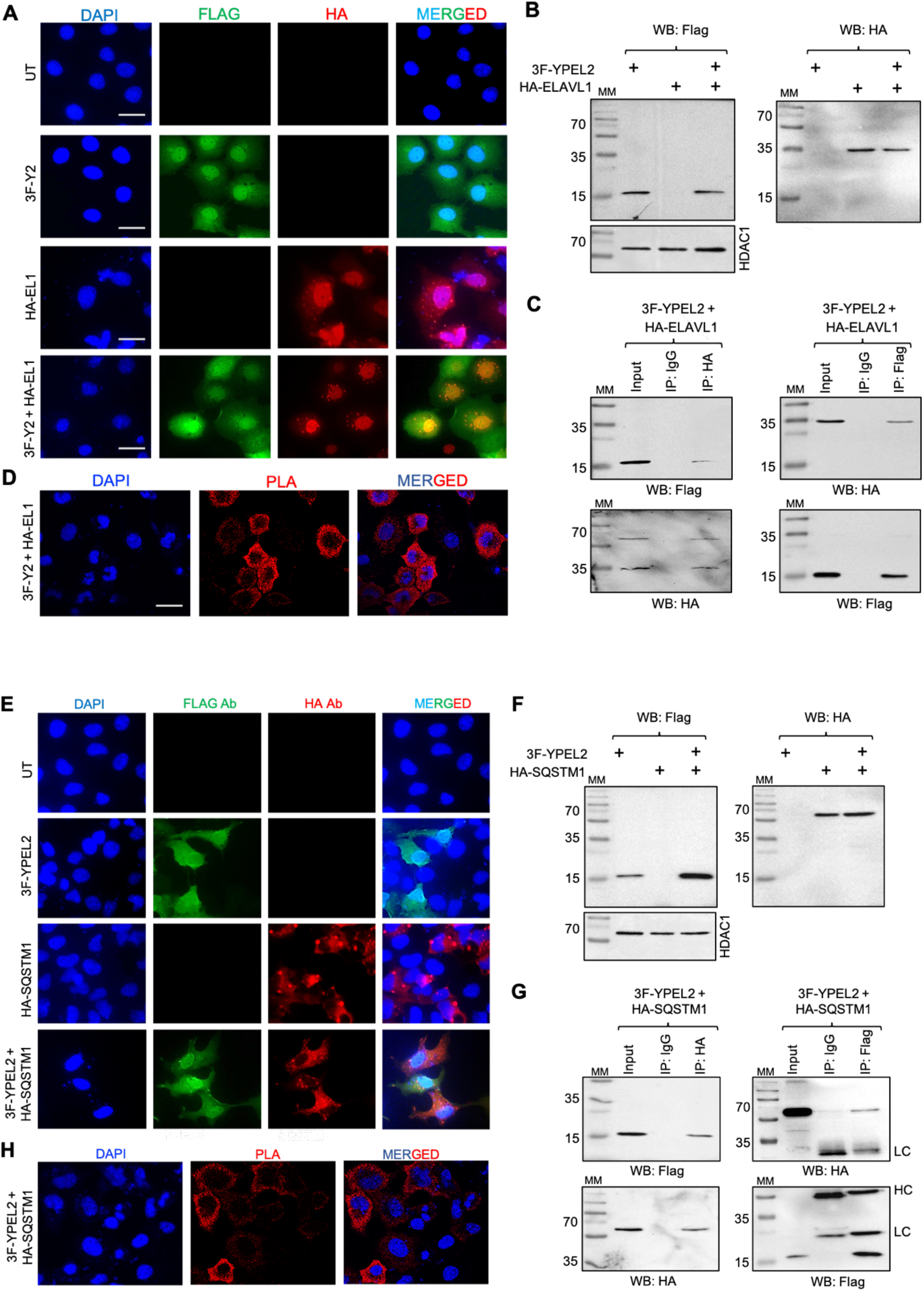
Interaction of 3F-YPEL2 and HA-ELAVL1. (**A**) To assess the intracellular localization of 3F-YPEL2 and/or HA-ELAVL1, COS7 cells grown on coverslips un-transfected (UT) or transiently transfected for 36 h with the expression vector pcDNA3.1(-) bearing the 3F-YPEL2 or HA-ELAVL1 cDNA were stained with the Flag or the HA antibody. DAPI was used to indicate the nucleus. The scale bar is 20 µm. (**B**) To examine the protein synthesis, COS7 cells were transfected (+) with the expression vector bearing 3F-YPEL2 and/or HA-ELAVL1 cDNA. The synthesis of proteins was assessed by WB using the Flag or the HA antibody. HDAC1 used as a loading control was probed with the HDAC1 antibody. (**C**). The cellular extracts (500 μg) of transiently co-transfected COS7 cells were subjected to Co-IP with the HA, Flag, or isotype-matched IgG. 50 μg of lysates was used as input control. The precipitates were subjected to SDS-10%PAGE followed by WB using the Flag or the HA antibody. Molecular masses (MM) in kDa are indicated. (**D**) To assess *in cellula* interaction of 3F-YPEL2 and HA-ELAVL1, the proximity ligation assay was carried out in transiently transfected COS7 cells grown on coverslips. Cells were fixed, permeabilized, blocked, and probed with the HA and/or the Flag antibody. Cells were then subjected to fluorescent probes for circular DNA amplification for proximity interaction foci. DAPI was used for nuclear staining. Images were captured with a fluorescence microscope. The scale bar is 20 µm. (**E**) The intracellular localization, (**F**) protein synthesis, (**G**) interaction, and (**H**) PLA of 3F-YPEL2 and/or HA-SQSTM1 in transiently transfected COS7 cells were as assessed as described in legend of A-D. HDAC1 was used as a loading control.

Based on these results, we further characterized the interaction of YPEL2 with ELAVL1 or SQSTM1. In transiently transfected COS7 cells with expression vectors bearing the 3F-YPEL2 and/or HA-ELAVL1 cDNA, YPEL2 alone shows diffuse staining encompassing both the nucleus and cytoplasm (Fig. 7A). Although HA-ELAVL1 localizes primarily to the nucleus^103^, we also observe the localization of HA-ELAVL1 in the cytoplasm with intensely stained cytoplasmic foci wherein the 3F-YPEL2 staining shows overlaps. This suggests that ELAVL1 and YPEL2 co-localize to cytoplasmic structures, reminiscing of PGs/SGs. HA-ELAVL1 in transfected cells shows discrete electrophoretic migration with an MM of about 37 kDa; whereas 3F-YPEL2 displays an electrophoretic species migrating at 17 kDa MM (Fig. 7B). Immunoprecipitation of cellular extracts of transiently transfected cells that co-synthesize 3F-YPEL2 and HA-ELAVL1 using the HA or the Flag antibody together with protein A and G magnetic beads followed by immunoblotting with the Flag or the HA antibody indicates the presence of both HA-ELAVL1 and 3F-YPEL2 in the immunoprecipitants (Fig. 7C). The co-localization or Co-IP results were not due to the nature of the tags as we observed 3F-ELAVL1 and HA-YPEL2 interact and co-localize to PGs/SGs-like structures when co-synthesized (data not shown). These results demonstrate that 3F-YPEL2 and ELAVL1 interact only in the cytoplasm. Since ELAVL1 is predominantly found in the nucleus but can translocate to the cytoplasm through phosphorylation of residues located in the hinge region of the protein^103^, our results suggest that YPEL2 may interact with the phosphorylated ELAVL1.

To further verify that ELAVL1 is an interacting protein partner of YPEL2, we carried out the proximity ligation assay (PLA) as we described previously^54^. PLA utilizes species-specific secondary antibodies conjugated with distinct DNA primers. A hybridization step followed by circular DNA amplification with fluorescent probes to the conjugated DNA primers allows the visualization of proximity spots by fluorescence microscopy^104^. In transiently transfected COS7 cells, we observed prominent fluorescence signals in the cytoplasm, only when cells co-synthesize HA-ELAVL1 and 3F-YPEL2 in the presence of both the HA and the Flag antibodies (Fig. 7D), as we observe virtually no fluorescence signal in the presence of a single antibody even when cells synthesize both interacting partners (Supplemental Information, Fig. S11). These results lend credence to the conclusion that ELAVL1 is an interacting partner of YPEL2 when both proteins are present in the cytoplasm. Similar studies by the use of IP, WB, ICC, and PLA in transient transfection into COS7 cells with expression vectors bearing the 3F-YPEL2 and/or HA-SQSTM1 cDNA indicated that SQSTM1 is also an interaction partner of YPEL2 (Fig. 7E-H).

### Localization of the endogenous YPEL proteins in Stress Granules in response to Sodium Arsenite

Interaction of YPEL2 with ELAVL1, or SQSTM1, and co-localizations in PB/SG-like structures in the cytoplasm suggest that YPEL2 is involved in processes associated with the formation and/or maintenance of PBs/SGs thereby cellular stress responses. Because overexpression in experimental systems could induce SG formation without stress stimuli^105, 106^, to ensure that our results are not experimental artifacts, we examined the involvement of the endogenous YPEL in SG formation in cellular responses to a stress inducer. Since cells respond to various internal and external stresses including heat shock, amino acid deprivation, hypoxia, and oxidative stress by inducing the formation of SGs^63^ and since our Phyre2 results assign possible RNA binding and oxidoreductase activities to YPEL proteins (Fig. 1C), we chose sodium arsenite (SA) as an oxidative stress inducer to instigate SG formation^65, 107–109^.

To assess SGs formation in COS7 cells in response to SA, we treated cells without (- SA) or with 200 µM SA (+ SA), which was based on the minimal concentration of SA to induce SG-like loci in all cells in 1h (Supplemental Information, Fig. S3). Cells were then subjected to RNA-FISH with an ATTO488-conjugated oligo-(dT)_50_ probe for mRNA detection and/or RNA-FISH coupled ICC using a rabbit polyclonal antibody specific to G3BP1 as an essential nucleator of SG assembly to ensure that SA treatment of COS7 cells faithfully reconstitutes SG formation. We observed that oligo-(dT)_50_ and G3BP1 signals merge to cytoplasmic foci only in response to SA, indicating that SA-induced cytoplasmic foci are *bona fide* SGs (Fig. 8A).

**Fig. 8.**
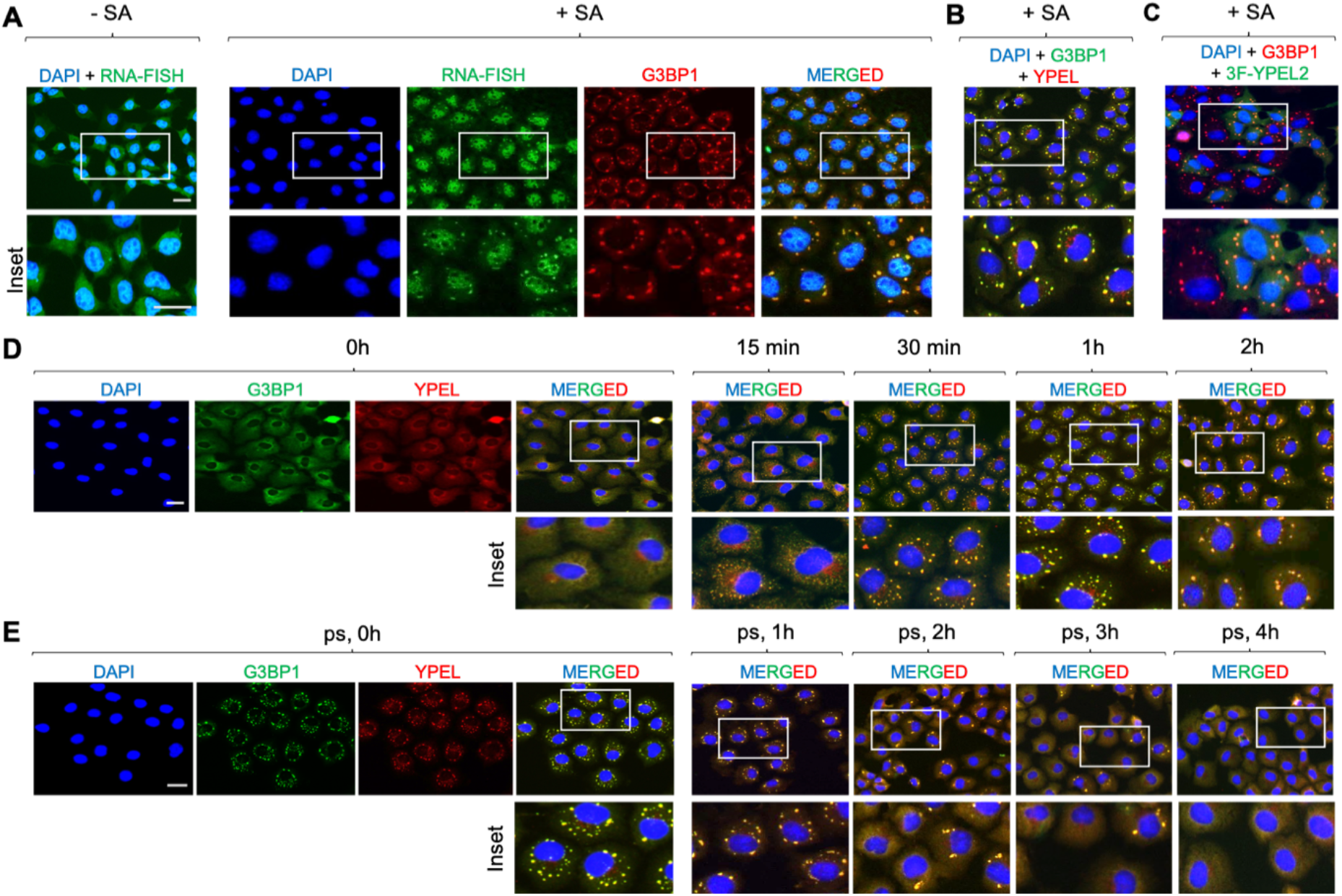
Stress granule formation in COS7 cells in response to sodium arsenite. (**A**) To assess the formation of stress granules, COS7 cells grown on coverslips were treated without (-SA) or with 200 µM sodium arsenite (+SA) for 1 h. Cells were fixed and subjected to hybridization-couple ICC using an ATTO488 conjugated oligo-dT probe and the polyclonal G3BP1 antibody (ab200550) followed by an Alexa Fluor-647 conjugated secondary antibody. DAPI indicates the nucleus. The scale bar is 20 µm. (**B**) To assess the co-localization of the endogenous YPEL proteins (YPEL) with G3BP1, cells treated with SA for 1 h were fixed and subjected to ICC using the monoclonal G3BP1 antibody (SCBT, sc-365338) followed by an Alexa Fluor-488 conjugated secondary antibody (405319, Biolegend) and the pan-YPEL antibody (SCBT, sc99727) followed by an Alexa Fluor-647 conjugated goat-rabbit secondary antibody (Abcam, ab150083). DAPI was used to indicate the nucleus. (**C**) To assess whether 3F-YPEL2 also co-localizes with G3BP1 in SGs, cells transfected with the expression vector pcDNA3.1(-) bearing 3F-YPEL2 cDNA were subjected to ICC using the polyclonal G3BP1 antibody (Abcam, ab200550) followed by an Alexa Fluor-647 conjugated secondary antibody and the monoclonal Flag antibody (Sigma-Aldrich, F1804) followed by an Alexa Fluor 488-conjugated secondary antibody. DAPI indicates the nucleus. (**D**) To examine the co-localization of YPEL and G3BP1 during the formation and disassembly of SGs, COS7 cells grown on coverslips were treated with 200 µM sodium arsenite (+SA) for 0, 15 min, 30 min 1 h, and 2 h. (**E**) Following 1 h of 200 µM sodium arsenite (+SA) treatment, cells (post-stress, PS), were washed and incubated in the growth medium without SA for 0, 1, 2, 3, or 4 h. (**D & E**) Fixed cells were then subjected to ICC using the monoclonal G3BP1 antibody (sc-365338) and the pan-YPEL antibody (sc99727) followed by secondary antibodies for visualization with a fluorescence microscope. DAPI staining indicates the nucleus. The scale bar is 20 µm.

Under identical conditions, co-localization of endogenous YPEL, assessed with the pan-YPEL antibody, with G3BP1 probed with a mouse monoclonal antibody specific to G3BP1 indicated that the endogenous YPEL proteins (YPEL) also localize to SGs (Fig. 8B). In transiently transfected COS7 cells, co-localization of 3F-YPEL2 assessed with the Flag antibody with the endogenous G3BP1 in cytoplasmic foci in response to 200 µM SA shows that 3F-YPEL2 as the endogenous YPEL is involved in SG-associated processes as well (Fig. 8C).

SG assembly and disassembly occur dynamically through sequential multistep processes that involve an initial nucleation event which follows a growth phase requiring the accumulation of substrate molecules and finally the fusion of the small initial stress granules into larger assemblies^65^. To examine whether or not the endogenous YPEL is also involved in dynamic SG assembly and/or disassembly of SGs, we treated COS7 cells with 200 µM SA for 15 min intervals up to 2h. Cells were then probed with the pan-YPEL and G3BP1 antibodies (Fig. 8D). Results revealed that 15 min after SA treatment, the endogenous YPEL and G3BP1 co-localized to numerous and various sizes SGs. These SGs coalesced to form larger size SGs thereby decreasing the number of SGs at later time points up to 2h within which YPEL and G3BP1 co-localized. Following a 1h SA treatment (post-SA, PS, treatment), SGs completely disappeared within 4h of the SA withdrawal leading to diffuse staining of both YPEL and G3BP1 (Fig. 8E). These results suggest that the YPEL protein is associated with the dynamic assembly and disassembly processes of SGs.

HA-G3BP1 did not interact (Supplemental Information, Fig. S10) but co-localized with 3F-YPEL2 in SG-like structures in the cytoplasm of transiently transfected COS7 cells (Fig. 8B, D & E). On the other hand, HA-ELAVL1 interacted and co-localized with 3F-YPEL2 in SG-like structures (Fig. 7). This raises the possibility that the interaction between ELAVL1 and YPEL is critical for the localization of the endogenous YPEL in SGs. To examine whether or not reducing levels of the endogenous ELAVL1 protein decreases/represses the localization of YPEL in SGs, we transiently transfected COS7 with AllStar control siRNA (AS-siR) or a siRNA pool that targets ELAVL1, EL1-siR, for 24 h. WB analysis indicated that EL1-siR effectively reduces both cytoplasmic and nuclear levels of ELAVL1 (Fig. 9A) reflected in ICC as well (Fig. 9B). To examine the SG formation, COS7 cells transfected with EL1-siR were treated without (Fig. 9C) or with 200 µM SA (Fig. 9D) for 1 h. We observed that even in the presence of reduced cytoplasmic levels of ELAVL1 in cells transfected with EL1-siR, YPEL localizes to SG-like structures in response SA (Fig. 9D) as observed in cells transfected with AS-siR (Fig. 9B). This suggests that YPEL localization to SGs is independent of ELAVL1. Furthermore, we observed that G3BP1 and YPEL co-localize to SGs with smaller sizes and substantially higher numbers (Fig. 9E) in cells transfected with EL1-siR compared to those in cells transfected with the control AS-siR, as reported previously^110^. This implies that ELAVL1 is a critical substrate for the maturation of SGs as shown previously^98, 99^.

**Fig. 9.**
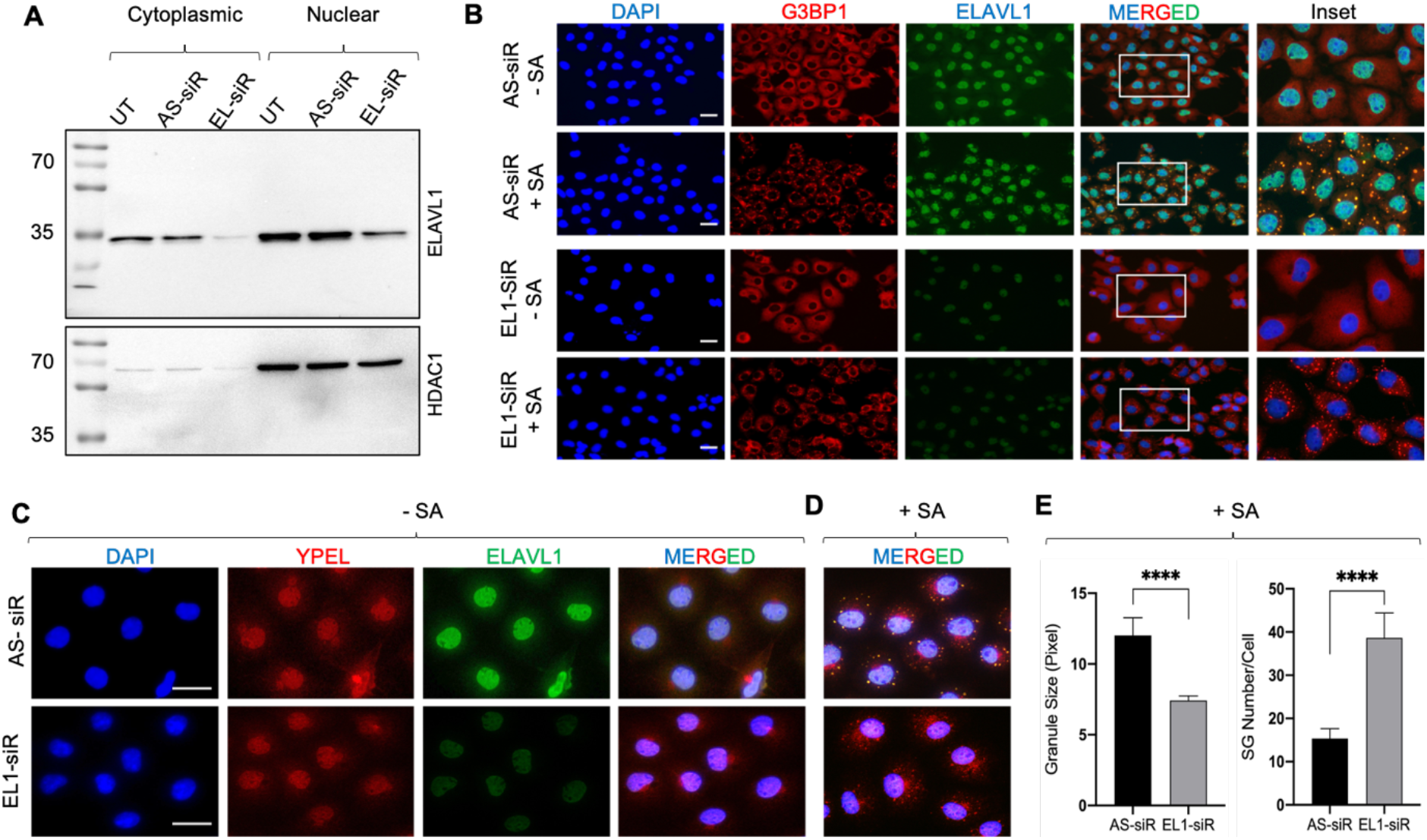
Effects of suppressed levels of ELAVL1 by a siRNA approach on the co-localization of YPEL and G3BP1 in SGs. (**A**) To examine whether or not a reduction of ELAVL1 levels in cells alters the location of YPEL in SGs, COS7 cells were left un-transfected (UT) or transiently transfected with a control siRNA AllStar (AS-siR) or a siRNA pool specific to ELAVL1 transcripts (EL-siR) for 24 h. Cytoplasmic and nuclear extracts (50 µg) were subjected to SDS-10%PAGE for WB analysis using a monoclonal antibody specific to ELAVL1 (SCBT, sc-5261). HDAC1 probed with the HDAC1-specific antibody (Abcam, ab19845) was used as the loading control. (**B**) Transfected COS7 cells with the AllStar siRNA or the ELAVL1 siRNA pool for 24 h were also treated without (-SA) or with 200 µM sodium arsenite (+SA) for 1 h. Cells were then fixed and subjected to ICC using the ELAVL1 antibody or the pan-YPEL antibody. DAPI staining indicates the nucleus. The scale bar is 20 µm. (**C & D**) To assess the co-localization of ELAVL1 and YPEL in SGs when ELAVL1 protein levels were reduced, COS7 cells transfected with (**C**) the control (AS-siR) or with (**D**) the siRNA pool specific to ELAVL1 transcripts (EL-siR) for 24 h were treated without (-SA) or with 200 µM sodium arsenite (+SA) for 1 h and subjected to ICC with the ELAVL1-specific monoclonal antibody and the pan-YPEL antibody. DAPI staining indicates the nucleus. The scale bar is 20 µm. (**E**) To assess the effect of the reduced levels of ELAVL1 on the size and numbers of SG, cells shown in Fig. D were subjected to quantification using the CellProfiler image analysis program. Asterisks indicate significant changes (P< 0.0001).

## DISCUSSION

We here initially explored the identification of dynamic YPEL2 protein interaction networks as a means to delineate the functional features of YPEL2 in mediating cellular processes using inducible YPEL2 synthesis in COS7 cells under steady-state conditions followed by dynamic TurboID-MS analyses. Our results indicate that the proximity interaction partners of YPEL2 encompass proteins largely involved in RNA metabolic processes including RNA folding, mRNA maturation, stability, export, translation, and degradation. Since the proximity interacting partners of YPEL2 also functionally overlap in processes associated with the formation and disassembly of cytoplasmic RNP granules involved in cellular stress responses, as we observe here with sodium arsenite an oxidative stress inducer, we suggest that YPEL2 is involved in stress surveillance mechanisms critical for cellular survival.

Our *in silico* analyses show that YPEL proteins with MMs ranging from 13.5 to 17.5 possess only a single structural feature: the Yippee domain. Homology modeling, as an attempt to infer a function(s) for YPEL proteins through the Yippee domain, known also as MeDIY^111^ or CULT/β-tent fold domain^112^, indicated matches for proteins with oxidoreductase, RNA binding/hydrolase, and chaperon functions in several protein families of prokaryotes and eukaryotes. The presence of yippee-like globular domains in oxidoreductase MSRA/MSRB proteins of prokaryotes and eukaryotes, E3 ligase cereblon protein, RNA helicases DHR-3, RIG-I/DHX58, and MDA5 proteins, as well as chaperon MSS4/RABIF protein of eukaryotes, likely indicate an evolutionary relationship, as suggested^77, 112^.

However, distinct ligand specificities of the evolutionarily conserved yippee domain in these proteins rendered predictions of mechanisms involved in ligand recognition hence assigning a physiological ligand(s) for YPEL2 in particular and YPEL proteins in general difficult. Despite the Yippee fold, differences in amino acid compositions of the Yippee fold in proteins could generate specialized molecular interaction networks that drive the formation of discrete interacting surfaces resulting in selective ligand recognition. For example, the deposition and restoration level of CENP-A at centromeres is primarily mediated by the MIS18 complex composed of MIS18Α, MIS18Β, and MIS18BP1^78^. Structural studies indicate that the heterodimerization and subsequent formation of the oligomeric state of MIS18Α and MIS18Β are initiated by the Yippee domains through specific residues in the dimerization interfaces. The Yippee domain of MIS18Α in the oligomer then specifically recruits two monomers of MIS18BP1 to form the MIS18 complex, whereas the Yippee domain of MIS18Β through a distinct ligand interacting surface directly associates with CENP-C for subsequent CENP-A deposition^111, 113^. Similarly, members of the RIG-I-like RNA helicase (RLR) family including DRH3 of *C. elegans*, the human RIG-I/DHX58, and MDA5 proteins are involved in the endogenous RNAi biogenesis as well as in recognition of cytoplasmic viral RNA as sensors. This family of proteins possesses a central helicase domain and a carboxyl-terminally located Yippee domain (also referred to as the regulatory domain, RD) with different RNA binding specificities^114^ due to the protein-specific regional conformation^115^. DRH3, for example, binds both single-stranded and double-stranded RNAs independently of 5’-triphosphate^116^, whereas the Yippee domain of MDA5 preferentially binds long, capped 5’ mono- or di-phosphate containing double- or single-stranded RNAs. On the other hand, the Yippee domain of RIG-I associates with short double-stranded or 5’-triphosphates of uncapped single- or double-stranded RNAs^114, 117, 118^. Interestingly, as one of the YPEL proximity interactors ZCCHC3 acts as a co-receptor for both RIG-I and MDA5 by binding to RNA and interacting with the Yippee domains of RIG-I and MDA5, thereby enhancing RNA binding of these proteins^119^. Moreover, the recruitment of E3 ligase TRIM25 to the RIG-I/MDA5/ZCCHC3 complex by ZCCHC3 induces the ubiquitylation and activation of RIG-I and MDA5^119^.

In contrast to the substrate specificities of modular RLR proteins, proximal interaction partners of YPEL2 possessing only the Yippee domain show remarkable diversity. We observed under steady-state conditions in COS7 cells that YPEL2, as the endogenous YPEL protein(s), shows diffuse intracellular staining encompassing both the nucleus and cytoplasm and interacts with functionally diverse proteins involved in cellular processes ranging from RNA metabolism to cellular proliferation. Importantly, a substantial portion of the interaction partners of YPEL2 at all time points coincided with proteins critical for the formation/resolution of PB and SG, as indicated by interactions and co-localizations of YPEL2 with RNA binding proteins including ELAVL1 under both the steady-state and oxidative stress condition in response to SA. These observations suggest that YPEL2 can temporarily associate with functionally integrated and spatially restricted multicomponent complexes involved in various cellular processes and raise the possibility that YPEL2 uses a common feature to recognize and interact with the putative partners.

One of the predicted functional features for YPEL by our homology modeling involves oxidoreductase activity derived from the evolutionary conserved MSRA and MSRB proteins of eukaryotes^120–122^. Under steady-state conditions, reactive oxygen species (ROS), including superoxide anion (O_2_•−), hydroxyl radical (•OH), singlet oxygen (^1^O_2_), hydrogen peroxide (H_2_O_2_), and peroxyl radicals (ROO•), are formed as physiological metabolic by-products of reduction-oxidation reactions mediated by a large number of oxidase enzymes and the mitochondrial electron transport chain^120–122^. Effects of ROS in cells are largely manifested in the reversible oxidation of methionine, cysteine, selenoproteins, lipids, and RNA/DNA leading to modification of substrate activities^120–122^. The formation and subsequent degradation of ROS are regulated by cellular defense systems, including superoxide dismutases, catalase, thioredoxin/thioredoxin reductase, and glutathione peroxidase as scavenging enzymes capable of removing oxidants or their precursors^120–122^. Non-enzymatic antioxidants such as tocopherols and ascorbic acid, and also metal binding proteins delay oxidation reactions or prevent the development of reactive species. Repair and removal systems including methionine sulfoxide reductases, disulfide reductases/isomerases, and the ubiquitin-proteasome-system are critical for counteracting the effects of ROS^120–122^. Increased levels of ROS that exceed the ability of the cell defense system to remove oxidants result in the activation of intracellular signaling pathways for removal of, or adaptation to, stress or stress-induced cell death^120, 121^. Methionine, for example, in proteins acts as a reversible redox switch which constitutes an important antioxidant defense mechanism that affects many cellular functions. Methionine can be oxidized to methionine sulfoxide either non-specifically by ROS or as a specific, enzyme-catalyzed post-translational modification. On oxidation, the methionine sulfur atom becomes a new chiral center, producing methionine-R-sulfoxide or methionine-S-sulfoxide altering the local conformation of proteins^120, 121^. While the MSRA enzyme present throughout the cell reduces the S-epimer, MSRB enzymes (MSRB1, MSRB2, MSRB3A, and MSRB3B) with distinct intracellular distributions reduce the R-epimer to generate unmodified methionine. Of MSRBs, MSRB1 exhibits the highest methionine-R-sulfoxide reductase activity because of the presence of selenocysteine (Sec) in its active site^70, 123, 124^. For example, control of the assembly and disassembly of actin affecting various cellular events ranging from cell morphology to proliferation is shown to be mediated by the reversible oxidation of methionine residues by the MICAL family monooxygenases and subsequent reduction of methionine sulfoxides by MSRB1^125^. Similarly, Ca^++^/calmodulin-dependent protein kinase II (CaMKII) is involved in many signaling cascades critical for energy metabolism and adaptation to oxidative stress^126^. Calcium-independent activation of CaMKII through the oxidation of methionine residues leads to apoptosis, while MSRA reverses the autoactivation of CaMKII thereby inhibiting apoptosis^127^. Despite the Yippee fold, it is unlikely that YPEL proteins are members of the methionine-sulfoxide reductase family as they lack the invariably conserved cysteine-containing motifs essential for the catalytic activity of MSRA (GCPWG) or the MSRB (RXCXN) proteins^70, 123, 124^. Nevertheless, the yippee domain of YPELs as a ‘reader/sensor’ could associate with oxidized methionine in free form or in proteins.

The possible ‘reader/sensor’ function of YPELs could also be operational for the oxidation of cysteine residues of cellular proteins. As methionine, cysteine is a redox-sensitive site that can be covalently modified by ROS through reversible and irreversible oxidation by sensing levels of oxidative stressors^128, 129^. Due to the unique chemical characteristics of its thiol group, cysteine plays also a crucial role in the structure of proteins and protein-folding pathways through intra- and inter-molecular linkages with other cysteine residues. For example, cysteine oxidation of the SG-nucleating protein TIA1 is shown to lead to oligomerization and subsequent inhibition of SG assembly that result in the promotion of apoptosis^130^. This finding lends also credence to the conclusion that the removal of oxidation-induced damage on proteins is critical for cell survival. SQSTM1, one of the YPEL2 interacting partners, is a prototypical autophagic receptor that links ubiquitylated substrates to the nascent autophagic vesicles for cell survival^131^. Studies indicate that oxidation of cysteine residues in SQSTM1 in response to hydrogen peroxide, H_2_O_2_, at low concentrations triggers rapid but transient SQSTM1 oligomerization, in contrast to high concentrations that lead to aggregation and subsequent degradation of SQSTM1 with its ubiquitinated substrates through autophagy^132^. Moreover, TARDBP as one of the YPEL2 proximity interactors acts also as an RNA-binding protein. Cysteine oxidation of one of the RNA recognition motifs of TARDBP is shown to result in the loss of function by fragmentation and accumulation of fragments in insoluble aggregates^133^. Furthermore, BAG6 is a member of the Bcl-2 associated athanogene family which is an evolutionarily conserved, multifunctional group of co-chaperone regulators including BAG2 and BAG3, which are YPEL2 proximal interactors. BAG6 functions in the prevention of the aggregation of misfolded hydrophobic patch-containing proteins as a part of a cytosolic protein quality control complex^134^. It appears that BAG6 as a sensor for fragments of TARDBP facilitates the ubiquitylation of fragments for degradation thereby preventing their intracellular aggregation^135^. The possible reader/sensor function for YPEL2 of redox homeostasis is supportive of our observation that YPEL2 interacts with the members of the RIG-I/MDA5/ZCCHC3/TRIM25 complex which counteracts viral infection-generated increases in ROS levels^136^.

The possible ‘reader/sensor’ function of YPELs could allow YPELs to sequester ROS. The Yippee domain of YPEL proteins contains two cysteine pairs that are fifty-two amino acids apart (CX_2_C-X_52_-CX_2_C). Our structural similarity analyses suggest that the Yippee fold of YPELs forms a zinc (Zn^++^) binding pocket (Supplemental Information, Fig. S12), as, for example, observed with the MSRB proteins^70, 137^, Zn^++^-mediated cysteine residue coordination is critical for the formation of a stable protein structure and conformation that mediates protein function, protein-DNA, protein-RNA, and protein-protein interactions^138, 139^. Resulting in the release of Zn^++^, the oxidation of cysteine thiols of the Yippee fold could sequester ROS. Subsequent formation of intra/inter-molecular linkages with other cysteine residues within the protein or other proteins could lead to protein aggregations critical for the removal, elimination, and/or repair of oxidized substrates under both steady-state and oxidative-stress-induced conditions to ensure cell survival, as observed with TIA1^130^ and SQSTM1^132^.

The probable reader/sensor function of YPEL2 for oxidized substrates could also be critical for processes in the repair of oxidative stress-mediated damages/modifications to RNA. In addition to proteins, ROS under both steady-state and oxidative stress conditions induce nucleic acid 8-oxo-guanine (o^8^G) modification that occurs more frequently on coding- and non-coding RNAs, including rRNA and tRNA, than DNA^140–143^. o^8^G modification affects every stage of mRNA metabolism including RNA folding, maturation processing, stability, export, translation, translation fidelity, and decay^140^–_143_. For instance, XRN1, one of the YPEL2 proximity partners, is a member of the 5′→3′-exoribonucleases family that plays key roles in mRNA processing and turnover, no-go and nonsense-mediated decay, as well as the RNA interference pathways^144^. It is shown that o^8^G modifications induce translation stalling^145^ and decrease the processing efficiency of XRN1^146^. In addition, ribosome rescue upon translation stalling is carried out by a set of proteins including ABCE1, a proximity interacting partner of YPEL2, which participates in the alleviation of stalling-induced translational stresses^147^.

In summary, our results suggest that YPEL2 likely as a sensor/reader for ROS is involved in stress surveillance critical for cellular proliferation through proximity interactions with protein networks largely encompassing RNA metabolic processes. Our findings provide a point of departure to delve further into the involvement of YPEL2 in cellular events and also testable predictions about the structure/function of YPEL2. Since ROS act as damaging molecules and are also critical inducers of cellular signaling networks, deciphering the potential function and role of YPEL2 in stress surveillance mechanisms could contribute to a better understanding of the physiology and pathophysiology of cellular stress responses.

## LIST OF SUPPLEMENTARY INFORMATION

Figs. S1 to S12

Table S1 and S2

## ACKNOWLEDGMENT

This work is supported by the TUBITAK-1001-117Z213 grant (MM) and TUBITAK-1002-114Z738 (MM). The Genotype-Tissue Expression (GTEx) Project was supported by the Common Fund of the Office of the Director of the National Institutes of Health, and by NCI, NHGRI, NHLBI, NIDA, NIMH, and NINDS. The data used for the analyses were obtained from the GTEx Portal on 10/22/2022. We express our gratitude to Gizem Güpür who initiated the project. We gratefully acknowledge the critical guidance of Dr. Nurhan Özlü and Büşra Akarlar of the Koç University Proteomics Facility, Istanbul, Türkiye, for the execution and analysis of mass spectrometry. We thank Dr. Nurcan Tunçbağ for her guidance and suggestions in bioinformatics analyses. We thank Dr. Deniz Kahraman for allowing us to access the fluorescence microscope. We thank Dr. Begüm Akman Tuncer for the critical reading of the manuscript. We also thank members of the Muyan laboratory for stimulating discussions, contributions, and critical reading of the manuscript.

## AUTHOR CONTRIBUTIONS

Gizem Turan: data curation, formal analysis, investigation, methodology, and writing-original draft. and editing. Çağla Ece Olgun: data curation, formal analysis, investigation, methodology, writing-original draft, and editing. Hazal Ayten: investigation and methodology.

Pelin Toker: investigation and methodology.

Annageldi Ashyralyyev: investigation and methodology.

Büşra Savaş: investigation and methodology.

Ezgi Karaca: formal analysis, methodology, investigation, and editing.

Mesut Muyan: conceptualization, data curation, formal analysis, supervision, funding acquisition, methodology, writing-original draft and editing, and project administration.

## SUPPLEMENTARY MATERIAL

### Supplementary Figures

**Fig. S1.**
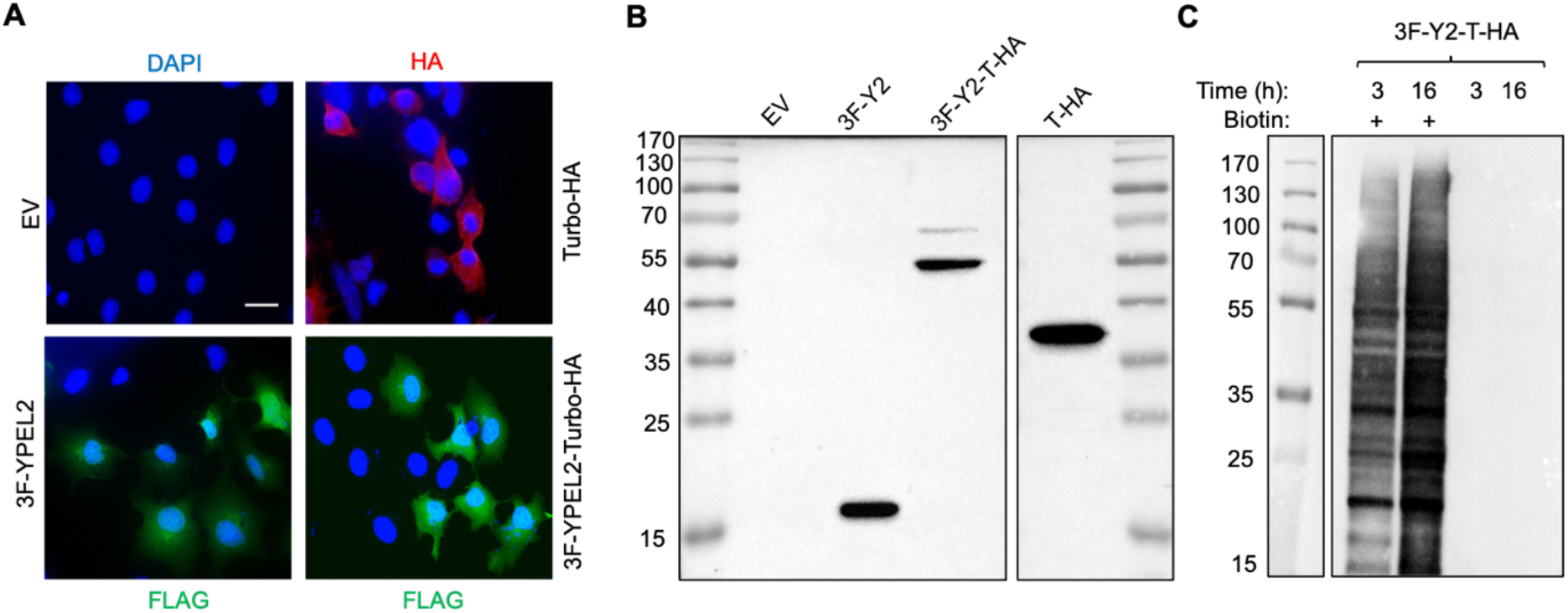
Effects of TurboID constructs on the biotinylation of intracellular proteins in COS7 cells. (**A & B**) COS7 cells were transfected with the pcDNA3.1(-) expression vector bearing none (EV), the 3F-YPEL2 (3F-Y2), 3F-YPEL2-Turbo-HA (3F-Y2-T-HA), or Turbo-HA (T-HA) cDNA. 24 hours after transfection, cells were subjected to (**A**) ICC or (**B**) WB using the Flag antibody (F-1804) for 3F-YPEL2 and 3F-YPEL2-Turbo-HA or the HA antibody (ab9119) for Turbo-HA. In ICC, DAPI staining was used to indicate the nucleus. The scale bar is 20 µm. For WB, molecular masses (MM) in kDa are indicated. (**C**) Assessing the duration of biotin incorporation into endogenous proteins. COS7 cells were transiently transfected with the pcDNA3.1(-) expression vector bearing 3F-YPEL2-Turbo-HA (3F-Y2-T-HA). Twenty-four hours after transfections, cells were incubated in fresh medium without or with (+) 50 μM biotin and 1 mM ATP for 3 or 16 h for the biotinylation of endogenous proteins. Cells were then collected, lysed, and equal amounts (50 µg) of protein extracts were subjected to SDS-10%PAGE electrophoresis followed by WB analysis using the Biotin (ab53494) antibody. Molecular masses (MM) are indicated in kDa.

**Fig. S2.**
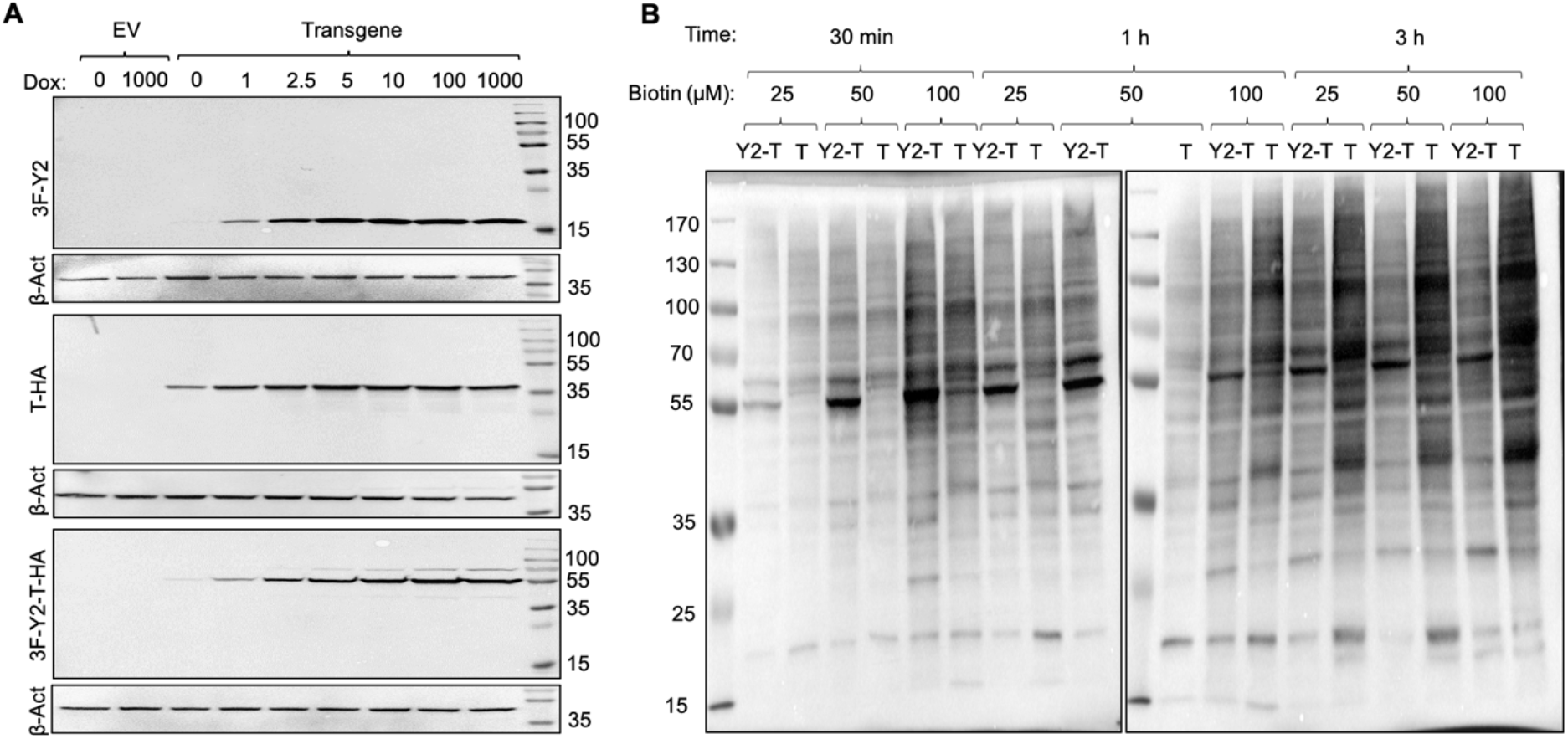
Inducible synthesis and effects of biotin concentrations on the biotinylation of intracellular proteins in COS7 cells by TurboID constructs. (**A**) COS7 cells were transfected with the pINDUCER20-MCS vector carrying none (EV), the 3F-YPEL2 (3F-Y2), 3F-YPEL2-Turbo-HA (3F-Y2-T-HA), or Turbo-HA (T-HA) cDNA. 24 hours after transfection, cells were treated without (0) or with various concentrations of Dox (1-1000 ng/ml) to induce protein synthesis for 24 h. Cells were then collected and equal amounts (50 µg) of protein extracts were subjected to SDS-10%PAGE electrophoresis followed by WB analyses using the Flag antibody (F-1804), for 3F-YPEL2 (3F-Y2) and 3F-YPEL2-Turbo-HA (3F-Y2-T-HA) or the HA antibody (ab9119) for Turbo-HA (T-HA). We also used β-actin (β-Act) as the loading control using the β-actin antibody (ab8227). Molecular masses (MM) are indicated in kDa. (**B**) To assess the kinetics of biotinylation of intracellular proteins with 3F-YPEL2-Turbo-HA or Turbo-HA (T-HA), cells transiently transfected with the pINDUCER20-MCS vector bearing 3F-YPEL2-Turbo-HA (3F-Y2-T-HA) or Turbo-HA (T-HA) cDNA for 24h were treated with 10 ng/ml Dox for the synthesis of transgene protein synthesis. Twenty-four hours after Dox induction, cells were treated with 25, 50, and 100 µM Biotin for 30 min, 1 h, or 3 h. Cells were collected, lysed, and total protein extracts (50 µg) were subjected to WB analysis using the Biotin antibody (ab53494).

**Fig. S3.**
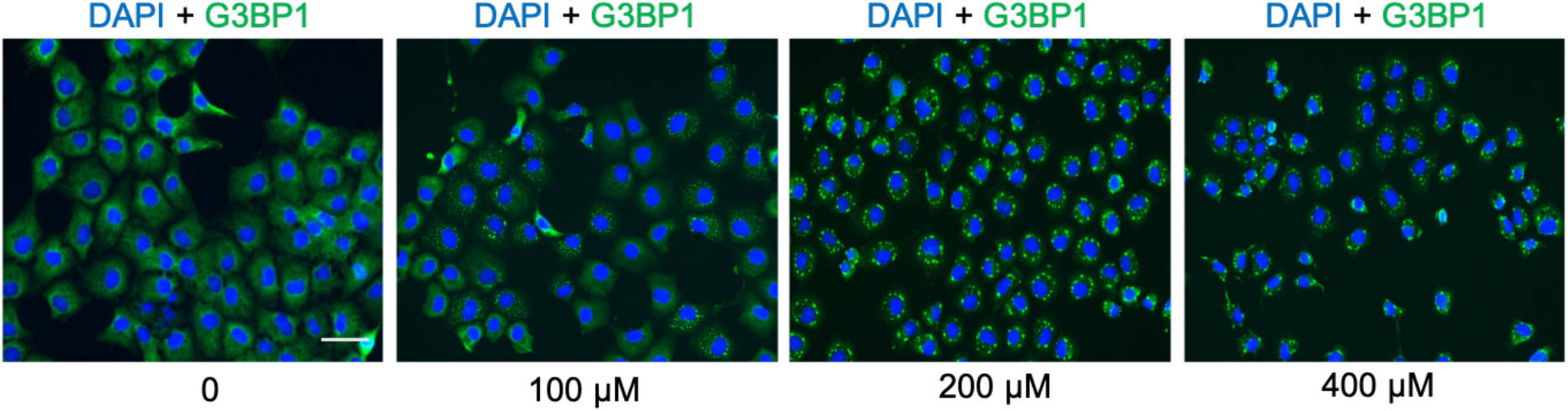
Effects of various concentrations of sodium arsenite (SA) on stress granule formation in COS7 cells. To assess the optimal concentration of SA, COS7 cells grown on coverslips were treated without (0) or with 100, 200, or 400 µM SA for 1h. Cells were then fixed, washed, permeabilized, and subjected to ICC using the monoclonal G3BP1 antibody (sc-365338) followed by an Alexa Fluor-488 conjugated secondary antibody (405319, Biolegend). DAPI staining was used to indicate the nucleus. The scale bar is 20 µm.

**Fig. S4.**
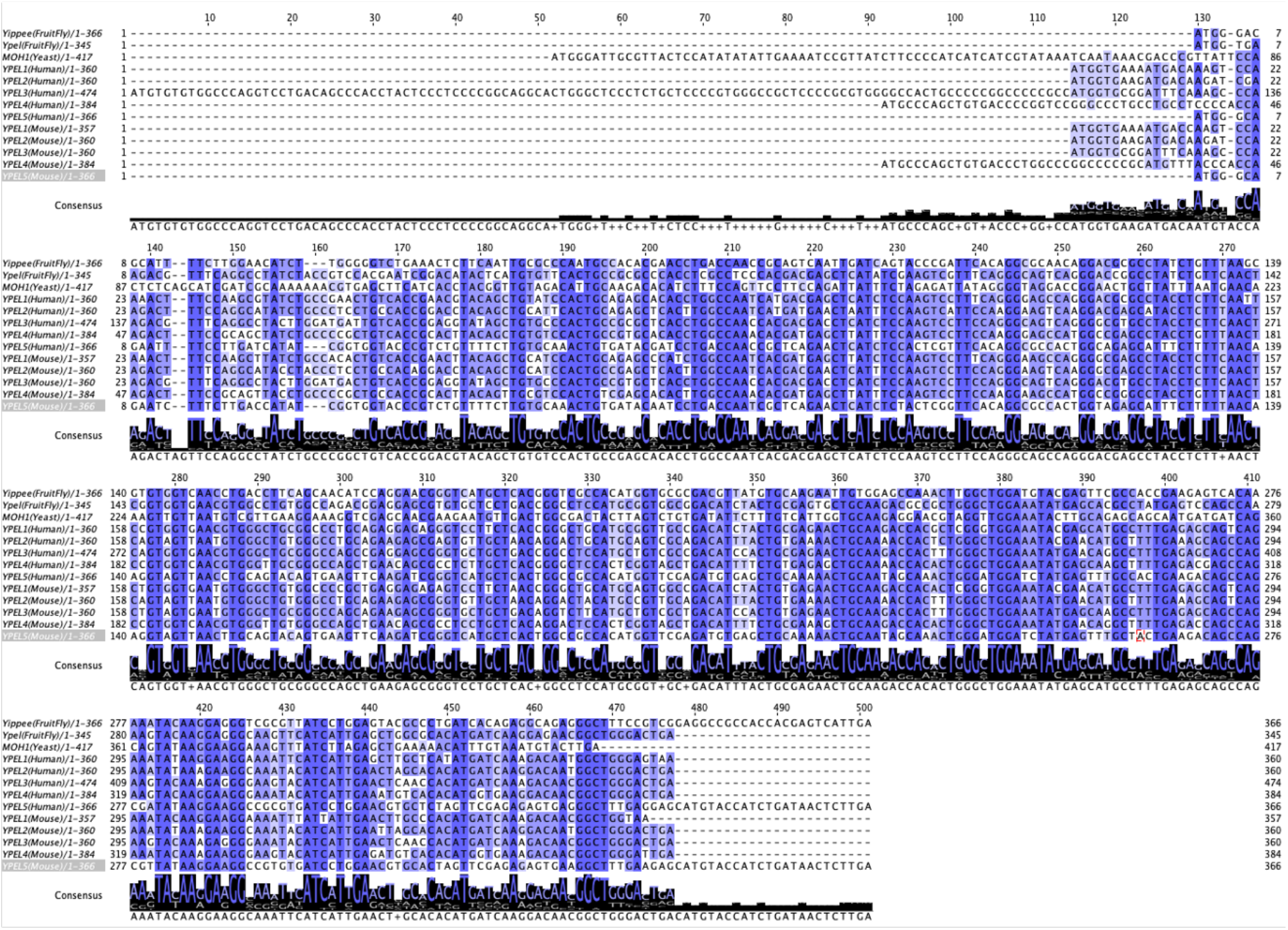
Alignment of ORF sequences of human YPEL genes in comparison with drosophila YIPPEE and yeast MOH1 genes. The alignment of ORF sequences of the *YPEL* family was generated with the Jalview program using the ClustalOmega plug-in.

**Fig. S5.**
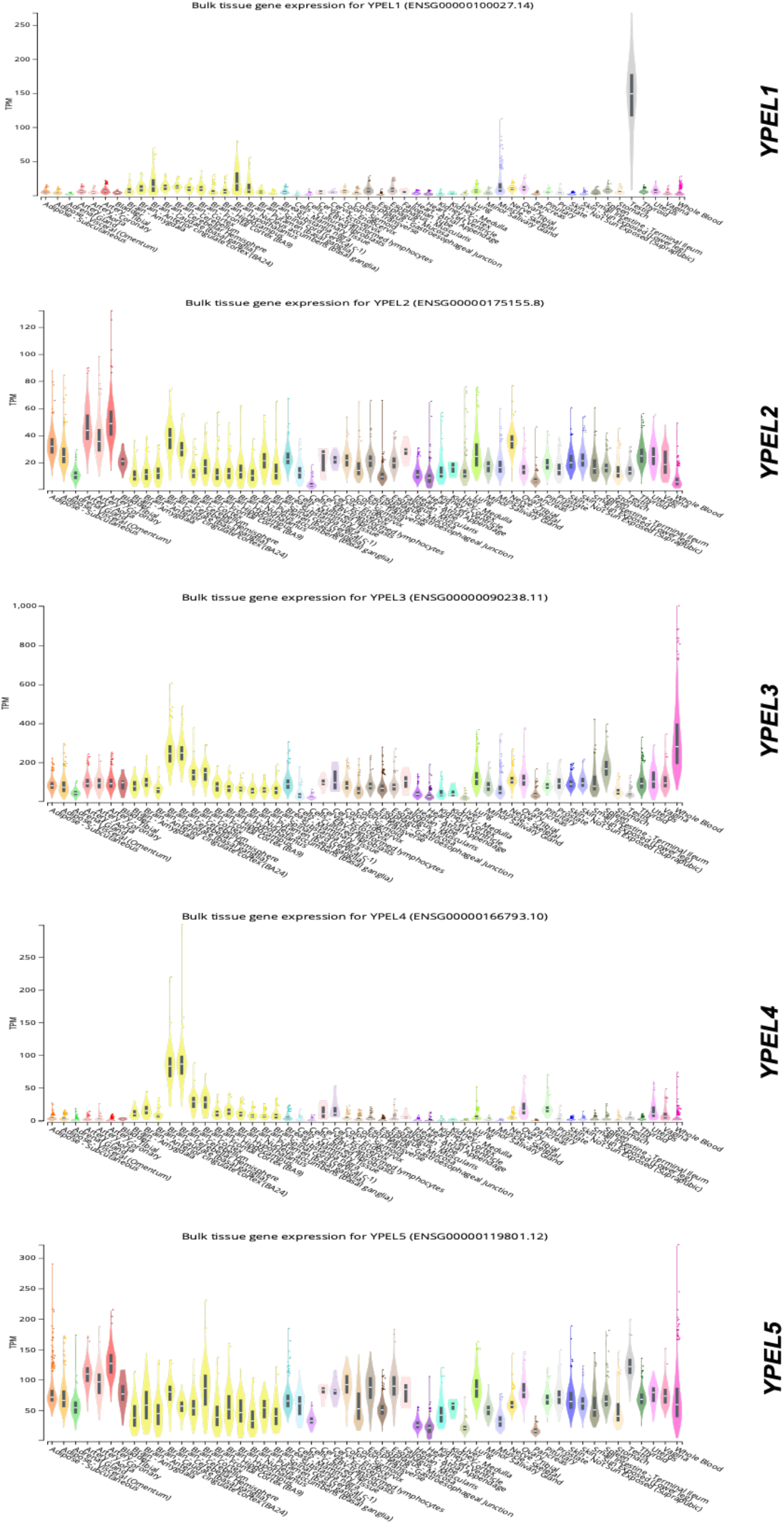
Expression of the YPEL family genes in human tissues. To assess the bulk tissue expressions of individual YPEL genes, we used the GTEx portal (https://gtexportal.org/home/).

**Fig. S6.**
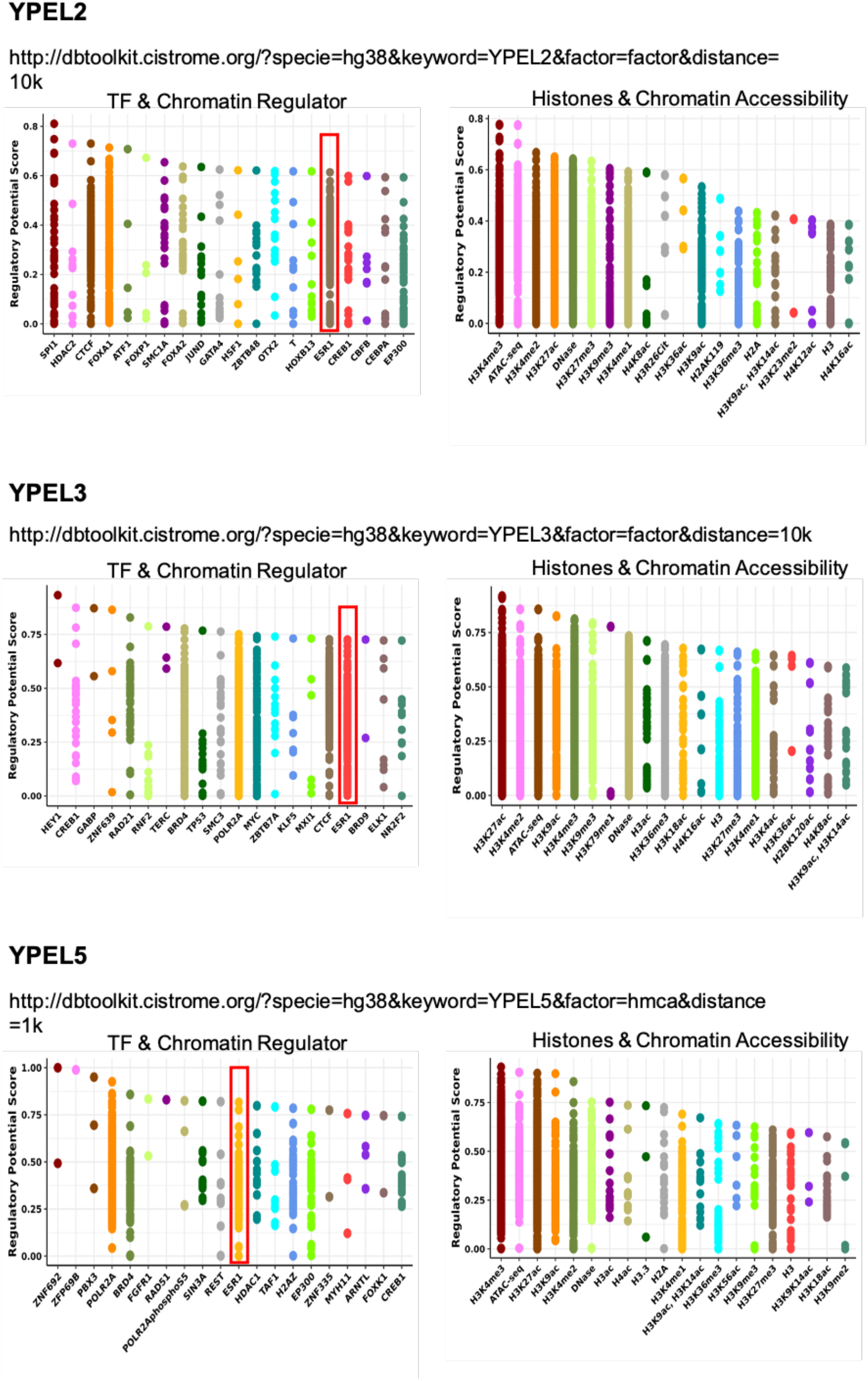
*In silico* analyses of chromatin structures and cis-regulatory elements of YPEL2, YPEL3, and YPEL5 genes. Analyses with the use of the Genome Browser (http://genome.ucsc.edu) and the Cistrome portal (http://dbtoolkit.cistrome.org/) for chromatin structures and cis-regulatory elements within a ±10 kb distance to TSS of putative promoters suggest that these promoters are located in, or vicinity of, CpG islands displaying accessible chromatin structures, and are associated with transcription regulators including ERα (red rectangular), BRD4, CTCF, FOXA1.

**Fig. S7.**
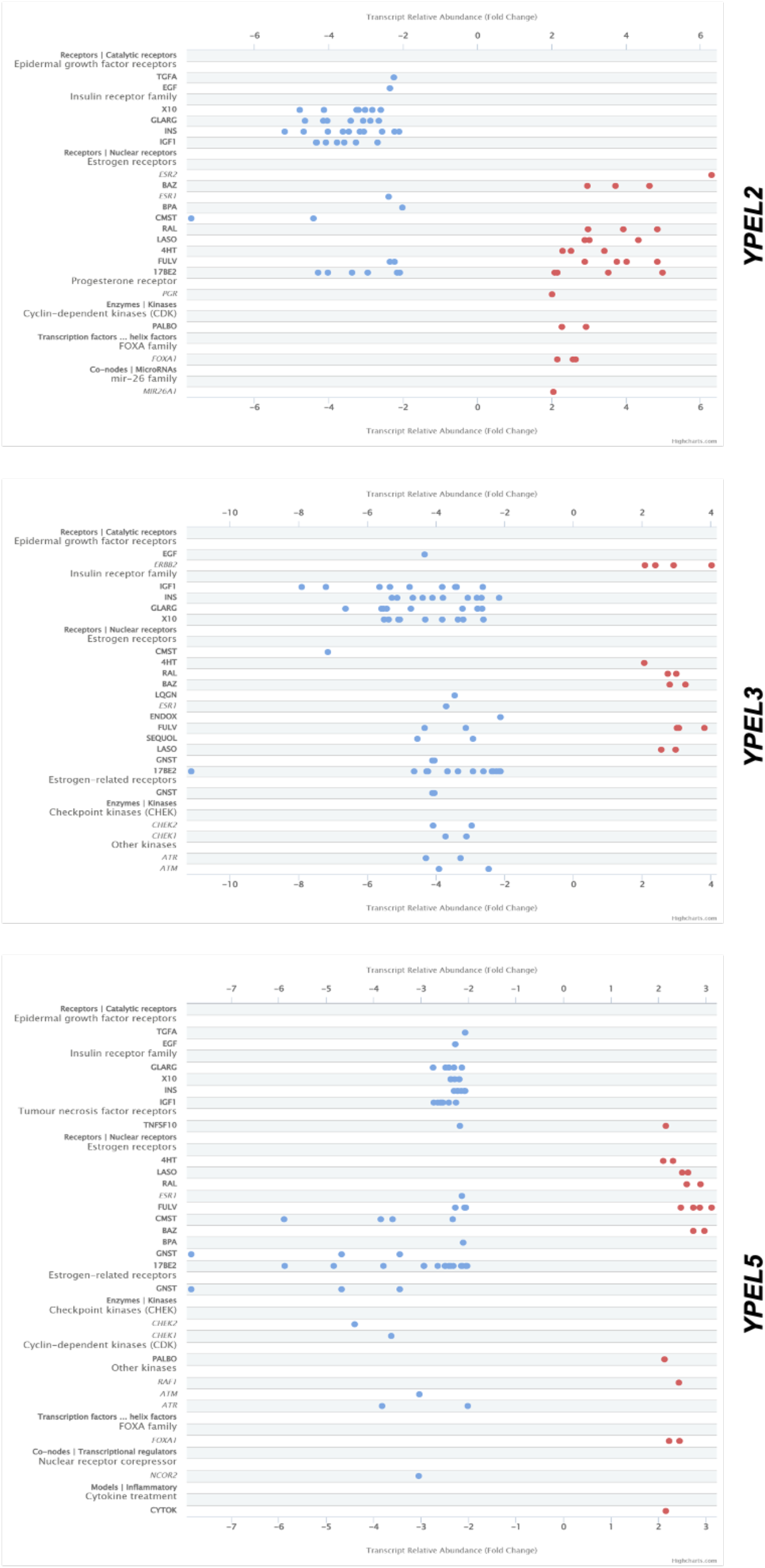
Regulation of *YPEL2*, *YPEL3,* and *YPEL5* by common signaling pathways. Analyses of the human mammary gland datasets with the Signaling Pathways Project^148, 149^ portal (https://www.signalingpathways.org/) suggest that EGF, the insulin family or estrogen signaling is involved in *YPEL2, YPEL3*, and *YPEL5* expressions.

**Fig. S8.**
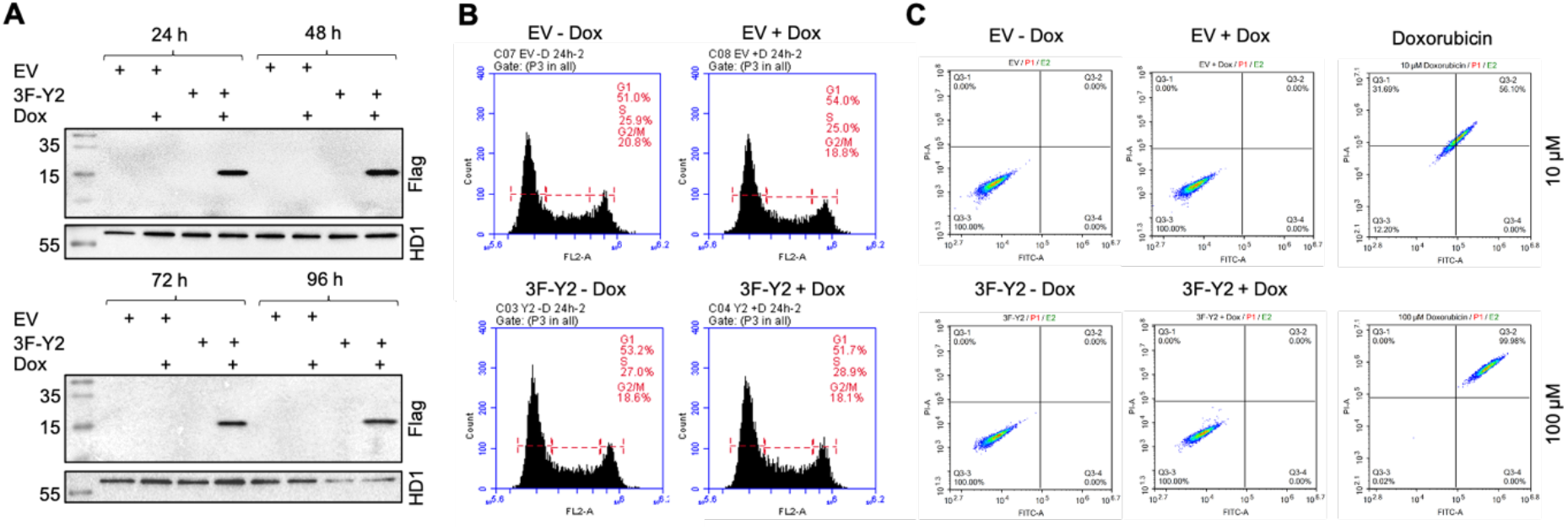
Assessing effects of 3F-YPEL2 on the growth of COS7 cells. (**A**) COS7 cells were transiently transfected with the pINDUCER20-MCS vector bearing none (EV) or the 3F-YPEL2 cDNA for 24 h. Cells were then treated without or with 10 ng/ml of Dox every 24 h for 96 h. Total cell extracts (50 µg) were then subjected to WB using the Flag antibody. Membranes were re-probed with the HDAC1 antibody. Molecular masses (MM) in kDa are indicated. (**B**) Transfected cells treated without or with 10 ng/ml of Dox for 24 h were also subjected to cell cycle analysis and (**C**) Annexin V staining. (**D**) Un-transfected cells were also treated with 10 or 100 µM apoptosis inducer Doxorubicin for 24h as control.

**Fig. S9.**
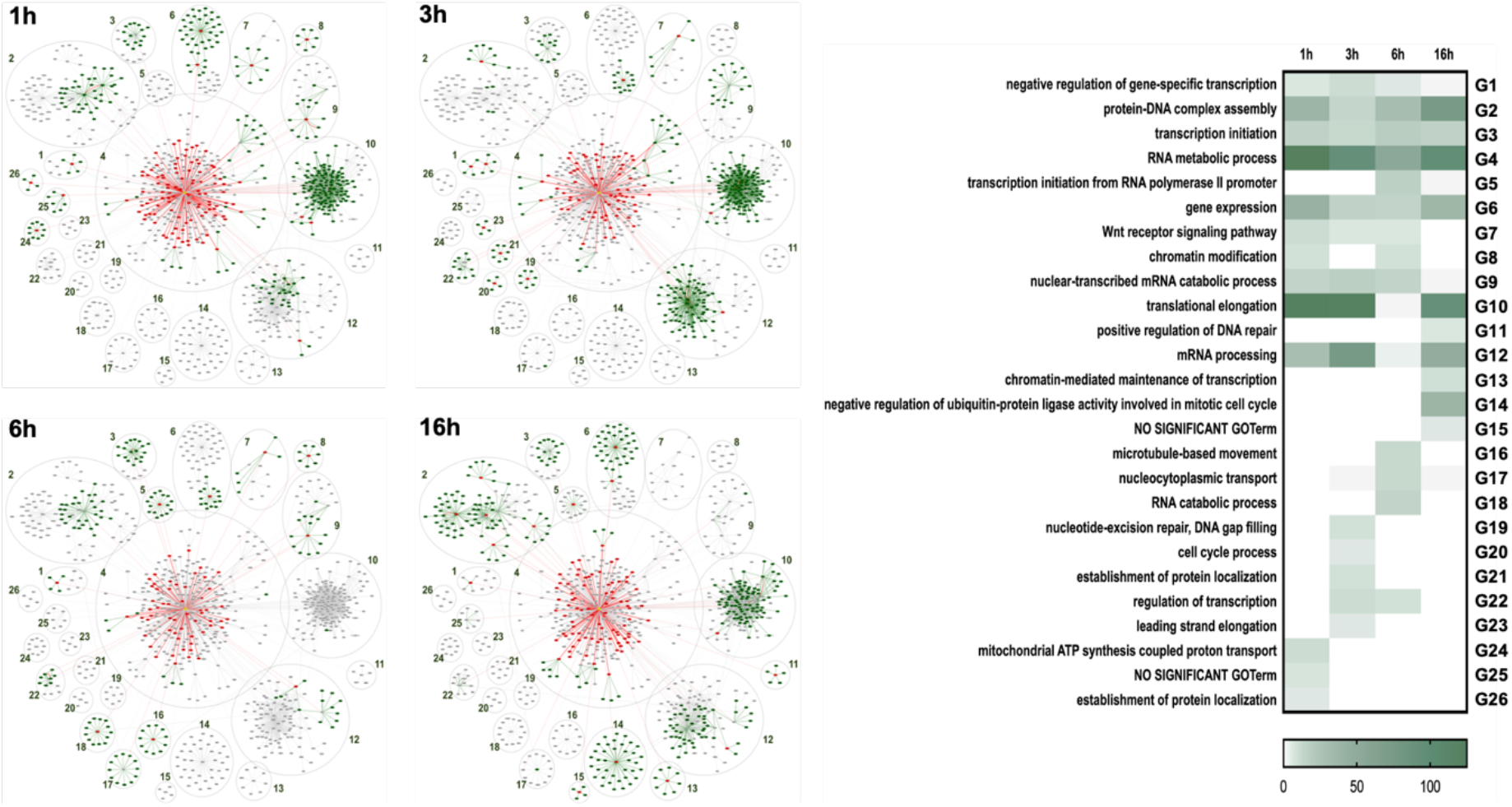
Secondary neighbor analyses of the proximity interaction partners of YPEL2. Proteins identified as primary interactors of YPEL2 were subjected to secondary neighbor analysis using the STRING database. Proteins in the human interactome with curated database or experimental scores above 900 that were associated with at least two of the primary interactors of YPEL2 were defined as secondary neighbors. For the visualization of the protein network including primary and secondary neighbors, the Cytoscape^150^ (http://www.cytoscape.org/) network visualization and analysis tool was used. When visualizing the dynamic protein network of YPEL2, the primary neighbors were colored in red, the secondary neighbors were colored in green and the proteins not seen at that time interval were indicated as gray for each experimental hour (1, 3, 6, and 16 h). For protein clustering, we used the CommunityClustering (Glay) option of the clusterMaker tool^151^ available as a plug-in in Cytoscape. After clustering, the GOTerm Biological Process annotations of each cluster, with the threshold value of 0.05, were assessed with the BINGO and the Benjamin and Hochberg False Discovery Rate correction of Cytoscape. The heatmap was constructed according to the number of genes enriched in GOTerms by using GraphPad Prism.

**Fig. S10.**
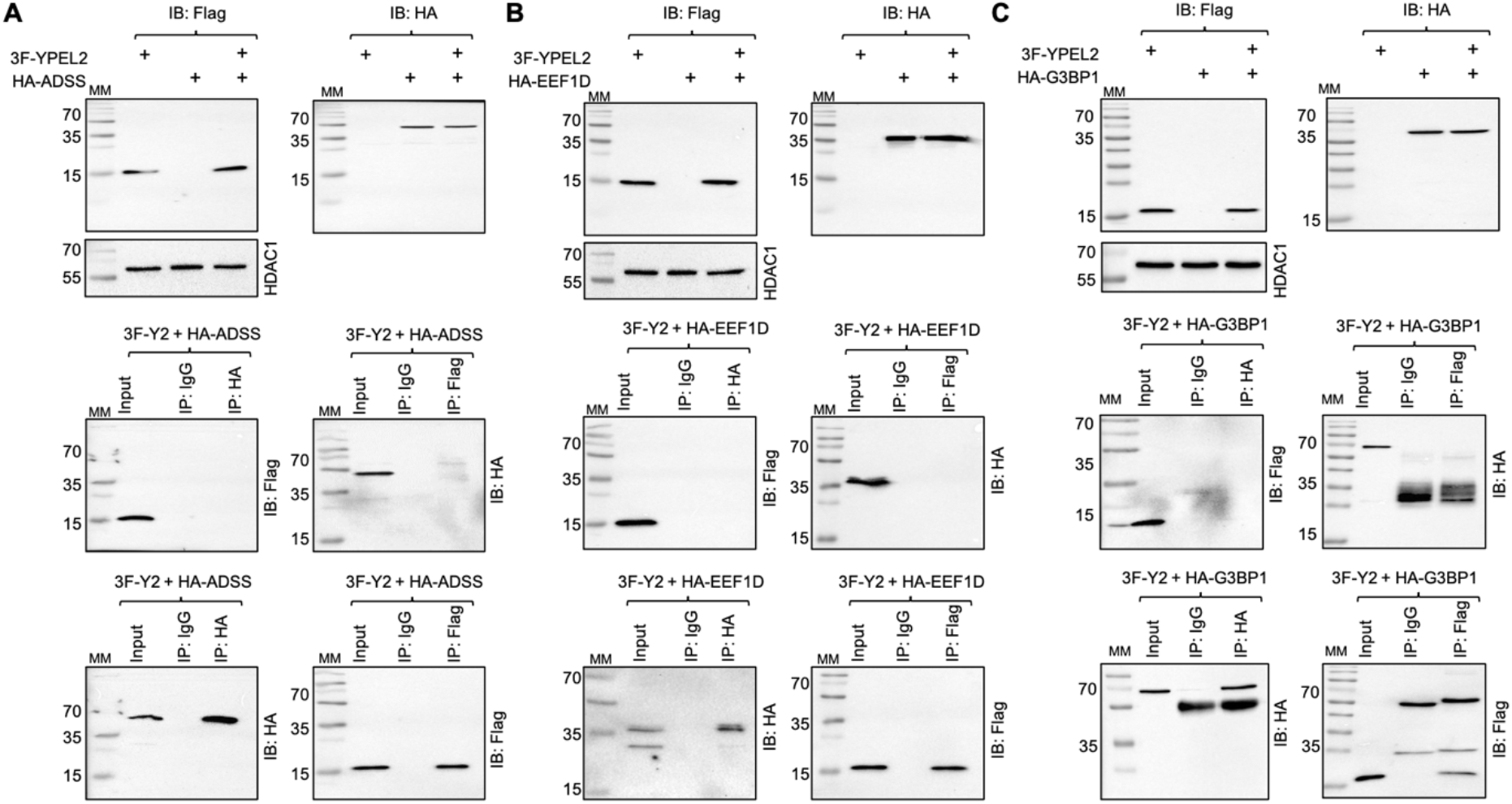
Interaction of 3F-YPEL2 with HA-ADSS, HA-EEF1D, or HA-G3BP1. (**A-C**) To assess the interactions of 3F-YPEL2 with putative interaction partners, the expression vector pcDNA3.1(-) bearing the 3F-YPEL2, (**A**) HA-ADSS, (**B**) HA-EEF1D, or (**C**) HA-G3BP1 cDNA were transiently transfected into COS7 cells for 24h. The synthesis of proteins was assessed by WB using the Flag (F-1804) or the HA (ab9119) antibody. HDAC1 used as a loading control was probed with the HDAC1 antibody (ab19845). Cellular extracts (500 μg) of transiently co-transfected cells were subjected to Co-IP with the Flag, HA, or isotype-matched IgG. The precipitates were then subjected to SDS-10%PAGE followed by WB using the HA or Flag antibody. Molecular masses (MM) in kDa are indicated.

**Fig. S11.**
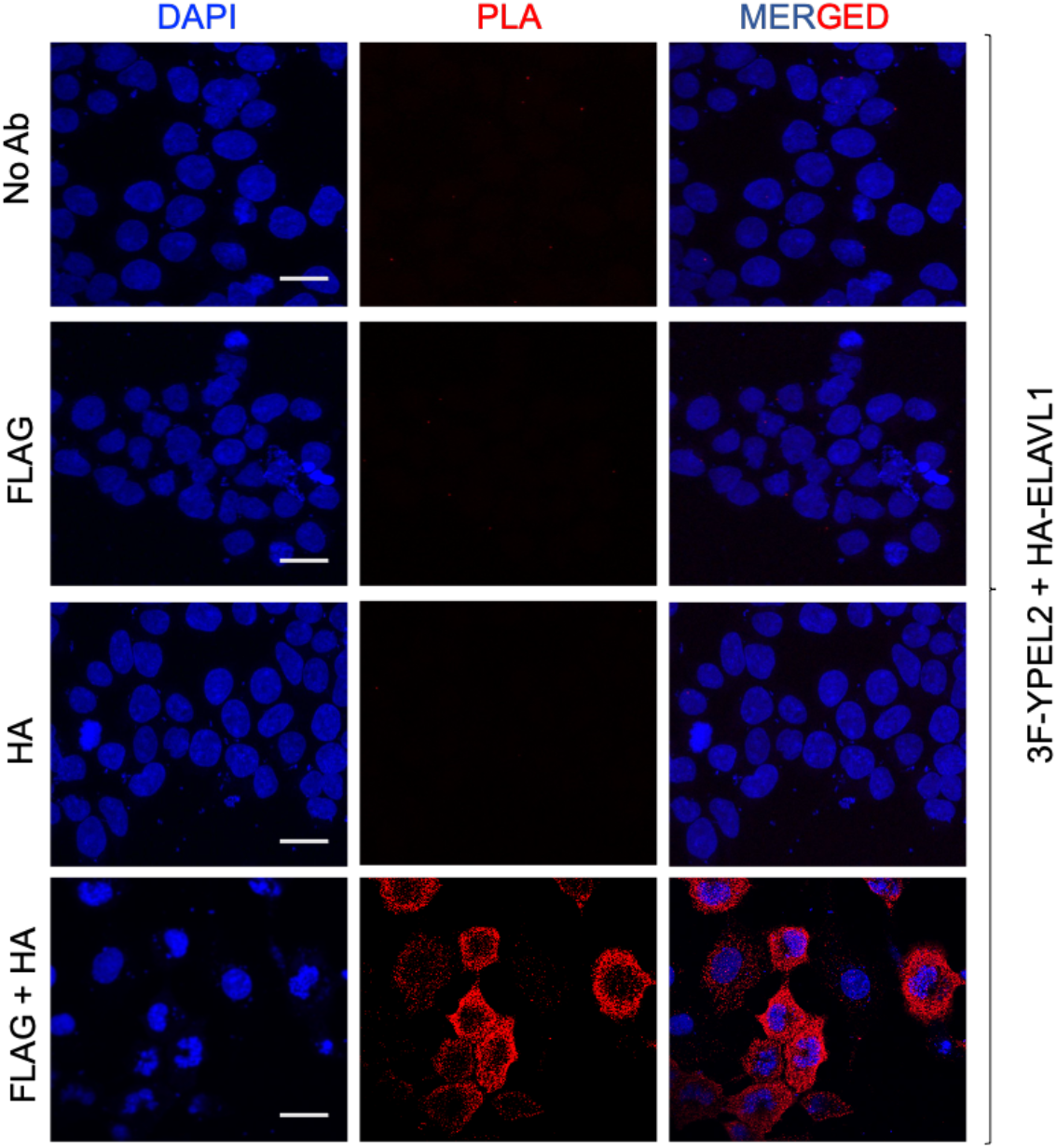
Proximity ligation assay for YPEL2 & ELAVL1 interactions. The interaction of 3F-YPEL2 and HA-SQSTM1 in transiently transfected COS7 cells was examined with proximity ligation assay (PLA). 36 h after transfections, cells were fixed, permeabilized, blocked, and probed without (No Ab) or with the Flag and/or the HA antibody. Cells were subsequently subjected to fluorescent probes for circular DNA amplification for PLA. DAPI staining indicates the nucleus. Images were captured with a fluorescence microscope. The scale bar is 20 µm.

**Fig. S12.**
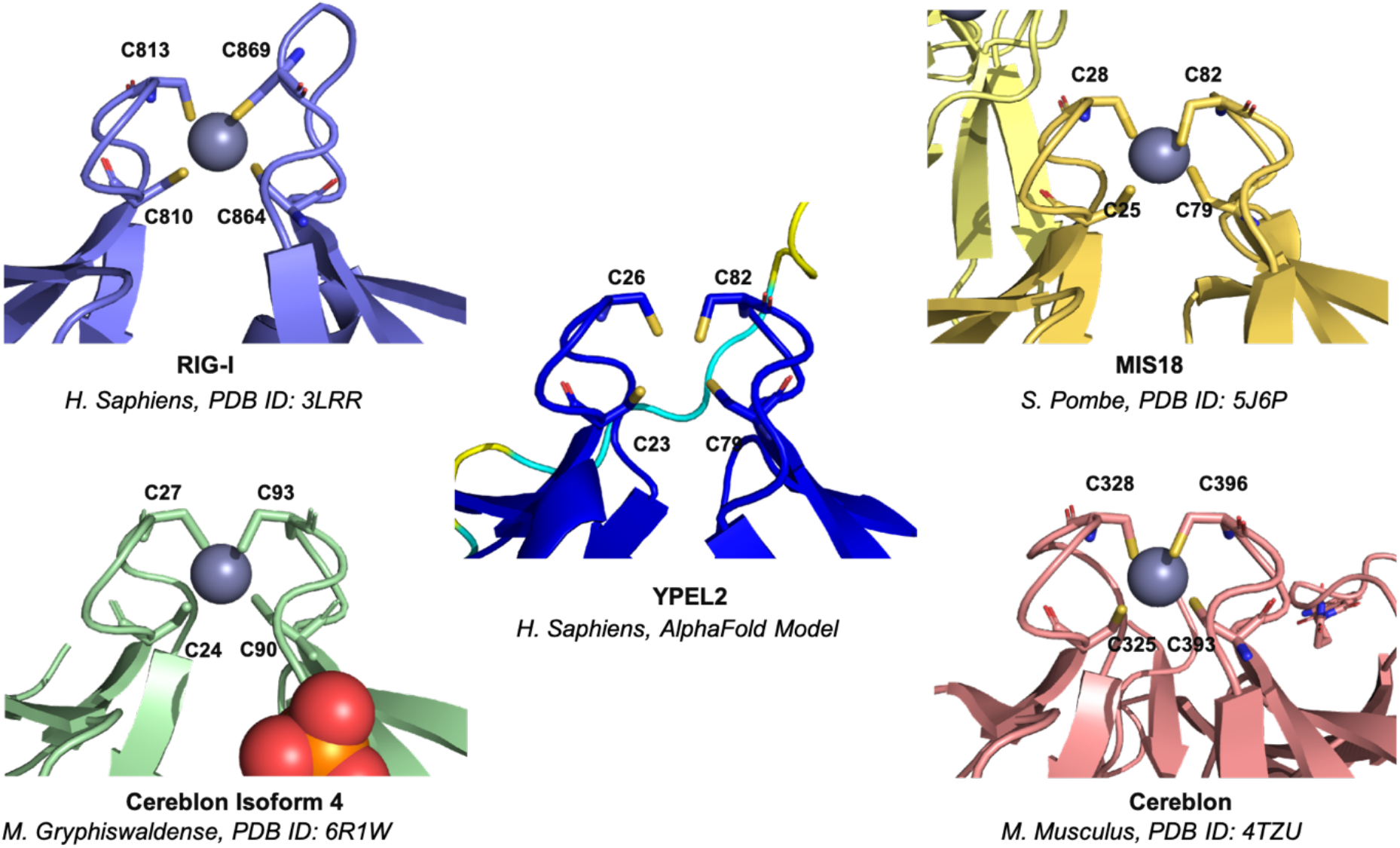
Zinc binding modeling of YPEL2. Structural similarity analysis reveals a conserved Zinc^++^ binding motif. The AlphaFold model of YPEL2 (https://alphafold.ebi.ac.uk/entry/Q96QA6) in the center is used as the basis for the structural similarity search. This analysis was performed using the PDBeFold server with a minimum acceptable match threshold set at 70%. As an outcome, we identified four distinct proteins, each originating from different organisms, all displaying a similar fold to YPEL2. Examination of these protein structures revealed a common Zinc^++^ binding motif mediated by four cysteine residues. Notably, while the AlphaFold model does not contain a Zinc ion, it accurately represents the spatial arrangement of cysteine residues observed in other structures. Zinc and cysteine residues are depicted in sphere and stick representations, respectively.

**Table S1.**
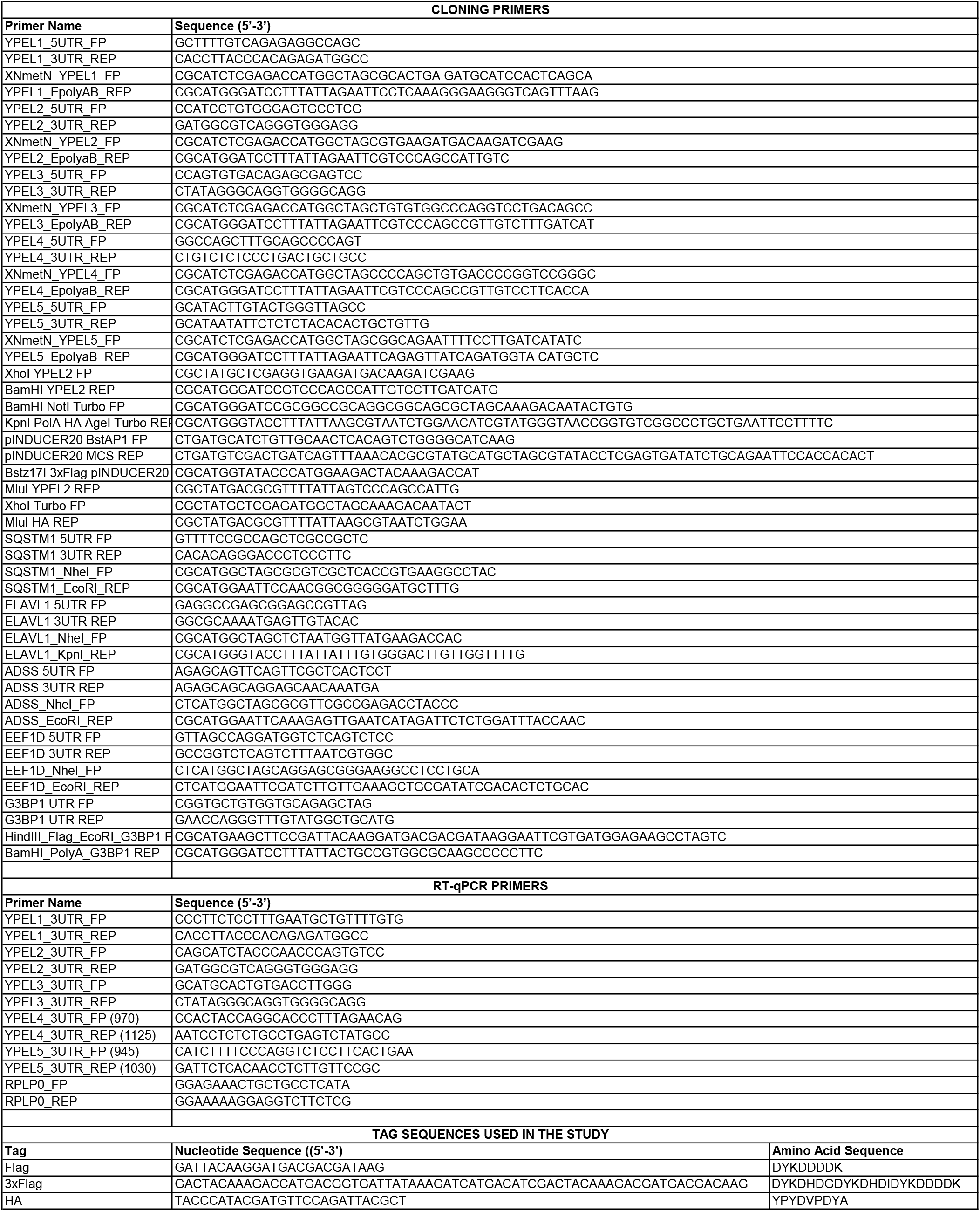

**Table S2.**
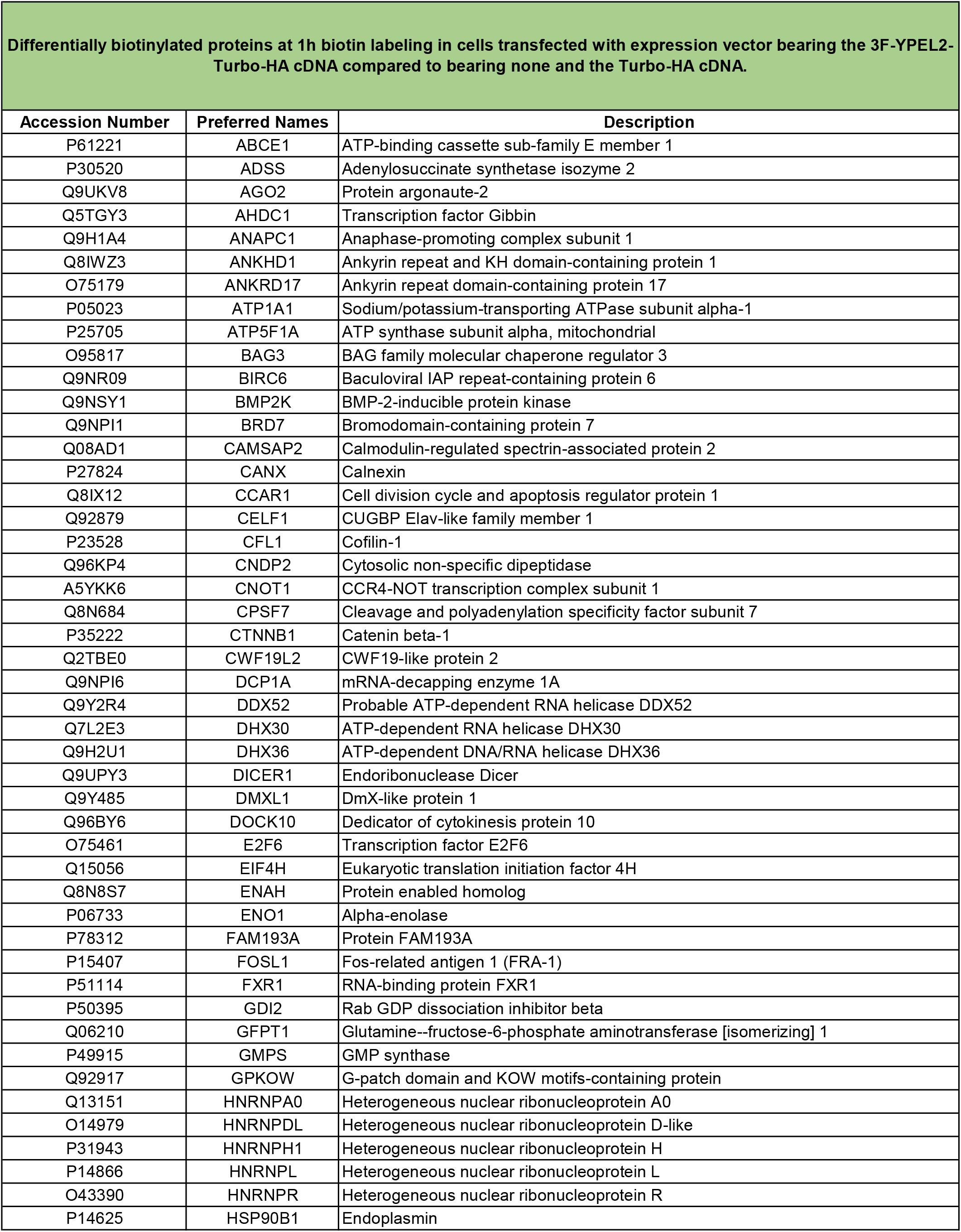
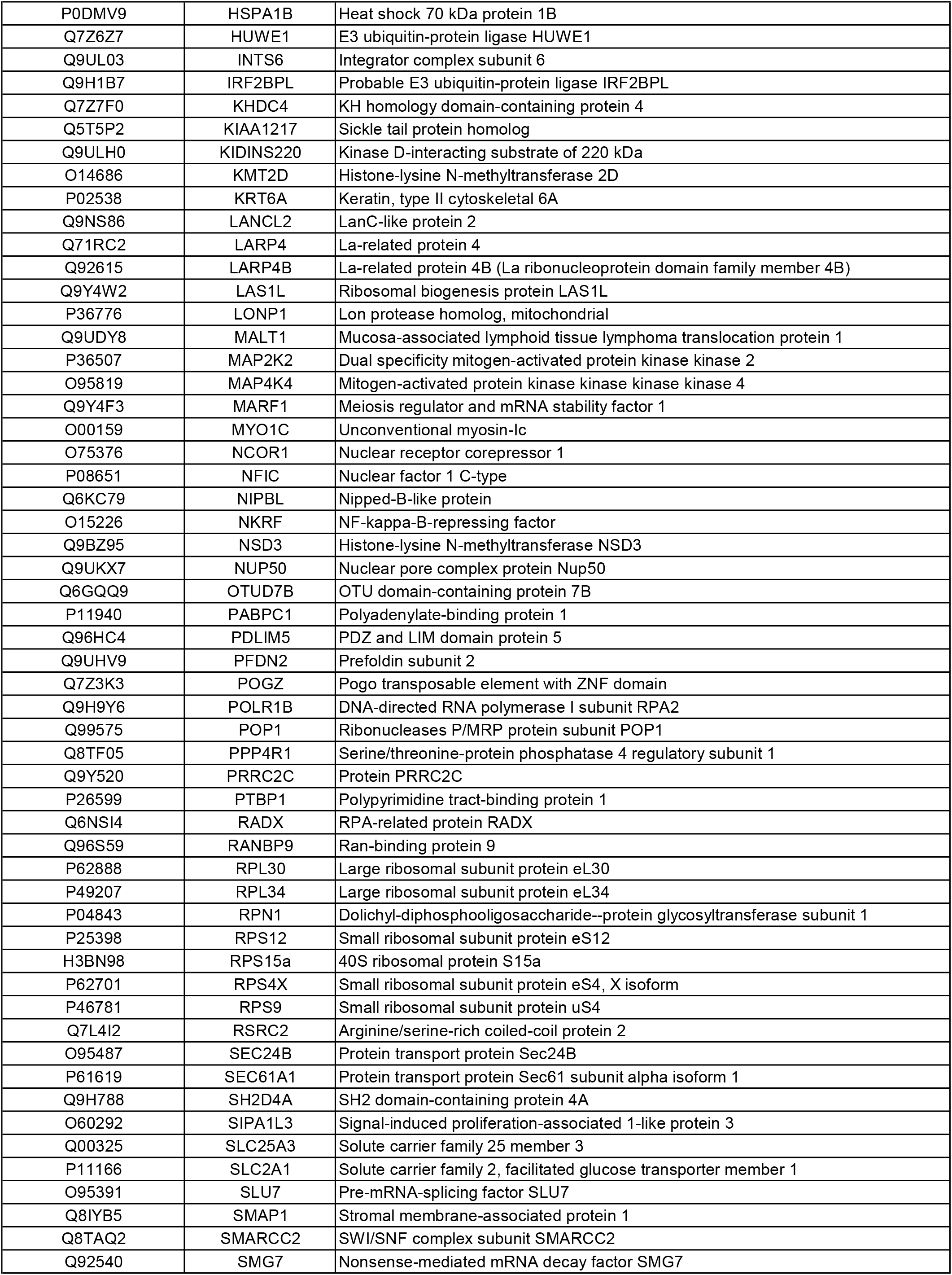
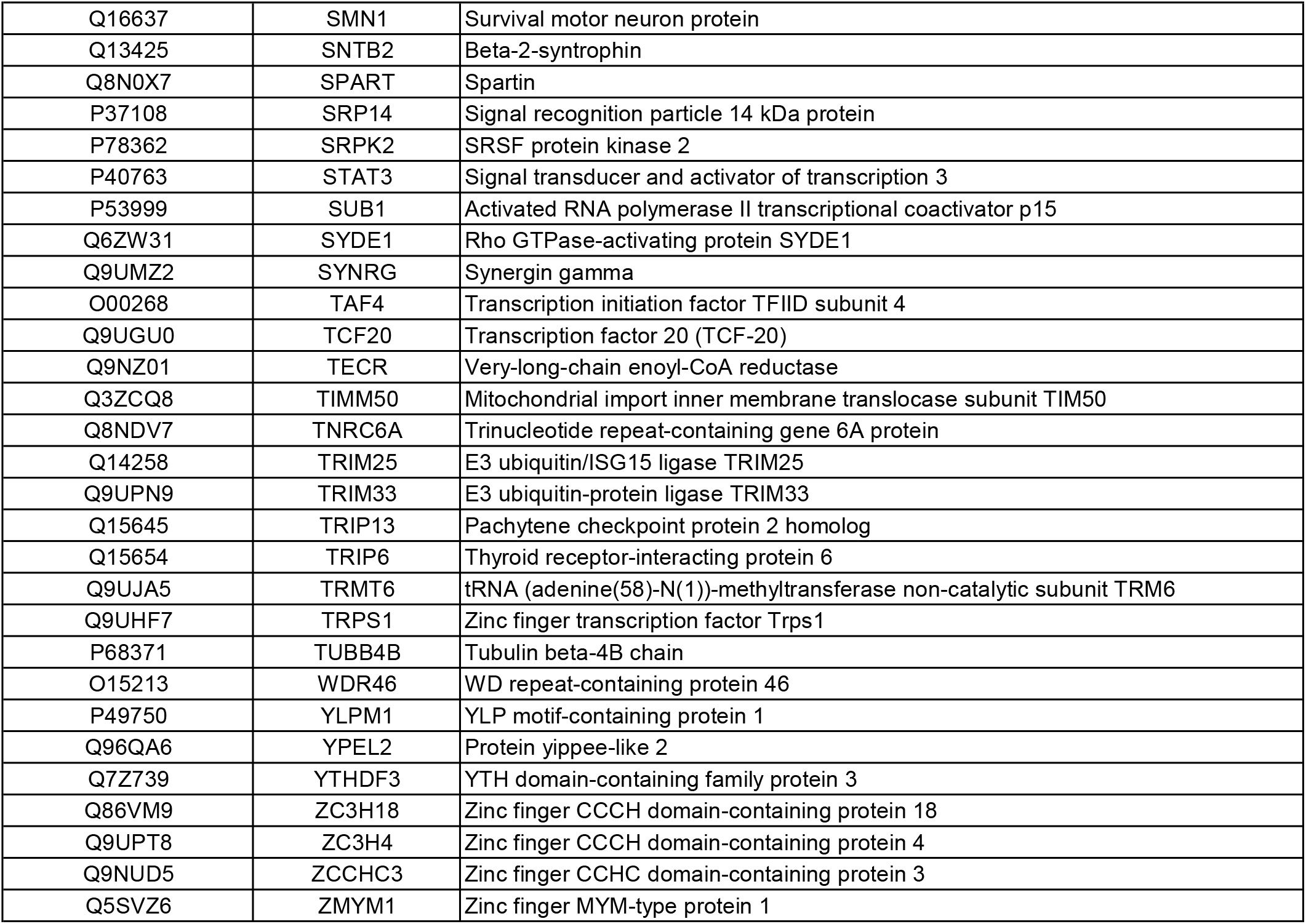
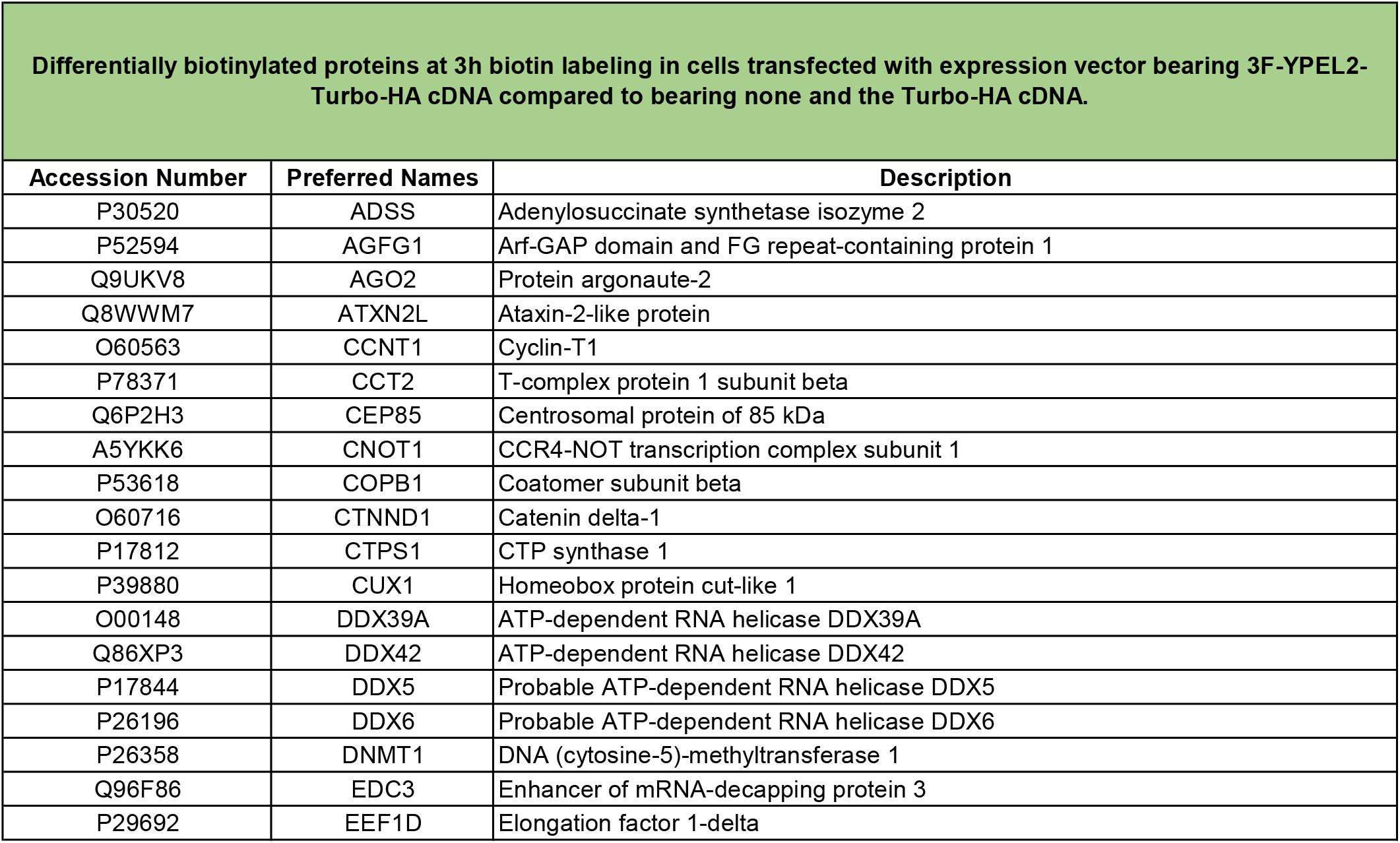
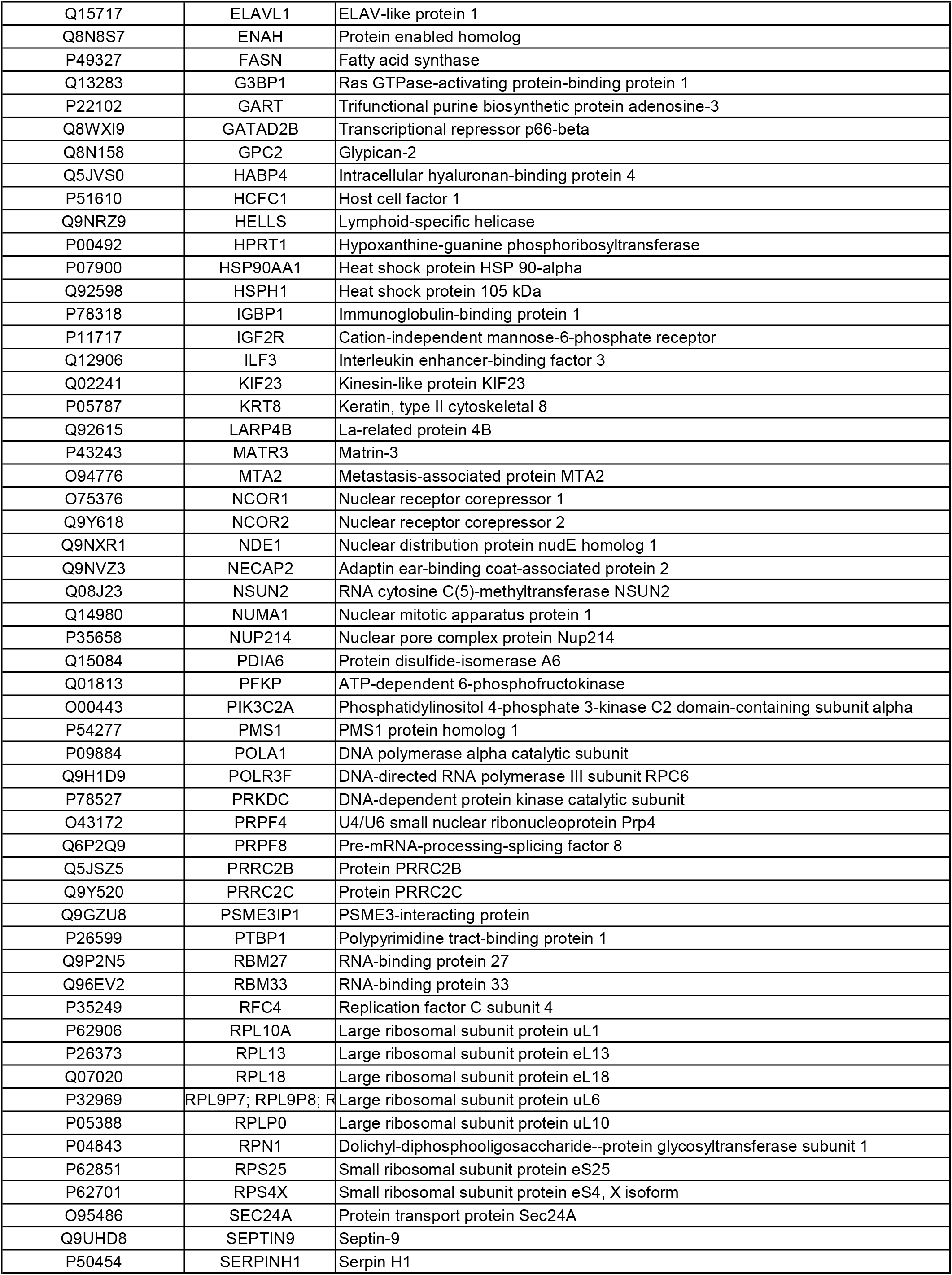
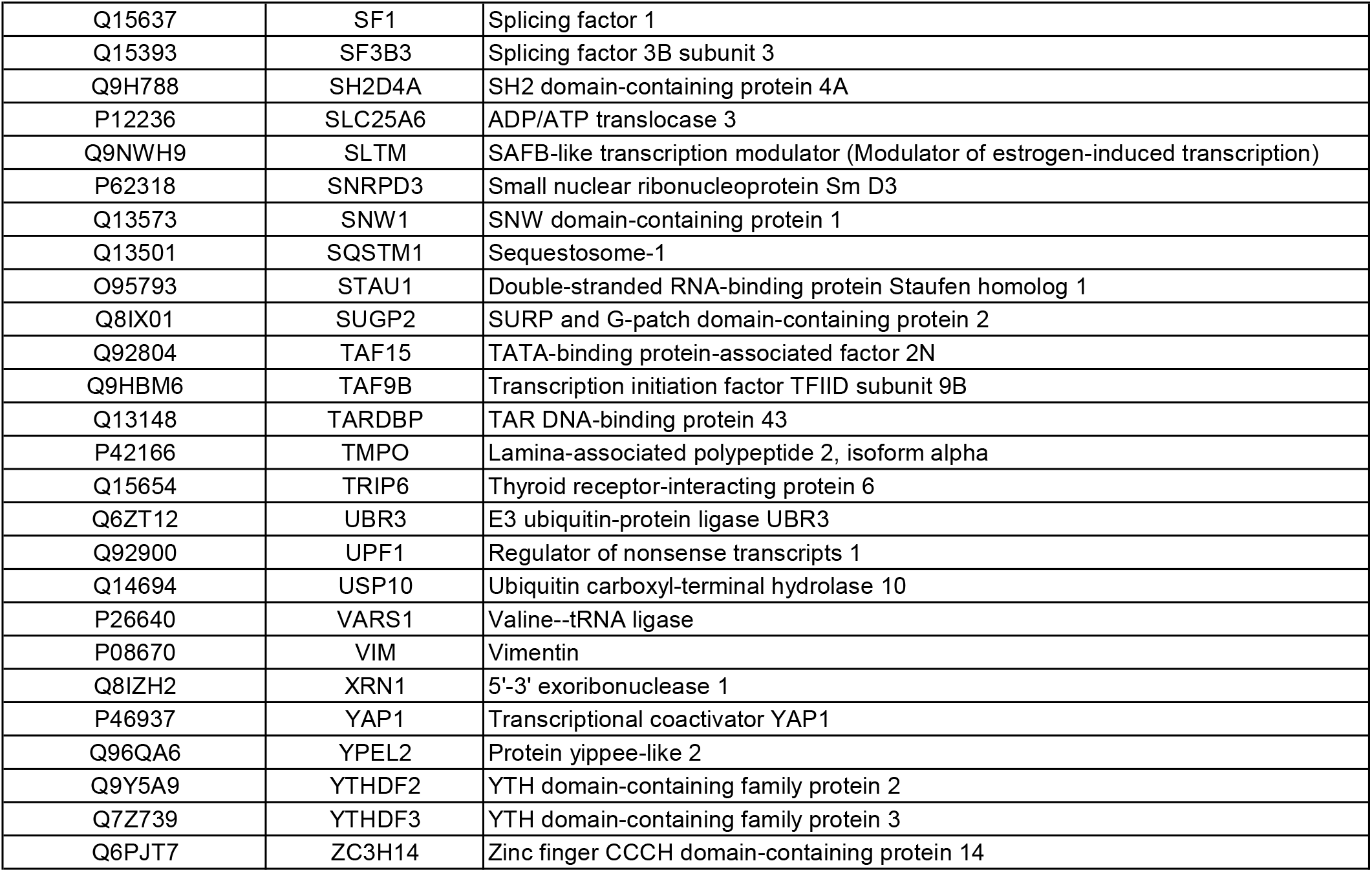
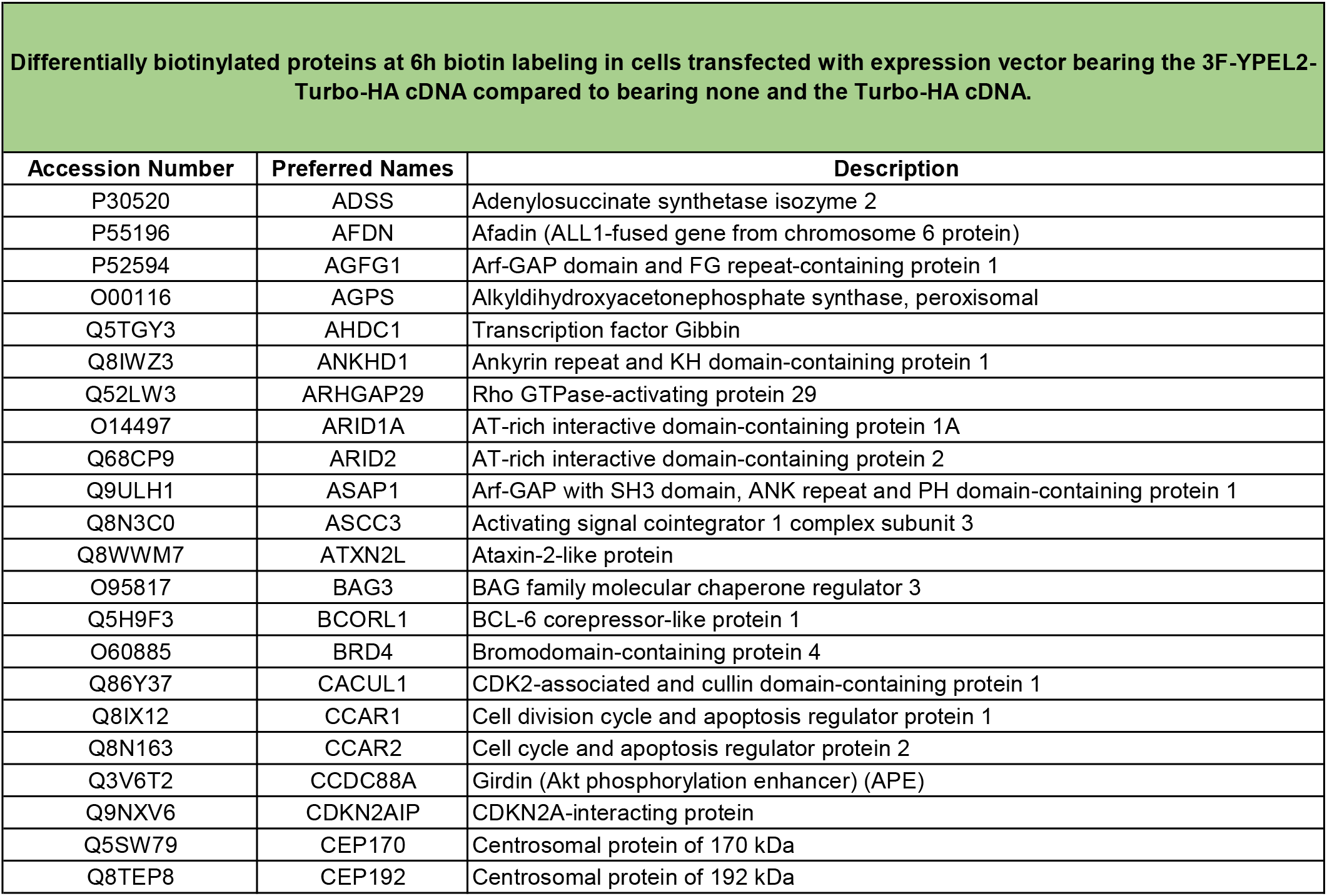
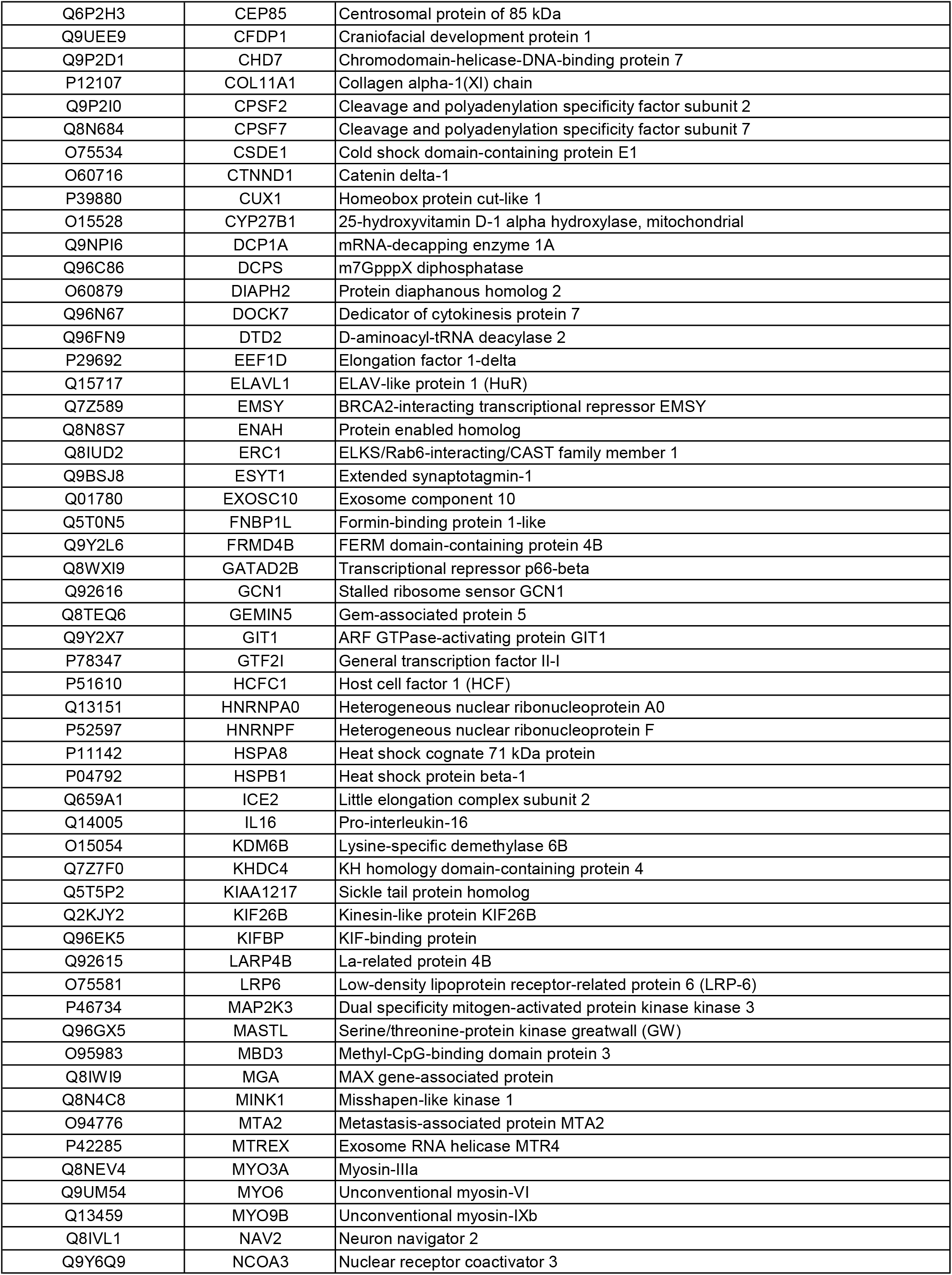
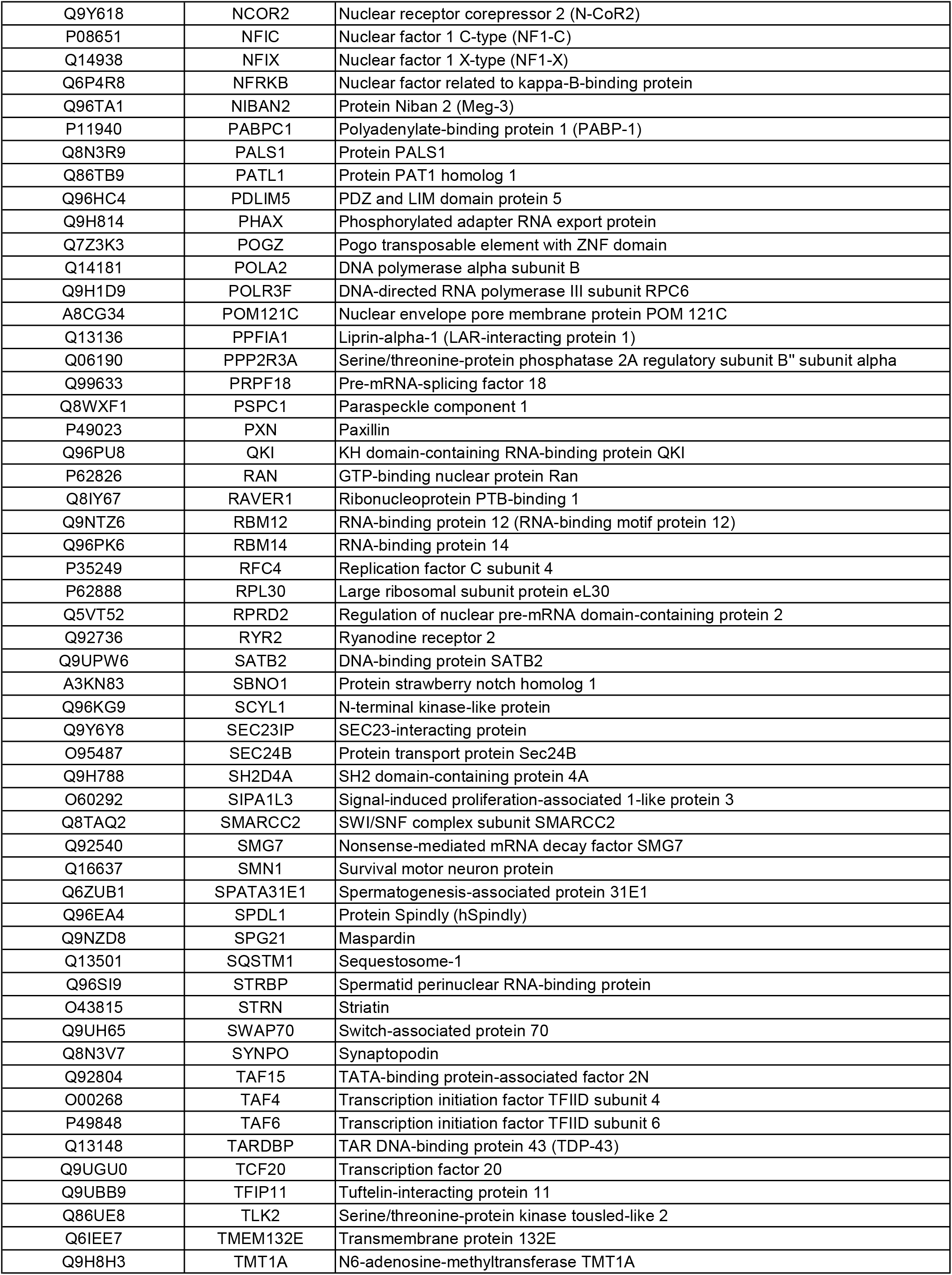
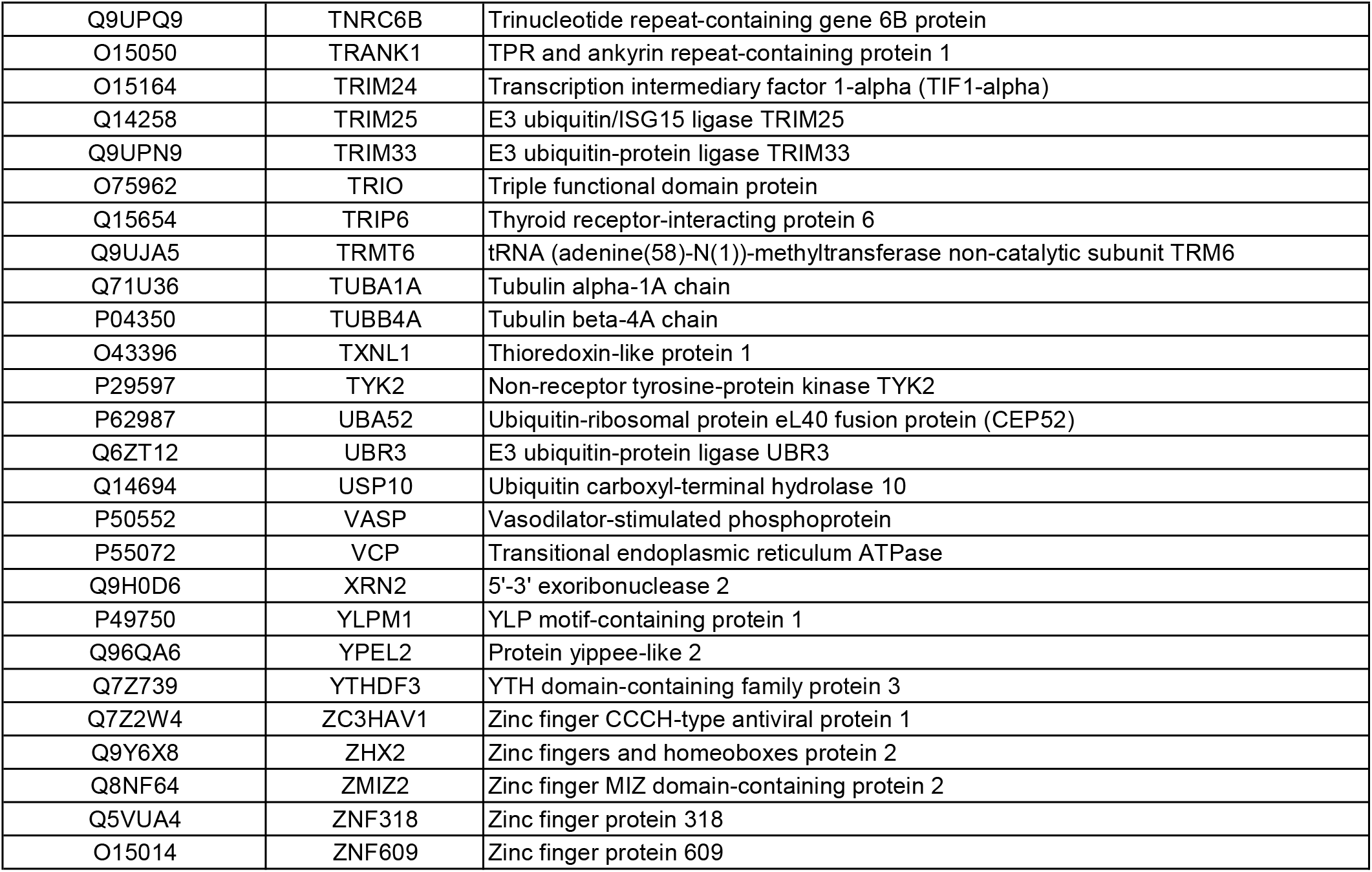
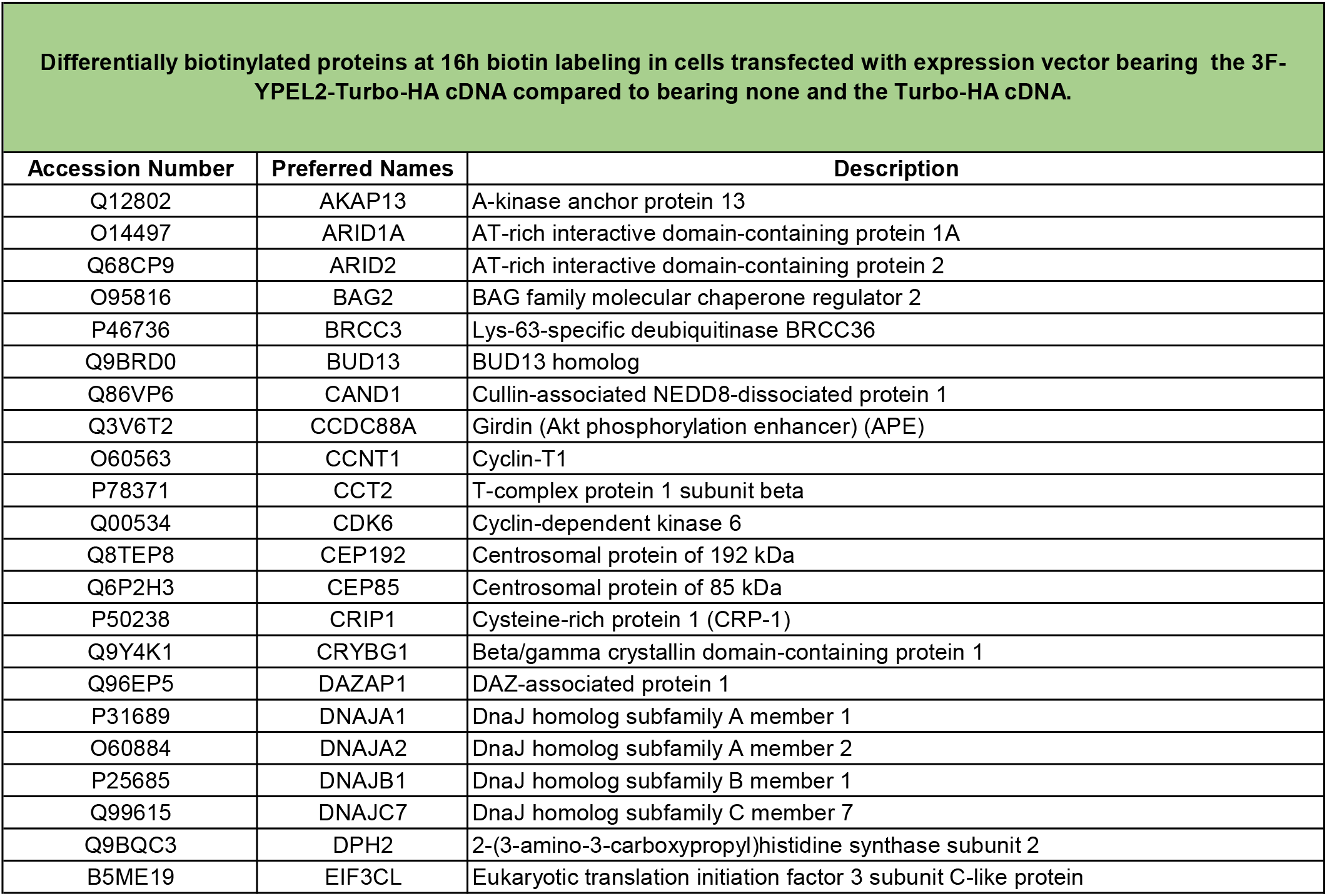
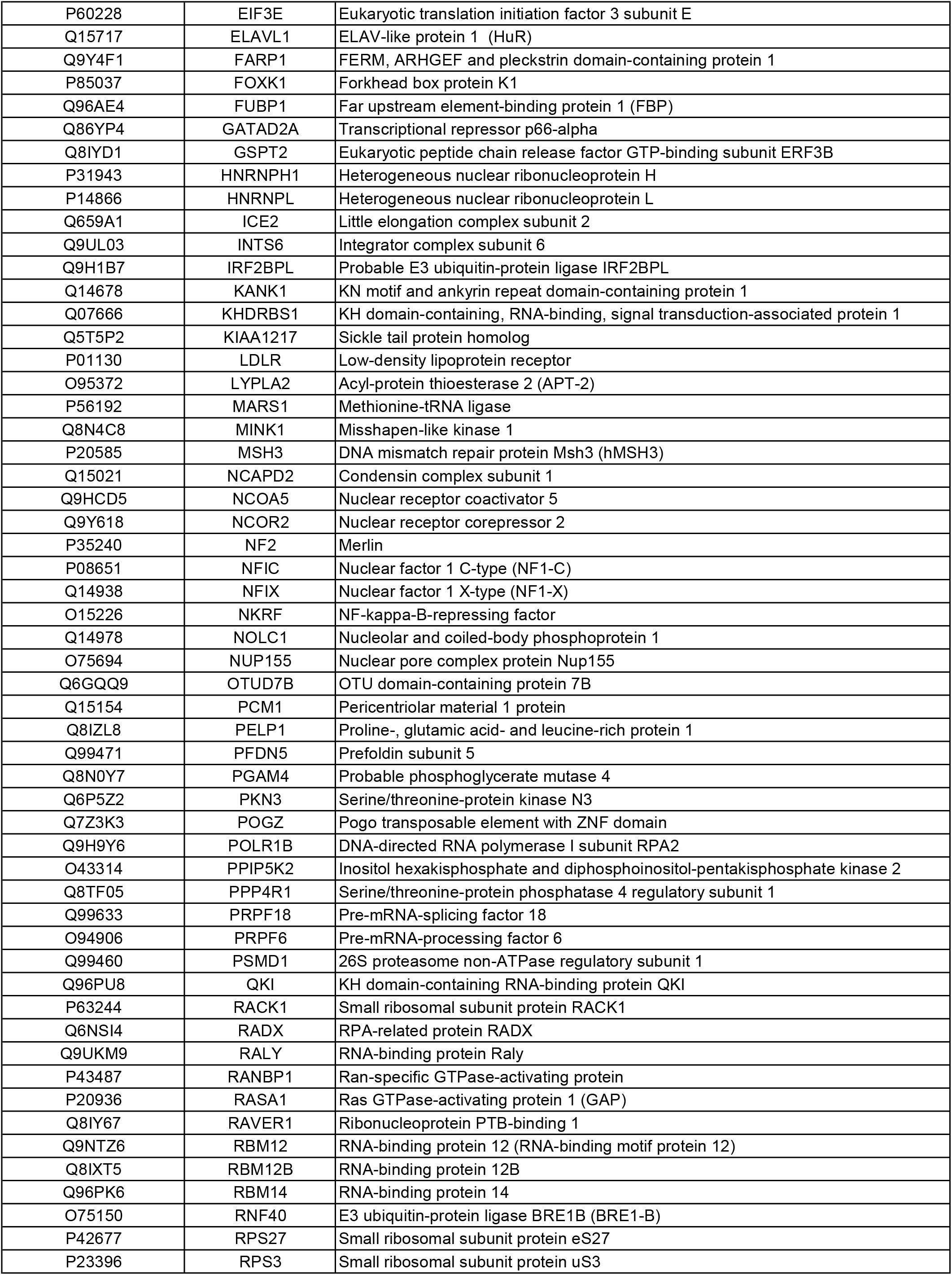
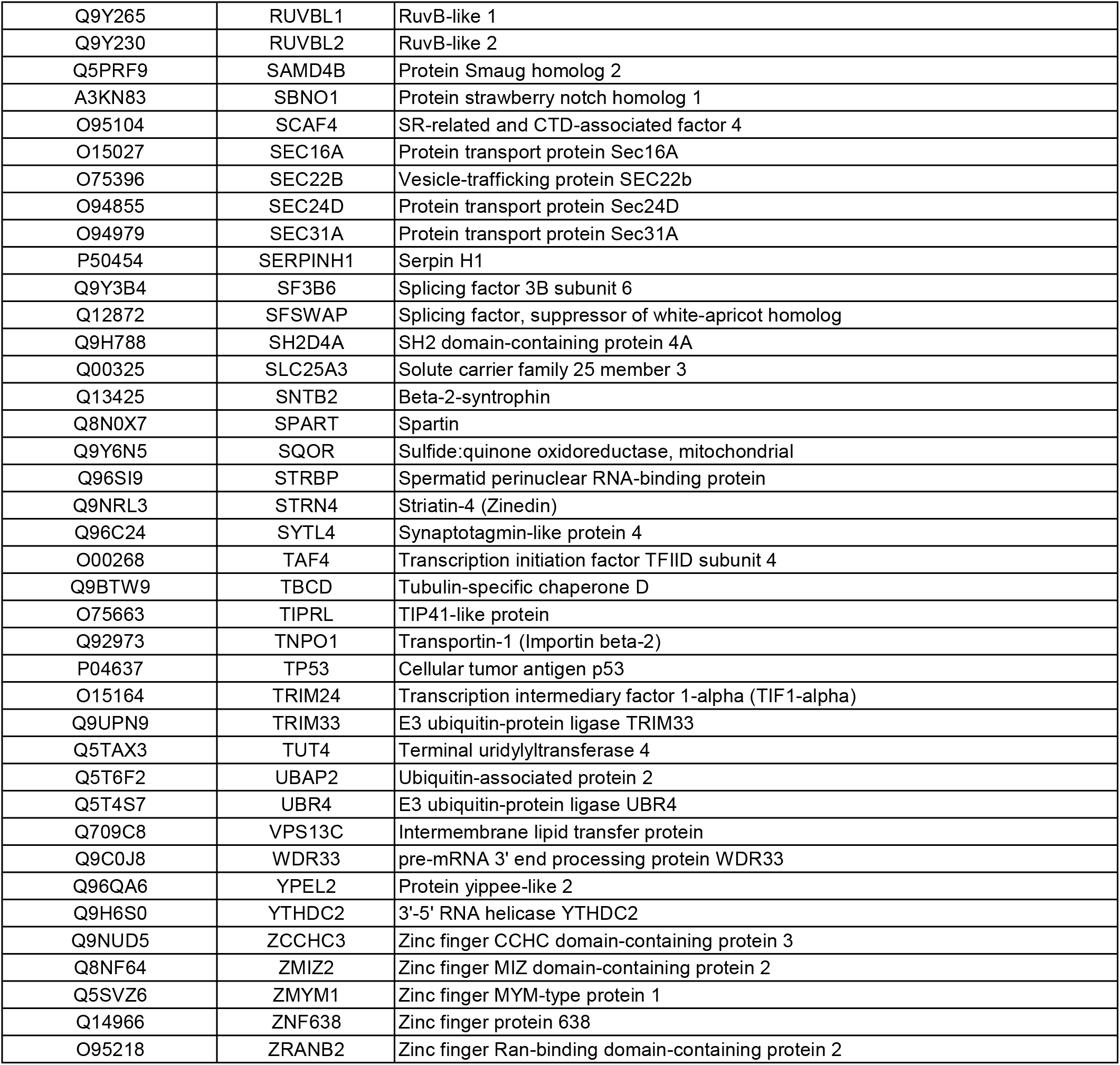

